# Cell cycle oscillations in a polarity network facilitate state switching by morphogenetic cues

**DOI:** 10.1101/2025.10.12.681824

**Authors:** KangBo Ng, Hadjar Sebaa, Nisha Hirani, Alex Chizh, Zeno Messi, Tom Bland, Kenji Sugioka, Nathan W. Goehring

## Abstract

The proper establishment of cell form, fate, and function during morphogenesis requires precise coordination between cell polarity and developmental cues. To achieve this, cells must establish polarity domains that are stable yet sensitive to guiding cues. Here we show that *C. elegans* germline blastomeres resolve this trade-off by creating a time-varying polarization landscape. Specifically, coupling the PAR polarity network to the cell-cycle kinase CDK-1 ensures that newborn cells operate in a low-feedback regime that lowers barriers to polarity state switching, allowing spatial cues to induce and orient PAR protein asymmetries. As CDK-1 activity rises at mitotic entry, increasing molecular feedback reinforces cue-induced asymmetries to yield robust and stable patterning of PAR domains. Consistent with this model, optogenetic and chemical perturbations show that low-CDK/low-feedback regimes destabilize PAR domains but are required for both *de novo* polarization and the reorientation of polarity in response to inductive cues. We propose that mitotic oscillations in cell polarity circuits dynamically optimize the polarization landscape to enable coordination of polarity with morphogenesis. Such temporal control of developmental networks is likely a general mechanism to balance robustness of cellular states with sensitivity to signal-induced state switching.

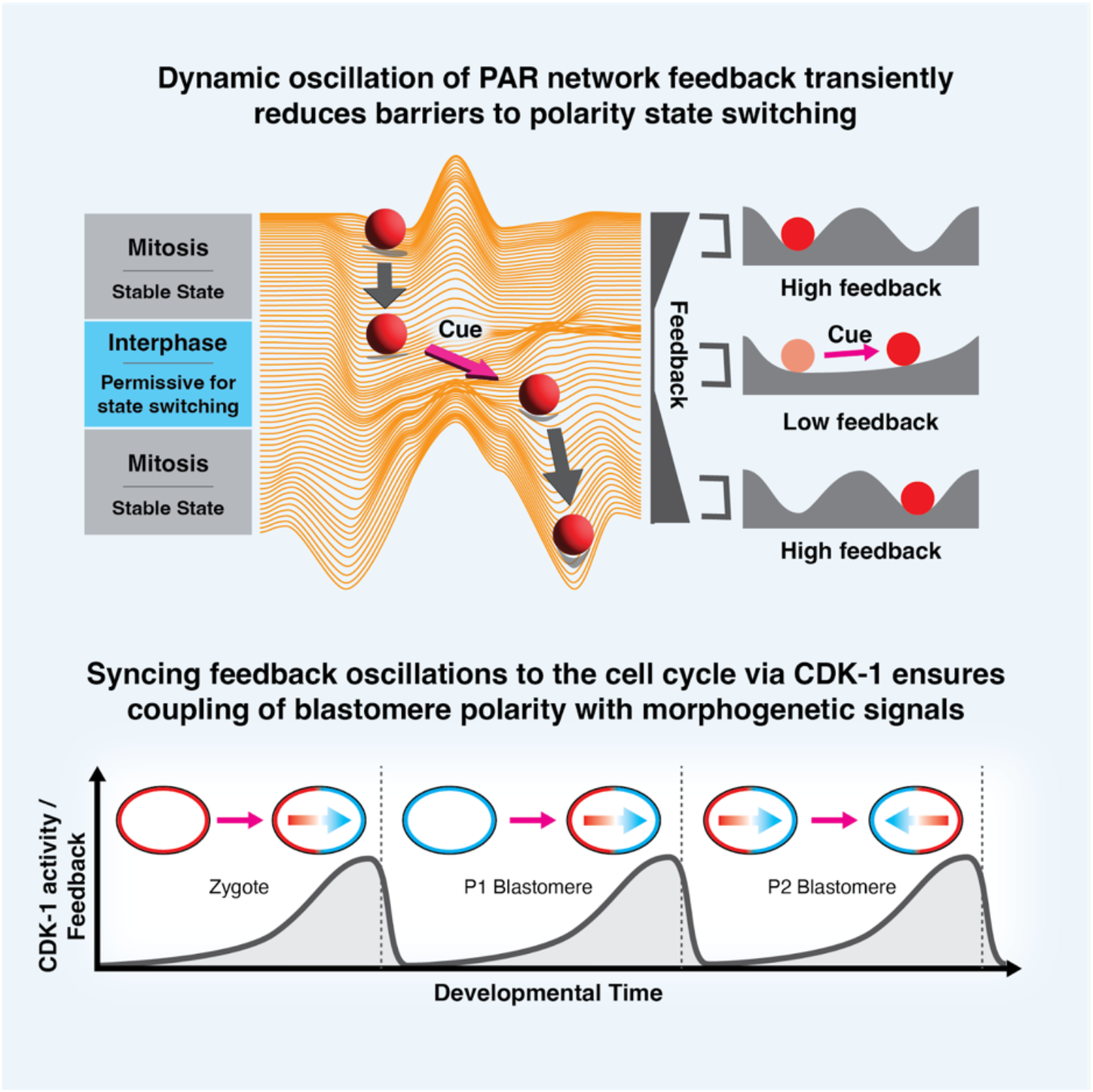

## Main Text

A key challenge for developmental signalling networks is to balance signal-sensitivity with output-stability. Robust signal responses are often attributed to feedback circuits that introduce bistable switch-like behavior or irreversibility into signaling networks ^1,2^. However, the same feedback that stabilizes outcomes against perturbations or stochastic variation can also reduce sensitivity, hindering the cells’ ability to respond to changes.

One context where this tradeoff is particularly evident is during cell polarization. Cell polarity describes the ability of cells to orient in space and typically involves self-organizing molecular networks which generate asymmetric protein patterns guided by spatiotemporal cues. One such network is the conserved PAR (*par*-titioning defective) network, which underlies a broad range of developmental processes in metazoans ^3–5^, including asymmetric cell division, cell migration, and organisation of tissue architecture. As the PAR network is continuously redeployed during development, it must remain sensitive to changing signals and cellular contexts so that polarity is properly oriented with respect to neighbouring cells or environmental cues. At the same time, once established, polarity must be sufficiently stable to reliably coordinate downstream processes in the face of perturbations. This dual requirement poses a paradox: cells that are too sensitive to signals will fail to maintain stable directionality, but introducing excessive stability into the network will hinder the ability of cells to respond and reorient with respect to spatial cues. Resolving this paradox is central to understanding how polarity is integrated into developmental programmes.

To address this question, we turned to the highly tractable *C. elegans* germline P lineage as a model. Beginning with the zygote (P0), P lineage blastomeres undergo a series of asymmetric divisions to generate the major founder lineages of the animal, including specification of the primordial germ cells ^6^. Importantly, these asymmetric divisions must be properly oriented by spatial cues so that the different cell types are correctly positioned within the embryo during development. This coordination requires that the polarity of P blastomeres adapt to dramatic shifts in cellular context, discriminate among competing cues, and respond to changing signals to polarize in the correct orientation ^3,6–12^; in other words, blastomeres must be highly sensitive to developmental cues, yet, once polarity is established it must be sufficiently stable to robustly inform downstream pathways, such as spindle position and segregation of fate determinants ^6^.

Despite blastomere-specific differences, the core principles of polarization are conserved. In each case polarization involves the cue-induced formation of two opposing, membrane-associated PAR domains, each harboring a distinct subset of PAR proteins termed anterior PARs (aPARs) ^13–17^ and posterior PARs (pPARs) ^18–21^, respectively. Mutual antagonistic feedback subsequently enforces segregation of these two PAR domains: the aPAR effector PKC-3 phosphorylates pPARs to exclude them from aPAR domains ^22,23^, whereas the pPAR effectors PAR-1 and CHIN-1 locally suppress aPARs within pPAR domains ^15,18,23–25^. Thus, comparative analysis of P blastomeres provides an ideal framework for dissecting how a single polarity network navigates the dual requirements for sensitivity and stability during development.

Here we reveal a central role for cell cycle-entrained oscillations in the PAR network in resolving this sensitivity-stability tradeoff. Examination of pPAR effectors reveals dynamic oscillation of their membrane localization in phase with CDK-1 activity. The resulting changes in network feedback shift blastomeres from a rheostat-like, low-feedback state early in the cell cycle, during which network behavior is highly responsive to instructive cues, to a high feedback, switch-like state that enforces stable patterns as the blastomere enters mitosis. We suggest that the pervasive mitotic oscillations in PAR protein behavior observed across metazoa ^26,27^ may reflect an ancestral design principle that allows cells to achieve both the signal-sensitivity and output-stability required for robust coordination of cell polarity with morphogenesis.

## Results

### CDK-1 drives oscillations in pPAR effector membrane association

The two key pPAR effectors, PAR-1 and CHIN-1, have been reported to exhibit changes in membrane concentration throughout the zygotic cell cycle ^15,20^. Quantifications of PAR-1 and CHIN-1 membrane levels with respect to a cell cycle marker (Histone H2B) revealed tight coupling to mitotic transitions: membrane concentrations were initially low, first increased following chromosome condensation, peaked around nuclear envelope breakdown (NEBD), and then declined at anaphase onset (Fig. **1b-d**, Supplementary Fig. **1**). This pattern of oscillation was repeated in the P1 blastomere, suggesting P lineage blastomere divisions are characterized by repeated cycles of membrane association and dissociation of pPAR effectors (Supplementary Fig. **1**).

**Figure 1.**
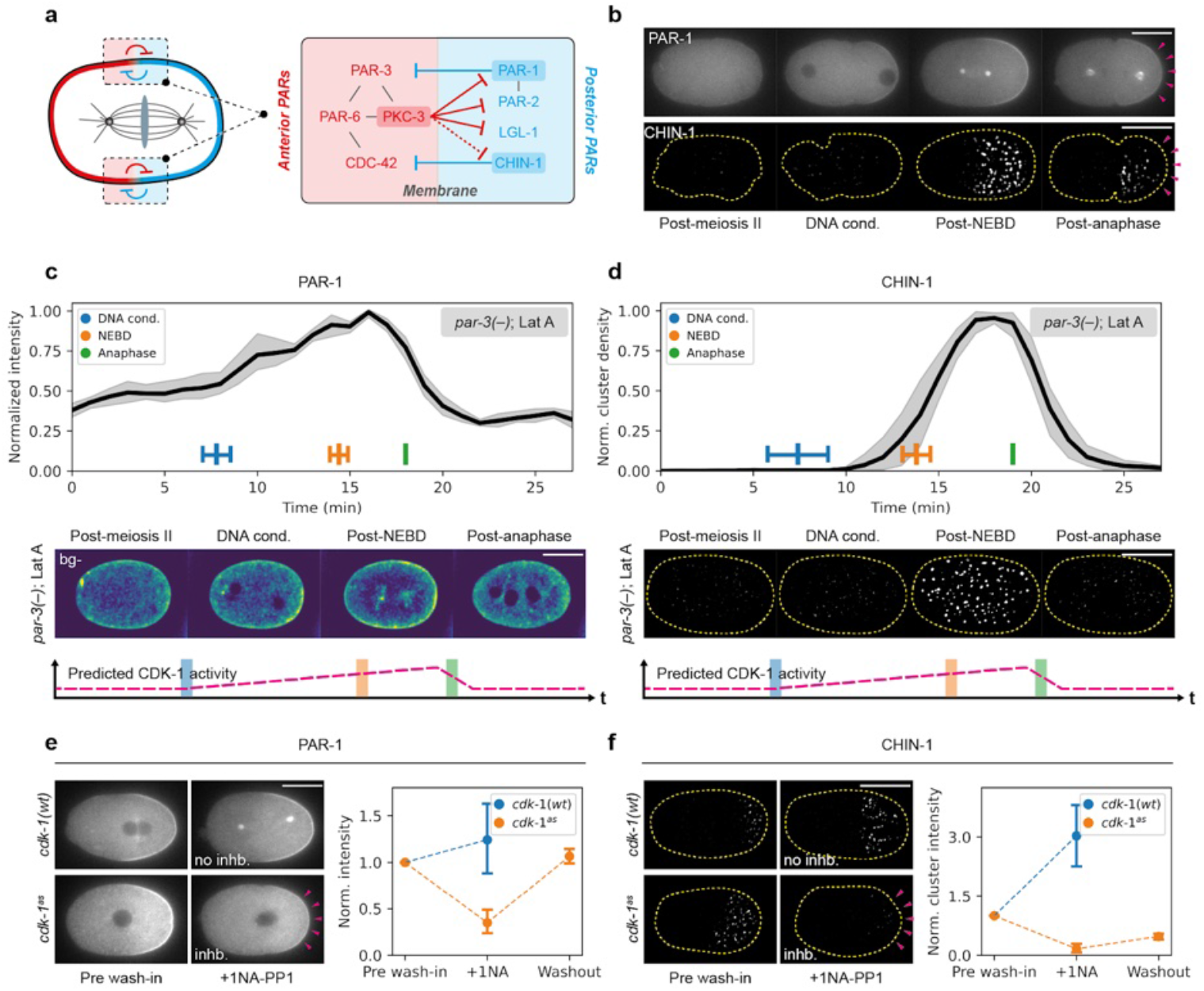
CDK-1 coupled oscillations in PAR behavior. **a**, A schematic illustrating mutual antagonistic feedback between aPARs (red) and pPARs (blue). For aPARs, the main effector is PKC-3, which locally removes all pPARs from the anterior membrane in the zygote. For pPARs, there are two effectors, PAR-1 and CHIN-1, which locally suppress PAR-3 and active CDC-42 from the posterior respectively. **b**, pPAR effectors exhibit oscillations in membrane levels over the cell cycle. Top, a time series of midplane confocal images of embryos expressing PAR-1::GFP with mCherry::PAR-2 (not shown) and H2B::mCherry in a *par-3(+/-)* background (NWG0434, roller phenotype) (n=4). Bottom, a time series of background subtracted (see Methods) cortical images of embryos expressing mNG::CHIN-1, H2B::mCherry and NMY-2::mKate2 (not shown) (NWG0528) (n=8). Magenta arrowheads indicate a sharp decrease in membrane levels following anaphase onset. Quantified (as in **c** and **d** in Supplementary Fig 1). **c**, Quantifications and images of embryos expressing PAR-1::GFP with mCherry::PAR-2 (not shown) and H2B::mCherry in a *par-3(-)* background (NWG0434, roller phenotype) treated with Latrunculin A throughout the cell cycle (n=5). Bottom shows the expected CDK-1 activity during the different cell cycle stages, which appears positively correlated with PAR-1 membrane levels. Times are normalized relative to anaphase. **d**, Same as **c**, but for embryos expressing mNG::CHIN-1, H2B::mCherry, NMY-2::mKate in a *par-3(-)* background (NWG0543) treated with Latrunculin A (n=5). Images in both **c** and **d** are background subtracted. **e**, PAR-1 membrane levels respond to changes in CDK-1 activity. Left, midsection confocal images of embryos expressing PAR-1::GFP and mCherry::PAR-2 (not shown) in either a *cdk-1(wt)* (NWG0042) (n=5) or *cdk-1*^*as*^ background (NWG0520) (n=7), acutely treated with 20μM 1NA-PP1 after pronuclear meeting (PNM). Note the reduction in PAR-1 membrane levels after CDK-1 inhibition (magenta arrowheads). Middle, the corresponding quantification for conditions shown on the left, showing also that PAR-1 membrane levels increase when CDK-1 inhibition is relieved. **f**, CHIN-1 membrane levels respond to changes in CDK-1 activity. Same as **e** but for cortical images of embryos expressing mNG::CHIN-1 in either a *cdk-1(wt)* (NWG0451) (n=4) or *cdk-1*^*as*^ background (NWG0518) (n=4). Mean and 95% confidence interval (bootstrapped) indicated. Scale bars, 20μm.

We next sought to determine whether the oscillatory membrane association of pPAR effectors reflects direct coupling to the cell cycle, or regulation by other cell cycle-dependent processes, such as changes in the activity or localization of aPAR proteins, remodeling of the cortical actin cytoskeleton, centrosome growth cycles, or changes in the activity of key PAR regulators LKBP/PAR-4 or 14-3-3/PAR-5, could all contribute to membrane oscillations of pPAR effectors ^15,17,23,25,28–31^. However, consistent with intrinsic cell-cycle-dependent regulation, we found that oscillations persisted in *par-3(-)* embryos, in *par-3(-)* embryos treated with latrunculin A, and in embryos subject to RNAi targeting other PAR proteins (*par-2, par-4, and par-5*) or the core centrosome component *spd-5* (Fig. **1b-d**, Supplementary Figs. **1-3**).

The pattern of pPAR effector oscillations - low early in the cell cycle, increasing rapidly after mitotic entry, and declining at anaphase onset - roughly mirrored the expected activity of the mitotic kinase CDK-1 ^32–34^. To explicitly test whether pPAR membrane concentrations were controlled by CDK-1, we generated an analog-sensitive *cdk-1* (*cdk-1*^*as*^) allele ^35–38^, allowing specific and acute inhibition of CDK-1 activity (Supplementary Fig. **4**). Acute inhibition of CDK-1^AS^ with the ATP analogue 1NA-PP1 reduced PAR-1 and CHIN-1 membrane signals, which were partially restored upon drug washout (Fig. **1e,f**, Supplementary Videos **1,2**). Finally, we confirmed that changes in PAR-1 and CHIN-1 membrane levels were not due to changes in aPAR activity (Supplementary Fig. **4**), strongly suggesting that pPAR effector oscillations are entrained by CDK-1 activity.

### CDK-1 drives oscillations between low and high feedback states

To assess whether these CDK-1-dependent changes in pPAR effector membrane association correlated with changes in the ability of pPARs to antagonize aPARs ^23,25,39–41^, i.e. pPAR to aPAR (P→A) feedback, we examined the response of the aPAR protein PAR-6 to CDK-1 inhibition. Normally, PAR-6 is excluded by PAR-1 and CHIN-1 from the pPAR domain following polarization ^23^. However, upon inhibition of CDK-1, PAR-6 invaded the posterior domain as marked by PAR-2, leading to overlap of the two proteins at the posterior (Fig. **2a,b**, Supplementary Fig. **5a**). This suggests that CDK-1 inhibition reduces P→A feedback. This phenotype did not appear to stem from off-target effects on the cytoskeleton or regulators downstream of CDK-1 such as AIR-1 and PLK-1, which have also been implicated in PAR polarization (Fig. **2c,d**, Supplementary Fig. **5**, Supplementary Video **3**) ^42–47^.

**Figure 2.**
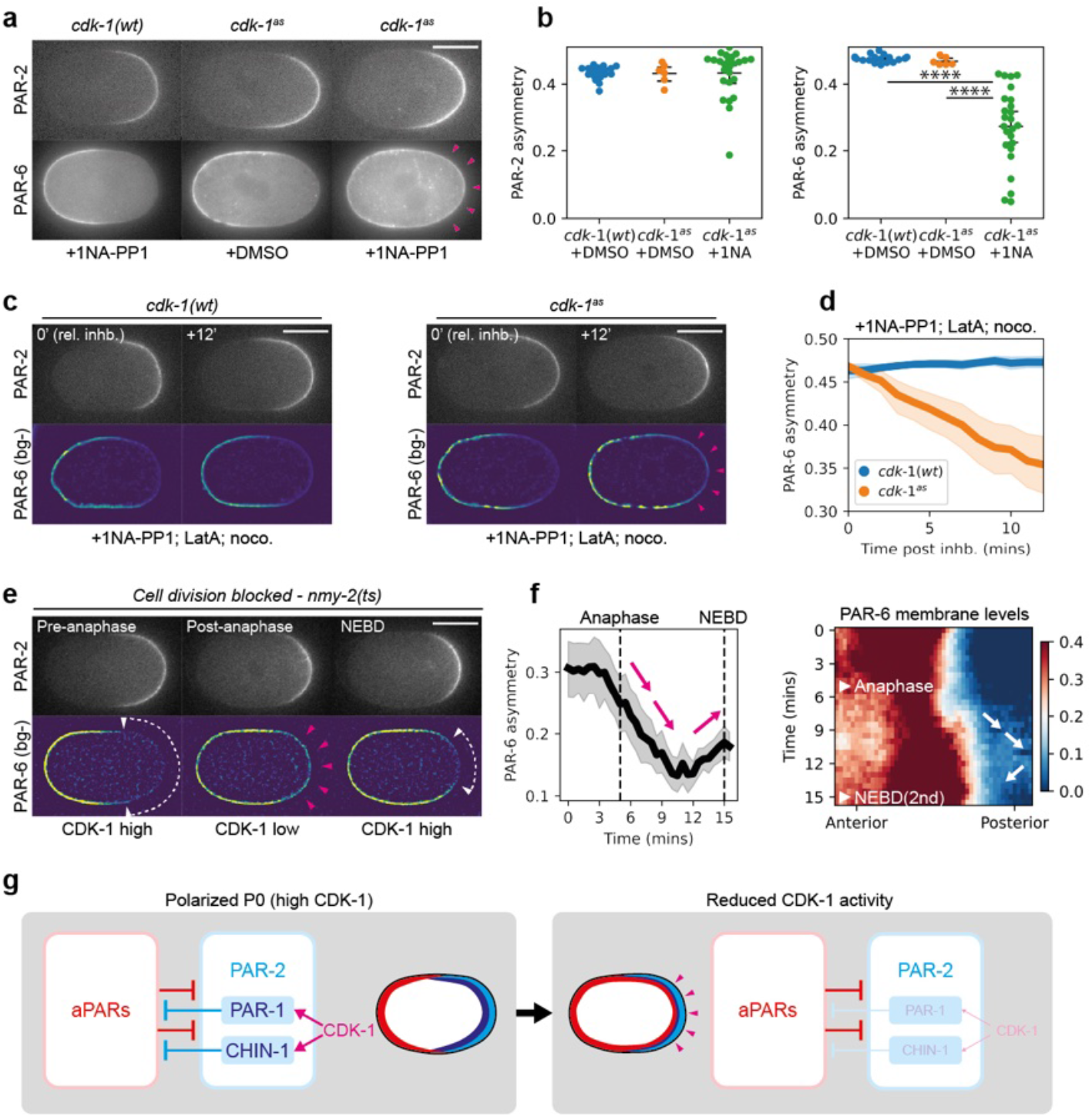
CDK-1 couples PAR feedback strength to the cell cycle. **a**, CDK-1 inhibition leads to reduction in PAR-6 asymmetry. Midsection confocal images of embryos expressing mCherry::PAR-2 and PAR-6::mNG in *cdk-1(wt)* (NWG0268) or *cdk-1*^*as*^ (NWG0443) background, treated with either DMSO or 50μM 1NA-PP1. Sample sizes: *cdk-1(wt)* + 1NA-PP1 (n=18), *cdk-1*^*as*^ + DMSO (n=8), *cdk-1*^*as*^ + 1NA-PP1 (n=25). Open magenta arrowheads indicate the presence of PAR-6 in PAR-2 occupied domains. **b**, Quantification of asymmetry for embryos corresponding to conditions shown in **a**. Each point represents data from a single embryo. **c**, PAR-6 invasion phenotype in CDK-1 inhibited embryos are not dependent on cytoskeleton. A time series of midplane confocal images, before and after combination treatment with 1NA-PP1, Nocodazole and Latrunculin A, for embryos expressing mCherry::PAR-2 and PAR-6::mNG in either a *cdk-1(wt)* (NWG0268) (n=6) or *cdk-1*^*as*^ (NWG0443) (n=7) background. Magenta arrowheads indicate PAR-6 invasion into posterior PAR-2 domains. PAR-6 images were background subtracted (bg-) to improve visibility due to poor signal-to-noise ratios. **d**, Quantification of PAR-6 asymmetry over time for conditions corresponding to **c. e**, PAR-6 invasion into PAR-2 domains is correlated with cell cycle stage. A time series of midsection confocal images of an embryo expressing mCherry::PAR-2 and PAR-6::mNG in an *nmy-2(ts)* background (NWG0509) (n=8). Acute temperature upshift to the restrictive temperature was performed to inhibit *nmy-2(ts)* activity after zygote polarization, decoupling cell division from cell cycle progression. White arrowheads with dotted lines indicate PAR-6 exclusion from PAR-2 domains. Open magenta arrowheads indicate the presence of PAR-6 in PAR-2 occupied domains. **f**, Quantification of PAR-6 asymmetry and membrane distribution over time for conditions corresponding to **e**. Magenta and white arrows both indicate loss of PAR-6 asymmetry following anaphase and increase in asymmetry before NEBD, corresponding to a presumptive loss and gain of CDK-1 activity respectively. **g**, Schematic showing that CDK-1 positively regulates P→A feedback via controlling PAR-1 and CHIN-1 membrane levels, leading to changes in the ability for PAR-2 domains to exclude aPARs. Mean and 95% confidence interval (bootstrapped) indicated. Scale bars, 20μm. Student’s t-test was performed, unpaired, two-tailed. Scale bars, 20μm. n.s. = not significant, *, p-value < 0.05, **, p-value < 0.005, *** p-value < 0.0005, **** p-value <0.00005.

We next looked at the response of PAR-6 to oscillations in CDK-1 activity through the cell cycle. In wild-type zygotes, we observed posterior spreading of PAR-6 following anaphase onset, when CDK-1 activity and P→A feedback is expected to be low (Supplementary Fig. **6a,b**). However, we are unable to accurately quantify the full extent of the spreading, as formation of the cleavage furrow is able to alter the positioning of PAR proteins ^11,48^. To circumvent this, we inhibited cytokinesis via acute inactivation of NMY-2 using a temperature sensitive *nmy-2(ts)* allele ^49^, or by depleting the formin CYK-1 ^50,51^. In both cases, PAR-6 transiently invaded the posterior PAR-2 domain following anaphase onset and was then cleared from the posterior as the embryos re-entered mitosis (∼NEBD) of the second cell cycle (Fig. **2e,f**, Supplementary Fig. **6c**).

Consistent with this result, we observed cycles of aPAR-pPAR overlap and clearance in later P blastomeres P1-P3 as previously described ^11,12^. Specifically, PAR-6 accumulated steadily throughout the membrane early in the cell cycle, despite these cells being born with uniformly high membrane levels of PAR-2 (Supplementary Fig. **6d,e**). PAR-6 only cleared from PAR-2 domains as the cells entered mitosis, resulting in mutually exclusive polarity domains.

Detecting reduced pPAR to aPAR feedback early in the zygotic (P0) cell cycle is complicated by the fact that pPARs are initially absent from the membrane following meiosis II due to the high initial levels of aPARs at the plasma membrane ^46^. However, by partially depleting PAR-6, we were able to generate zygotes in which pPARs were uniformly enriched on the plasma membrane (Supplementary Fig. **6f**). In these zygotes, the residual PAR-6 on the plasma membrane initially overlapped with PAR-2 early in the cell cycle, but was rapidly cleared at mitotic entry (NEBD). Altogether, our data indicates that P→A feedback oscillates in phase with CDK-1 activity, shifting from low feedback early in the cell cycle to high feedback late (Fig. **2g,h**).

### Oscillatory feedback allows robust response to polarizing cues from diverse initial states

We next investigated the potential impact of this oscillation between low and high feedback regimes by turning to a simplified two component reaction-diffusion model for polarity (see Methods) ^8,9,53^. As described previously, this system relies on interconversion between membrane-associated and cytoplasmic states, limiting pools, mass conservation and mutually antagonistic feedback. Previous work has shown that for high, balanced levels of feedback, the system is multistable, capable of supporting both uniform and polarized states ^8,52^.

To comprehensively capture system behavior during polarization, we constructed a landscape of different polarity states, described by the concentration difference between aPARs (A) and pPARs (P) in the anterior (A_a_-P_a_) and in the posterior (A_p_-P_p_) (Fig. **3**) ^54^. Consistent with the multistable behavior of the system, for any given initial state the system tends to evolve towards one of four steady-states that defines the four quadrants in the landscape: (i) uniform A high, (ii) uniform P high, (iii) PA polarized - P high in anterior, A high in posterior, (iv) AP polarized - A high in anterior, P high in posterior (Fig. **3a**, Supplementary Fig. **7**). Thus, high feedback creates well-defined wells or attractors, with polarization effectively represented by a transition between alternative polarity states. In the case of the zygote, polarization would reflect a transition between states (i) and (iv). Note that while we do not observe other potential transitions in the zygote (i.e. polarization from a uniform P high state (ii to iv) or polarity reversal (iii to iv)), such transitions are observed in later P blastomeres ^7,11,12^ (see below).

**Figure 3.**
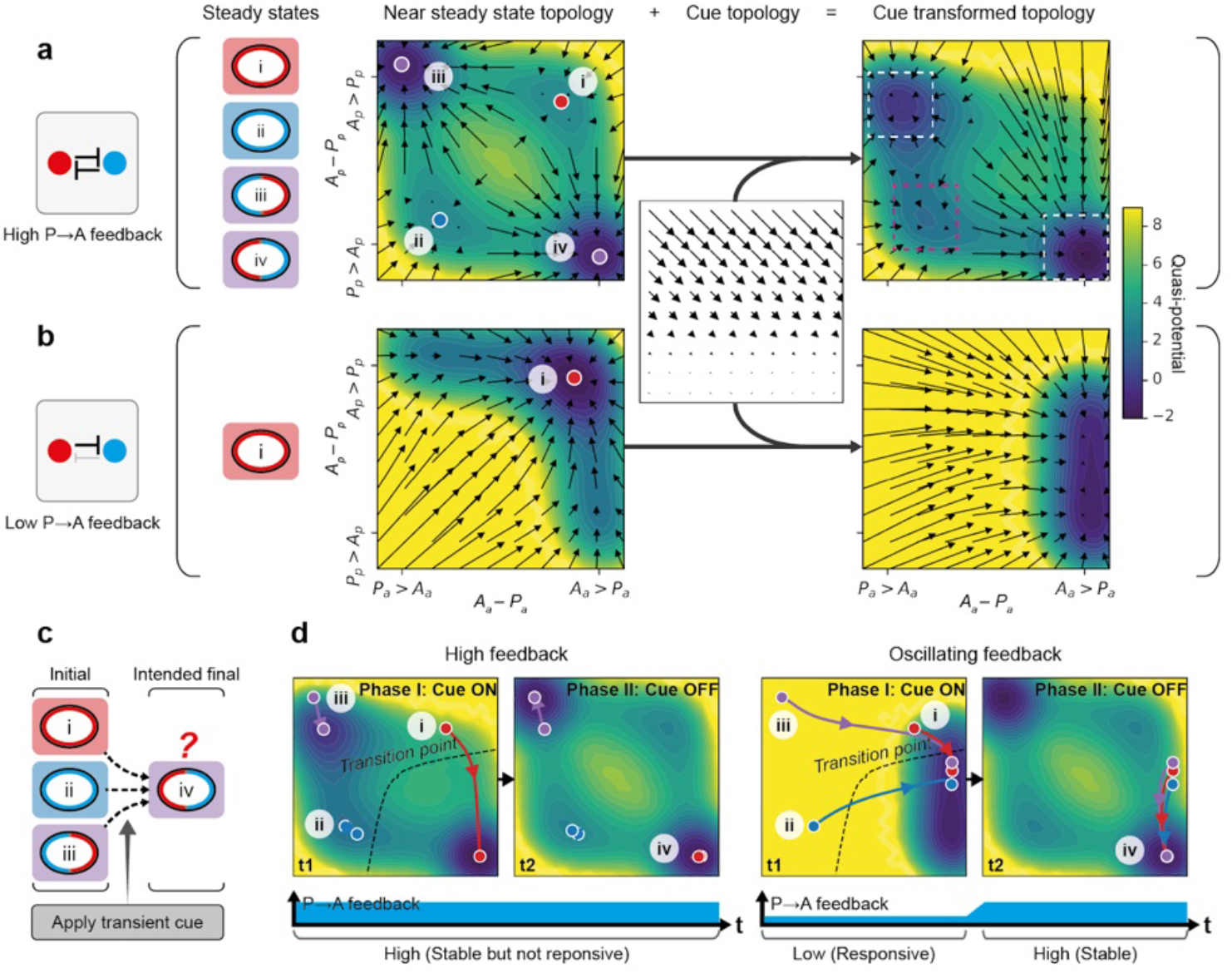
Oscillating feedback facilitates robust and cue-responsive polarization from diverse initial states. **a**, High feedback produces stable attractors for polarity states but is unable to respond to cues from a range of initial states. A balanced high feedback between aPARs (A) and pPARs (P) facilitates 4 possible steady states. An unstable steady state at the center of the diagram is not shown ^8,52^. State i - uniform A high, ii - uniform P high, iii - polarized with pPARs high at the anterior and aPARs high at the posterior, iv - polarized with aPARs high at anterior and pPARs high at posterior. These 4 steady states can be mapped onto a concentration-difference landscape, defined by the difference between A and P at the anterior (A_a_ - P_a_) and the difference between A and P at the posterior (A_p_ - P_p_). Quivers represent the directionality and velocity of movement towards each of these steady states for every system state in the landscape. System behavior under cue induction can also be visualised by transforming the landscape with the cue topology, which is represented as a fractional redistribution of aPARs from the anterior to posterior (A_p_ to A_a_) continually, inducing polarization towards state iv. In the case of high P→A feedback, there is a region near state ii and iii in the landscape that can not respond well to cue induction. Quasi-potential is defined as the -ln(probability) that a stochastic simulation, started from a random initial condition near the system’s steady state, arrives at a given position in the phase plane. We performed 10^7^ independent simulations to estimate this quantity (see Modelling Supplement). As the full system is four-dimensional (A_a_, A_p_, P_a_, P_p_), but we project and visualize the results in two-dimensions (A_a_ - P_a_, A_p_ - P_p_), the resulting “potential-like” surface averages over behaviours that differ along the hidden dimensions, which could lead to mismatch with the quivers. Nonetheless, the quasi-potential still conveys the relative stability of each state in the landscape, in the presence of noise. To confirm that these behaviors are similarly present in a non-dimensional-reduced model, we repeated the simulations in a single species polarity model, which is fully tractable; we observed that feedback can similarly act against the cue topology to prevent accurate polarization (Supplementary Fig. **9**). Dotted white boxes indicate steady state attractors of the system, whereas the magenta box indicates transient trapping of the system in a shallow well. **b**, Low feedback does not support stable polarized states, but promotes cue-responsiveness. Here, there is only one stable state, which is state i - uniform A high. However, in presence of a cue, all points across the landscape are correctly redirected towards the state iv quadrant, unlike in the case of high feedback. **c**, Schematic illustrating the full simulation of PAR polarization related to **d**, intending to polarize systems from state i, ii and iii to state iv using the same cue described above. **d**, Oscillatory feedback allows stable yet adaptable polarization from all points in the landscape. Left, in the constant high feedback condition, only points around state i are able to respond reliably to cues and move beyond the transition point for polarization towards state iv. Right, in the oscillatory feedback condition, temporary low feedback facilitates cue-responsiveness towards the transition point for all initial states, before increasing feedback locks in the stable polarized state. Importantly, all these properties are similarly captured in a full partial differential equation model, validating our approach (Supplementary Fig. **10**).

In early *C. elegans* embryos, cues are thought to operate via local depletion or inhibition of aPAR activity ^8,11,12,22,25,55–57^. To represent these cues generically, we implemented a continuous process of fractional redistribution of A from posterior to anterior. When transformed with this cue, we found that the uniform A high state (i) vanishes and initial states in this quadrant evolve towards the AP polarized state (iv). Thus, consistent with prior analysis, this high feedback regime effectively captures polarization of the zygote. By contrast, initial uniform P (ii) and reversed PA polarity states (iii) were more resistant to the cue, with the system remaining trapped near the initial state. Thus, for fixed high feedback and an aPAR-acting cue, proper cue-oriented polarization is only observed for systems beginning in an initially uniform A state.

We next considered a low feedback regime in which P→A feedback is reduced (Fig. **3b**). For sufficiently low P→A feedback, attractors states reflecting uniform P high (ii) and polarized states (iii, iv) vanish, leaving only a single uniform A high attractor (Supplementary Fig. **7**). Thus, no stable polarized states are possible. Transforming this low feedback landscape with the cue shifted the single attractor towards the AP polarized quadrant. Thus, in this regime, all initial states converge on a single state exhibiting properly oriented AP polarity (Supplementary Fig. **7**). However, when the cue is removed, the attractor reverts to a uniform A state and asymmetry is lost. Thus, while a low feedback regime is optimized for ensuring proper cue-induced asymmetry from divergent initial states, the resulting asymmetry is unstable.

Finally, we considered an oscillatory feedback regime in which cue induction is accompanied by a transient reduction in P→A feedback, similar to what is observed in embryos, where the low feedback state coincides with the time at which symmetry-breaking cues are active (Fig. **3c-e**, Supplementary Videos **4,5**). In this regime, all initial states converge towards a single attractor during the low feedback, cue “on” phase, before evolving towards the AP polarized state (iv) as the system returns to the initial high feedback, cue “off” regime. Thus, by transiently destabilizing the stable attractor states, oscillatory feedback drives convergence of the system from divergent initial states towards a single, stable polarity outcome. For completeness, we examined the behavior of the system when both P→A and A→P feedback were reduced simultaneously (Supplementary Fig. **8**). The results were qualitatively similar, confirming the general benefit feedback oscillations have for promoting responsiveness of patterning systems to spatial cues. Specifically, by destabilizing potential attractors that might otherwise trap the system, oscillatory excursions into low feedback regimes provide transient windows of responsiveness to signal induced changes in state.

Because of the selective nature of the polarity cue in early *C. elegans* embryos, polarization from a uniform A state (e.g. the zygote) is not dependent on oscillation in P→A feedback. However, these results predict that oscillations through a transient low feedback state become critical in two situations: (1) When blastomeres must polarize from a uniform P state or (2) when blastomeres must reverse or reorient a pre-existing polarized state. As we show below, the P blastomeres P1 and P2, respectively, provide clear examples of these two cases, and thus provide an ideal opportunity to test these predictions ^7,11,12,58^.

### Oscillatory feedback facilitates correct response to polarity cues

Polarization of the P blastomeres, P1 and P2, begins with pPARs uniformly occupying the membrane while aPARs remain largely cytoplasmic ^6^. In both cells, an early cue drives pPAR polarization toward the embryo posterior. This cue is linked to a modest enrichment of aPARs at the nascent cell contact site induced by cortical flows during cytokinesis ^11^. However, the two cells diverge in their subsequent polarization patterns. P1 cells possess at least two semi-redundant pathways that reinforce this initial asymmetry ^11,12^, stabilizing polarization of aPARs at the anterior and pPARs at the posterior. In contrast, P2 must override and reorient this initial polarity domain in response to a signal from its sister cell EMS ^7^. This second cue effectively reverses the PAR domains ^58^, leading to enrichment of pPARs at the anterior and aPARs at the posterior. Thus, P1 and P2 reflect ideal test cases for examining whether cue-driven polarization from pPAR high states and polarity reversal, respectively, require a transient period of reduced feedback.

We first focused on P1 polarization as a model for polarization from a uniform pPAR initial state (Fig. **4a,b**, Supplementary Fig. **11,12**). While we were unable to completely eliminate the low feedback state early in the cell cycle, we could significantly truncate it through inhibition of the CDK-1 inhibitor WEE-1 (Fig. **4c**) ^59,59–62^. Consistent with premature CDK-1 activation, when we inhibited WEE-1, embryos exhibited accelerated loading of PAR-1 and CHIN-1 onto the membrane as well as precocious removal of PAR-6 from PAR-2 enriched membranes (Supplementary Fig. **13**). Importantly, both effects were blocked by concomitant inhibition of CDK-1, confirming specificity of the effects on WEE-1 inhibition. These phenotypes were also unaffected by latrunculin A, ruling out contributions from altered cortical actomyosin reorganization that may ectopically induce asymmetries. In otherwise wild-type embryos, we found that WEE-1 inhibition did not prevent polarization, suggesting the initial, albeit shortened, period of low feedback was sufficient in these cases (Supplementary Fig. **13**). However, WEE-1 inhibition rendered embryos sensitive to even modest reductions in aPAR levels. Specifically, when we inhibited WEE-1 in heterozygote *par-6(+/-)* embryos, which otherwise polarize normally, polarization was strongly compromised as scored either by qualitative or quantitative measures of PAR-2 asymmetry (Fig. **4d,e**).

**Figure 4.**
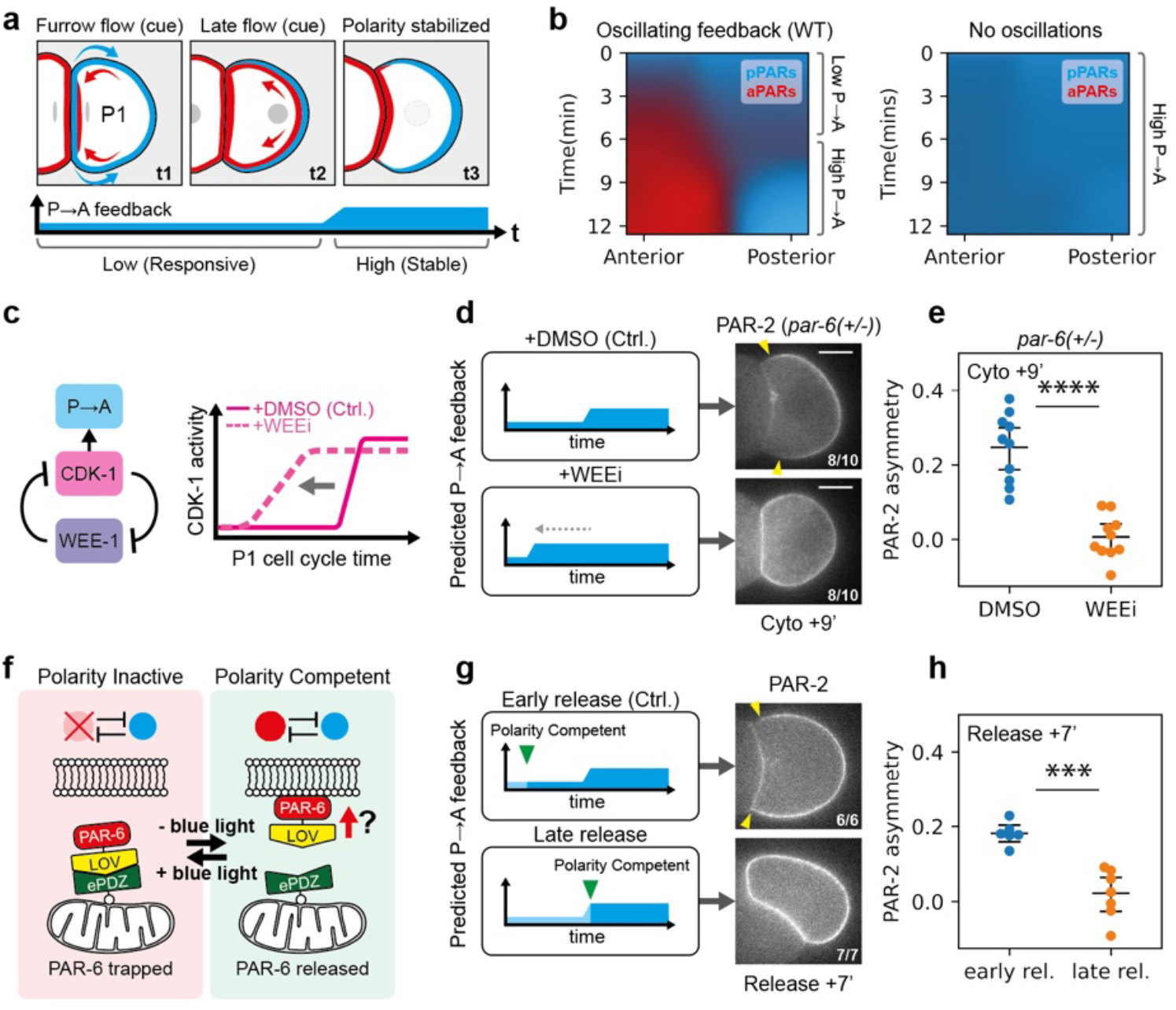
Transient reduction in feedback is required for robust polarization of P1 blastomeres. **a**, Schematic illustrating PAR polarization in P1, alongside the expected dynamics in P→A feedback throughout the cell cycle. **b**, Simulations of a modified PAR model representing P1 polarization suggests a requirement for oscillatory feedback (see Supplementary Fig. **11,12**). **c**, Schematic illustrating regulation between WEE-1, CDK-1 and P→A feedback. Inhibiting the negative regulator of CDK-1, WEE-1 leads to precocious CDK-1 activation, and could shorten the low P→A feedback period. **d**, Left, schematic showing the expected shortening of P→A feedback in acutely WEE-1 inhibited embryos (WEEi; PD0166825) relative to control DMSO treated embryos. Right, representative midplane confocal images of an embryo expressing GFP::PAR-2 in a *par-6(+/-)* background (NWG0323), acutely treated with either DMSO (n=10) or 20μM WEEi (PD0166825) (n=10). Yellow arrowheads indicate the extent of the polarized PAR-2 domain. **e**, the corresponding quantification of PAR-2 asymmetry (ASI) 9 minutes after cell birth. **f**, Schematic illustrating the optogenetic knocksideways approach to trap PAR-6. In the presence of blue light, PAR-6 is sequestered at the mitochondria, preventing access to plasma membrane binding and thus polarity competence. In absence of blue light, PAR-6 is now free to bind to the membrane, making the system competent to polarization. **g**, Left, schematic showing when the system is made polarity competent relative to the dynamic P→A feedback cycles, comparing between early release, where the system is made polarity competent during the period of low P→A feedback, or late release, where the system is made polarity competent after P→A feedback is high. Right, representative midplane confocal images of embryos expressing PAR-6::GFP::LOV, TOMM-20::ePDZ::Halo and GFP::PAR-2 (NWG0597), with mitochondria-trapped PAR-6 released early (∼3mins) (n=6) or late (∼7.5mins) (n=7) after cell birth. Yellow arrowheads indicate the extent of the polarized PAR-2 domain. **h**, the corresponding quantification of PAR-2 asymmetry (ASI) 7 minutes after PAR-6 release.

As an alternative to truncating the low feedback period, we implemented an optogenetic knocksideways approach (Fig. **4f**, Supplementary Fig. **14**, Supplementary Video **6**) ^63,64^. By transiently sequestering PAR-6 at the mitochondrial surface, we could delay when the PAR network becomes competent for polarization and thereby explicitly test whether polarization of P1 must be initiated during the low feedback state. We found that early release during the low feedback period resulted in rapid PAR-6 loading and eventual PAR-2 polarization (Fig. **4g,h**, Supplementary Fig. **14**). By contrast, when we delayed release to a time that corresponded to onset of the high feedback state, PAR-6 was effectively locked out of the membrane and PAR-2 polarization failed.

Next, we used P2 as a model to investigate cue-induced polarity reversal (Fig. **5a,b**). We again utilized WEE-1 inhibition to shorten the initial period of low feedback and performed the experiments in embryos harboring *par-2(MT-)* mutations ^25^, as polarity reversal is more obvious in this background (Fig. **5c,d**) ^11^. Consistent with prior descriptions, control *par-2(MT-)* embryos (treated with DMSO) exhibited initial PAR-2 polarization away from the cell contact towards the posterior (Fig. **5c,d**). This posterior enrichment then rapidly dissipated as PAR-2(MT-) was recruited to the EMS:P2 cell contact. Ultimately, PAR-2(MT-) was consolidated to a single anterior domain in the anterior of P2. WEE-1-inhibited embryos also showed initial posterior enrichment; however, in contrast to control embryos, this posterior PAR-2(MT-) domain persisted even as PAR-2 was recruited to the anterior EMS-P2 contact, leading to the transient coexistence of two PAR-2 domains (Fig. **5c,d**, Supplementary Video **7**). Eventually, the anterior PAR-2(MT-) domain dissipated and was consolidated into the initial posterior domain. In other words, although the effects of the cue are clearly visible in recruiting PAR-2 to the EMS-P2 contact, it cannot outcompete the initial furrow-induced posterior domain, which appears to be locked in by the premature increase in feedback. Thus, polarity reversal fails.

**Figure 5.**
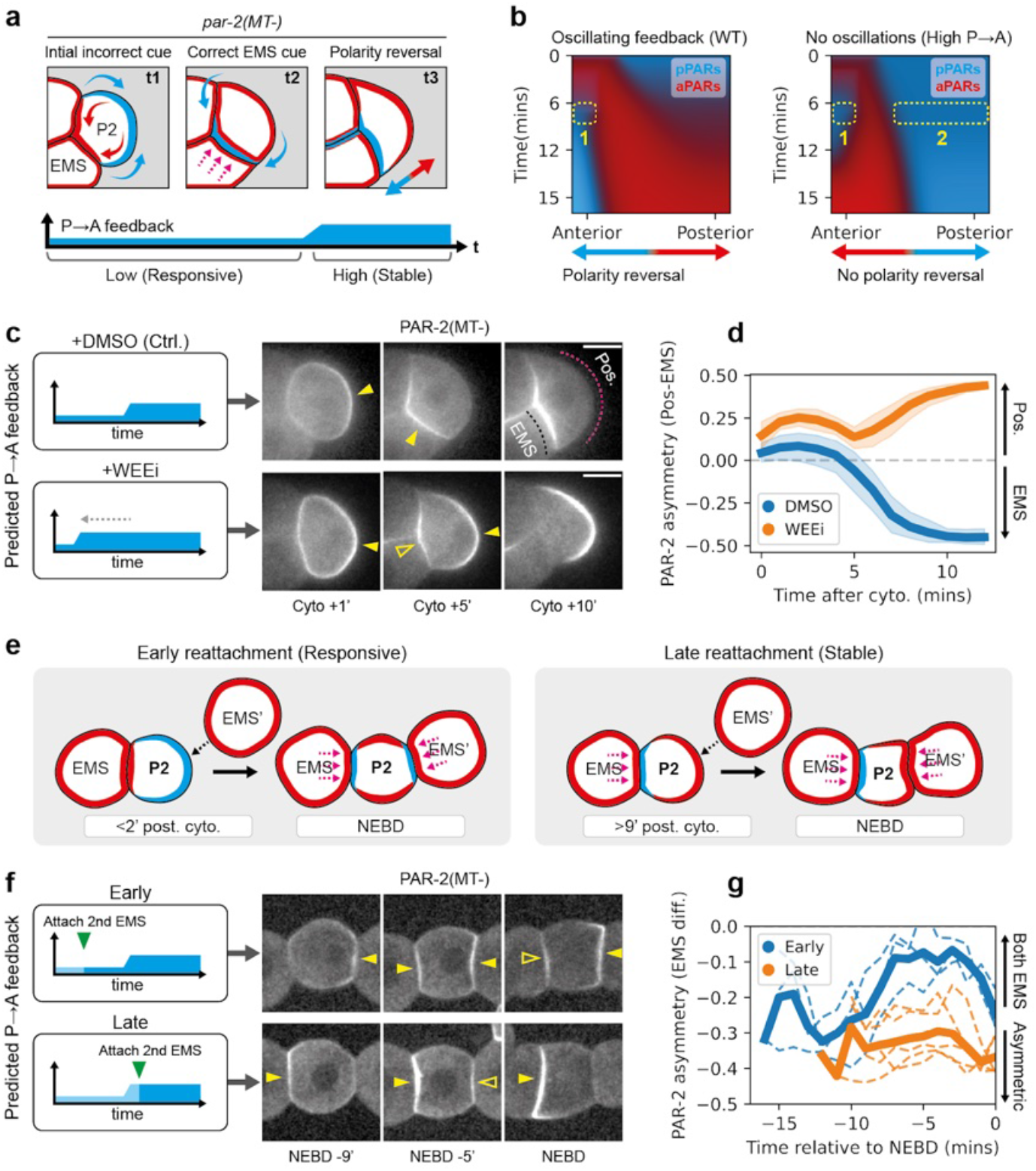
Cue-induced polarity reversal in P2 blastomeres requires a low feedback state. **a**, Schematic illustrating PAR polarization in P2 alongside the expected dynamics in P→A feedback throughout the cell cycle. P2 experiences two distinct cues, an initial incorrect cue that positions aPARs towards the anterior contact site and pPARs away towards the posterior. This is followed by a second cue provided by EMS, which recruits pPARs in the correct direction towards the anterior, effectively reversing the polarity induced by the initial cue. **b**, Simulations of a modified PAR model representing P2 polarization suggests a requirement for oscillatory feedback. Note that the oscillating and constant high feedback system behave differently when the second cue is introduced, while the former exhibits only one pPAR domain at the anterior while the posterior pPAR domain dissipates (dotted rectangles), the latter exhibits two pPAR domains at both the anterior and posterior end respectively. **c**, Left, predicted feedback dynamics in control or WEE-1 inhibited conditions. Right, time series of midsection confocal images of embryos expressing GFP::PAR-2(MT-) and PAR-6::mScarlet-I (NWG0639) (PAR-6 shown in Supplementary Fig. **15**), treated with either DMSO (n=9) or 20μM WEEi (n=8). Yellow closed arrowheads indicate polarized PAR-2(MT-) domains, while yellow open arrowheads indicate the appearance of a second smaller PAR-2(MT-) domain. Magenta and black dotted lines indicate the region used for quantifying PAR-2 at the EMS-contact or embryo posterior in **d. d**, PAR-2 membrane asymmetry over time between the EMS-contact and embryo posterior after cytokinesis completion (cell birth) for DMSO vs WEEi treated embryos. Mean and 95% confidence interval (bootstrapped) indicated. **e**, Schematic illustrating embryo attachment experiments. Dissected EMS-P2 cell pairs were attached with a second ectopic EMS, EMS’, either early in the cell cycle or late. **f**, Left, predicted feedback levels in P2, when the ectopic EMS’ is attached early or late. Right, time series of midsection confocal images of dissected embryos expressing GFP::PAR-2(MT-) (NWG0192) attached with an ectopic EMS’ early (n=3) or late (n=5). Yellow closed arrowheads indicate the presence of PAR-2(MT-) domains, while yellow open arrowheads indicate a second smaller PAR-2(MT-) domain. **g**, quantifications for conditions corresponding to **f**. Mean and results for individual experiments are shown, due to the variable cell cycle stage following completion of EMS’ attachment. Scale bars, 20μm.

To corroborate the WEE-1 inhibition experiments in P2, we turned to isolated blastomeres, which allowed us to control precisely when P2 received signals from EMS (Fig. **5e**). Specifically, isolated EMS–P2 cell pairs were brought into contact with a second, ectopic EMS (EMS′), positioned opposite the native EMS, either early in the P2 cell cycle (0–2 min, when feedback is low) or late (9–11 min, when feedback is high) (Fig. **5f,g**). In early P2 cells, PAR-2(MT-) exhibited a bias away from the native EMS contact, presumably due to furrow-induced asymmetry of aPARs. Attachment of an ectopic EMS′ to these early P2 cells typically produced PAR-2(MT-) domains of roughly equal size at both EMS contacts (ASI shifts towards 0), consistent with a robust polarization response to both cues (Fig. 5f,g, Supplementary Fig. **15)**. In late P2 cells, PAR-2(MT-) already showed a slight bias towards the native EMS contact. In this case, ectopic EMS′ attachment also generated an additional PAR-2(MT-) domain, consistent with P2 still receiving a signal (Supplementary Fig. **15)**. However, this secondary domain was unable to compete with the native domain: the domain was markedly weaker than the original and eventually dissipated, with PAR-2(MT-) ultimately consolidating at the native contact by NEBD. These findings are consistent with cue-sensitivity being limited to the early phase of the P2 cell cycle, which may explain previous reports that the ability of EMS to reorient pPAR domains was restricted to a narrow time window after birth of P2 ^7^.

Importantly, these behaviours were well captured by our model when modified to account for the particular features of P1 and P2, suggesting that changes in feedback dynamics alone are sufficient to account for the observed phenotypes (Fig. **4b**, **5b**, Supplementary Fig. **16-18**).

Taken together, our data demonstrate that the cycling of the PAR network into a low feedback regime early in the cell cycle creates a critical cue-sensitive window that allows cells to properly polarize from diverse initial states and reorient polarity in response to directional cues.

## Discussion

Coupling between cell polarity and the cell cycle has been observed in a broad range of biological systems from bacteria to humans ^26,27,37,46,65–73^. However, the role for such coupling is not clear. In fact, mitotic destabilization of polarity has been reported to sensitize proliferative epithelia to oncogenic transformation due the associated disruption of tissue integrity ^73^, raising the question of why transient destabilization is so conserved. Our data indicates that coupling of feedback in the PAR network to CDK-1 activity cycles the system between low and high feedback states, allowing temporal optimization of network activity. The resulting dynamic polarity landscape facilitates cellular responses to guiding cues by transiently reducing barriers to polarity state switching.

Combining experiments and theory, we show that high antagonistic feedback has opposing effects on cue-driven polarization. On one hand, strong feedback imposes a switch-like behavior in the system, locking in patterns and committing the cell to a specific polarity configuration. However, this comes at the cost of rendering embryos refractory to new cues. Oscillation of PAR activity resolves this conflict by transiently lowering feedback, allowing cells to enter a rheostat-like regime in which the network is highly sensitive to signals and asymmetries largely reflect spatial inputs. These asymmetries can then be reinforced and stabilized as feedback increases again. By temporally ordering periods of network sensitivity and stability, the introduction of cell cycle oscillations in network activity allows cells to sustain robust polarization, while providing for transient periods of cue-sensitivity that allow tight coordination between polarity and morphogenesis.

This paradigm of resolving conflict between cue-sensitivity and pattern stability via oscillations has roots in theoretical work on reaction-diffusion based models of cell polarity ^74–78^. Like the PAR system, these models often rely on stabilizing feedback to support stable polarization in response to weak, transient cues. However, self-stabilizing feedback similarly renders these polarity models resistant to reorientation by secondary cues, posing a problem for systems that must adapt to changing signals ^76,77,79^. One solution is to incorporate delayed negative feedback into the network, which renders polarity fronts self-limiting. Conceptually similar to what we describe in this study, the resulting periodic destabilization of polarity in these so-called excitable systems provides an opportunity to repolarize in a new direction. In chemotaxing cells, it is thought that such oscillatory behaviour balances exploration (sensitivity) and persistence (stability), enabling them to accurately track time-varying signals ^74,76,77,80^. Although these free-running oscillations are ideal for exploratory chemotaxing cells, it is less compatible for systems that need to strictly coordinate polarity behavior with developmental events. In these cases, such as the PAR system, entrainment of oscillations to a developmental controlled clock such as the cell cycle provides an attractive solution, as it allows coupling of the sensitivity of the system to guiding cues with developmental events, e.g. cell division.

The widespread prevalence of links between the cell cycle and cell polarity networks suggests that this paradigm of cell cycle-dependent feedback oscillations will be a common solution for balancing the stability of polarized states with their sensitivity to spatial cues, even if the precise timing and mechanisms are not strictly conserved. This includes stem cell systems such as neuroblasts which must coordinate polarity with niche signals ^37,69,81–84^, dividing epithelial cells which must ensure proper coordination of both apical basal and planar cell polarity with the surrounding tissue ^67,68,72,73,85–89^, and even budding yeast which must integrate spatial cues to ensure proper bud site selection ^90–92^. In each case, we speculate that low feedback regimes provide critical windows for priming cells to respond and adapt polarity guiding cues - in some cases effectively resetting polarity - before cells commit to a stable polarized state that is robust to perturbation.

While we have focused on changes in CDK-1-dependent regulation of pPAR to aPAR feedback, it is important to note that most PAR proteins exhibit some form of cell cycle dependent behavior ^15,23,26,37,81,93–95^, and we cannot rule out similar effects of feedback oscillations in aPARs. Both PAR-3 and CDC-42 behavior vary in a cell cycle dependent manner, though direct evidence for changes in aPAR to pPAR (A→P) feedback is currently lacking. Our theoretical analysis indicates that such coincident oscillation of both P→A and A→P feedback has similar effects on balancing stability and sensitivity, and, in fact, tends to further enhance and generalize the sensitivity of the PAR network to spatial cues, particularly to those that may not act via aPARs as we consider here.

Overall, we suggest that oscillatory signaling dynamics offer systems a way to temporally modulate network dynamics and thus satisfy the often paradoxical requirement for cells to exhibit sensitivity to signals and stability in outputs. This dynamical systems view of signalling network behavior giving rise to a time-varying polarization landscape, in this case entrained by the cell cycle, is particularly attractive for understanding how cells optimize their ability to sense, respond, and adapt to the continuously changing demands of developmental programmes.

## Supporting information

Supplemental Materials - Compiled

## Acknowledgements

We thank Mohit Dalwadi, Jens Januschke, Nic Tapon, and the Goehring lab for comments on the manuscript and advice on theory and experimental design. We thank Miguel Zamora Porras and Neil McDonald for help with preliminary biochemistry experiments, Dan Dickinson, Tobias Dansen, Sander van den Heuvel, and Erik Griffin for strains, and Silvia Santos and Richard Poole for advice and reagents. Some strains were provided by the Caenorhabditis Genome Center (CGC), which is funded by the NIH Office of Research Infrastructure Programs (P40 OD010440).

## Funding

The Francis Crick Institute CC2119 (NWG)

Cancer Research UK CC2119 (NWG)

UK Medical Research Council CC2119 (NWG)

Wellcome Trust CC2119 (NWG)

EMBO ALTF 1053-2022 (ZM)

Canadian Institutes of Health Research Project Grant PJT-169145 (KS)

Natural Sciences and Engineering Research Council of Canada Discovery Grant: RGPIN-2019-04442 (KS)

Michael Smith Health Research BC Scholar Award SCH-2020-0406 (KS)

AC was supported by a PhD Studentship jointly funded by King’s College London and the Francis Crick Institute

## Author Contributions

Conceptualization: KN, NWG

Methodology: KN, AC, ZM, TB

Resources: KN, NH

Investigation: KN, HS, KS

Funding acquisition: KS, ZM, NWG

Project administration: NWG

Supervision: NWG

Writing – original draft: KN, NWG

Writing – review & editing: All authors

## Data and materials availability

All source data are included in the manuscript or Supplementary material. All code will be made available at https://github.com/goehringlab/cdk-par-paper. Materials and all other data will be provided by the lead author upon request.

