## Supplemental Materials - Compiled for "Cell cycle oscillations in a polarity network facilitate state switching by morphogenetic cues"

### Supplementary Materials

Material and Methods

Supplementary Figures S1-18
Supplementary Tables S1-3
Supplementary Videos 1-7

Modelling Supplement

### Materials and Methods

| REAGENT or RESOURCE | SOURCE | IDENTIFIER |
| --- | --- | --- |
| Bacteria | | |
| E. coli: OP50: E. coli B, uracil auxotroph | CGC | WB Strain: OP50 |
| E. coli: HT115(DE3): F-, mcrA, mcrB, IN(rrnD-rrnE)1, rnc14:: Tn10(DE3 lysogen: lavUV5 promoter-T7 polymerase). | CGC | WB Strain: HT115(DE3) |
| Chemicals, Peptides, and Recombinant Proteins | | |
| Chemically defined lipid concentrate | ThermoFisher | 11905031 |
| PP1 Analog, 1NA-PP1 | Merck | 529579 |
| PP1 Analog II, 1NM-PP1 | Merck | 529581 |
| Nocodazole | Merck | M1404 |
| Latrunculin A | Enzo | BML-T119-0100 |
| PD0166285 | | |
| Alisertib | Selleckchem | 1028486-01-2 |
| Alt-R™ S.p. Cas9 Nuclease V3 | IDT | 1081058 |
| Alt-R® CRISPR-Cas9 tracrRNA | IDT | 1072532 |
| Experimental Models: Organisms/Strains | | |
| *C. elegans*: KK1254: *par-2(it315[mCherry::par-2]) III* | Ken Kempheus | WB Strain: KK1254 |
| *C. elegans*: KK1273: *par-2(it328[GFP::par-2]) III* | CGC/Ken Kempheus | WB Strain: KK1273 |
| *C. elegans*: LP216: *par-6(cp45[par-6::mNeonGreen::3xFlag + LoxP unc-119(+) LoxP]) I; unc-119(ed3) III* | Dan Dickinson | WB Strain: LP216 |
| *C. elegans*: LP637: *par-2(cp329[mNG::par-2]) III* | Dan Dickinson | WB Strain: LP637 |
| *C. elegans*: N2: wild type | CGC | WB Strain: N2 |
| *C. elegans*: NWG0042: *par-2(it315[mCherry::par-2]) III; par-1(it324[par-1::gfp::par-1 exon 11a]) V* | This paper | |
| *C. elegans*: NWG0132: *par-2(it315[mCherry::par-2]) III; lon-1(e185) par-3(it71) /qC1[dpy-19(e1259) glp-1(q339) qIs26] III; par-1(it324[par-1::gfp::par-1 exon 11a]) V* | This paper | |
| *C. elegans*: NWG0141: *par-6(tm1425)/In(ile-1 Y18D10A.2 In(dnj-27 dkf-1)) [unc-75(Pmyo-2::Venus)]) I* | Rodrigues et al | |
| *C. elegans*: NWG0192: *par-2(crk30[par-2(R183-5A)::gfp]*KK1273)* | Ng et al | |
| *C. elegans*: NWG0268: *par-6(cp45[par-6::mNeonGreen::3xFlag + LoxP unc-119(+) LoxP]) I; par-2(it315[mCherry::par-2]) III; unc-119(ed3) III?* | Ng et al | NWG0268 |

| | |
|---|---|
| *C. elegans*: NWG0323: *par-6(tm1425)/In(ile-1 Y18D10A.2 In(dnj-27 dkf-1)) [unc-75(Pmyo-2::Venus)]) I; par-2(it328[GFP::par-2]) III* | Rodrigues et al |
| *C. elegans*: NWG0332: *par-2 (it315[mCherry::par-2]) III; par-1(ax4206) V* | Ng et al |
| *C. elegans*: NWG0365: *cdk-1(crk101[cdk-1(F98A)]) III* | This paper |
| *C. elegans*: NWG0434: *par-2(it315[mCherry::par-2]) III; lon-1(e185) par-3(it71)/ qC1[dpy-19(e1259) glp-1(q339) qIs26] III; unc-119(ed3) III (?); itIs37 IV; stIs10226 (?); par-1(it324[par-1::gfp::par-1 exon 11a]) V* | This paper |
| *C. elegans*: NWG0441: *par-6(cp45[par-6::mNeonGreen::3xFlag + LoxP unc-119(+) LoxP]) I; par-2(it315[mCherry::par-2]) III; plk-1(egx3[C52V, L115G]) III; unc-119(ed3) III?* | This paper |
| *C. elegans*: NWG0443: *par-6(cp45[par-6::mNeonGreen::3xFlag + LoxP unc-119(+) LoxP]) I; par-2(it315[mCherry::par-2]) III; cdk-1(crk101[cdk-1(F98A)]); unc-119(ed3) III?* | This paper |
| *C. elegans*: NWG0451: *chin-1(crk146[mNG::CHIN-1]) III* | This paper |
| *C. elegans*: NWG0455: *chin-1(crk146[mNG::CHIN-1]) III; unc-119(ed3) III (?); itIs37 IV; stIs10226 (?)* | This paper |
| *C. elegans: NWG0467: chin-1(crk146[mNG::CHIN-1]) III; lon-1(e185) par-3(it71)/ qC1[dpy-19(e1259) glp-1(q339) qIs26] III* | This paper |
| *C. elegans*: NWG0468: *chin-1(crk146[mNG::CHIN-1]) III; lon-1(e185) par-3(it71)/ qC1[dpy-19(e1259) glp-1(q339) qIs26] III; unc-119(ed3) III (?); itIs37 IV; stIs10226 (?)* | This paper |
| *C. elegans*: NWG0471: *par-2(it315[mCherry::par-2]) III; lon-1(e185) par-3(it71) cdk-1(crk160[cdk-1(F98A)])/ qC1[dpy-19(e1259) glp-1(q339) qIs26] III; par-1(it324[par-1::gfp::par-1 exon 11a]) V* | This paper |
| *C. elegans*: NWG0474: *chin-1(crk146[mNG::CHIN-1]) III; lon-1(e185) par-3(it71) cdk-1(crk160[cdk-1(F98A)])/ qC1[dpy-19(e1259) glp-1(q339) qIs26] III* | This paper |
| *C. elegans*: NWG0509: *nmy-2(crk181[nmy-2(L981P)]) I; par-6(cp45[par-6::mNeonGreen::3xFlag + LoxP unc-119(+) LoxP]) I; par-2(it315[mCherry::par-2]) III; unc-119(ed3) III?* | This paper |
| *C. elegans*: NWG0518: *chin-1(crk146[mNG::CHIN-1]) III; cdk-1(crk101[cdk-1(F98A)]) III* | This paper |
| *C. elegans*: NWG0520: *par-2(it315[mCherry::par-2]) III; cdk-1(crk101[cdk-1(F98A)]) III; par-1(it324[par-1::gfp::par-1 exon 11a]) V* | This paper |
| *C. elegans*: NWG0528: *nmy-2(cp69[nmy-2::mkate2 + LoxP]) I; chin-1(crk146[mNG::CHIN-1]) III; unc-119(ed3) III (?); itIs37 IV; stIs10226 (?)* | This paper |
| *C. elegans*: NWG0543: *nmy-2(cp69[nmy-2::mkate2 + LoxP]) I; chin-1(crk146[mNG::CHIN-1]) III; lon-1(e185) par-3(it71)/ qC1[dpy-19(e1259) glp-1(q339) qIs26] III; unc-119(ed3) III (?); itIs37 IV; stIs10226 (?)* | This paper |
| *C. elegans*: NWG0559: *par-6(cp45[par-6::mNeonGreen::3xFlag +* | This paper |

| | |
|---|---|
| *LoxP unc-119(+) LoxP]) I; cdk-1(crk101[cdk-1(F98A)]) III; unc-119(ed3) III?* | |
| *C. elegans*: NWG0567: *par-2(it315[mCherry::par-2]) III; cdk-1(crk101[cdk-1(F98A)]) III; par-1(ax4206) V* | This paper |
| *C. elegans*: NWG0584: *par-6(he322 [par-6::gfp(smu-1 introns)::glo-lov]) I; utdSi44[mex-5p::tomm-20::glo-epdz::glo-halo::tbb2(3'UTR)] II* | This paper |
| *C. elegans*: NWG0598: *par-6 (he322 [par-6::gfp(smu-1 introns)::glo-lov])/ In(ile-1 Y18D10A.2 In(dnj-27 dkf-1)) [unc-75(Pmyo-2::Venus)]) I; utdSi44 [mex-5p::tomm-20::glo-epdz::glo-halo::tbb2(3'UTR)] II; par-2(djd7 [mSC::PAR-2]) III* | This paper |
| *C. elegans*: NWG0623: *par-6(djd4 [PAR-6::mScarlet-I::Myc]) I; par-2(cp329[mNG-C1^PAR-2]) III* | This paper |
| *C. elegans*: NWG0629: *par-6 (he322 [par-6::gfp(smu-1 introns)::glo-lov]) I; mSc::aPKC pkc-3(djd15[mSc::Myc::aPKC]) II; utdSi44 [mex-5p::tomm-20::glo-epdz::glo-halo::tbb2(3'UTR)] II* | This paper |
| *C. elegans*: NWG0639: *par-2(crk30[par-2(R183-5A)::gfp]*KK1273)/ sC1(s2023) [dpy-1(s2170) umnIs21] III; par-6(djd4[PAR-6::mScarlet-I::Myc]) I* | This paper |
| *C. elegans*: SV2109: *par-6(he322 [par-6::gfp(smu-1 introns)::glo-lov]) I; RuIs57(Ppie-1::Tub::GFP) V* | Fielmich et al |
| *C. elegans*: TBD298: *utdSi44[mex-5p::tomm-20::glo-epdz::glo-halo::tbb2(3'UTR)] II; unc-119(ed3) III; utdSi43[mex5p::PH::glo-mtagbfp2::co-lov::tbb-2(3'UTR)] V* | De Henau et al |
| *C. elegans*: UTX11: *par-6(djd4 [PAR-6::mScarlet-I::Myc]) I* | Dan Dickinson |
| *C. elegans*: UTX31: *par-2(djd7 [mSC::PAR-2]) III* | Dan Dickinson |
| Oligonucleotides | |
| CDK-1(F98A) sgRNA #1:<br>5' – /AltR1/rCrG rGrUrC rArUrU rArUrG rCrArG rGrArG rArArC rGrUrU rUrUrA rGrArG rCrUrA rUrGrC rU/AltR2/ – 3' | IDT DNA |
| CDK-1(F98A) sgRNA #2:<br>5' – /AltR1/rArU rCrGrU rUrUrC rArArG rUrCrG rArArA rGrArC rGrUrU rUrUrA rGrArG rCrUrA rUrGrC rU/AltR2/ – 3' | IDT DNA |
| CDK-1(F98A) repair template (**PstI restriction site**):<br>5'– ATAGGGTGTT CCATCAACGG CTGTGCGAGA GATCAGCTTG CTCAAAGAGC TGCAGCATCC GAATGTTGTT GGATTGGAAG CGGTCATTAT GCAGGAGAAC CGACTTTTCC TGATCGCCGA ATTCTTGTCT TTCGACTTGA AACGATACAT GGATCAGTTG GGAAAAGATG AATACCTTCC GCTCGA –3' | IDT DNA |
| CDK-1(F98A) FWD ODN genotyping primer:<br>5'–CAACAATCCTTCTCAGCGCG–3' | IDT DNA |
| CDK-1(F98A) REV ODN genotyping primer:<br>5'–GGTGTGCCGAGAACTCTGAA–3' | IDT DNA |

| | |
|---|---|
| NMY-2(L981P) sgRNA #1:<br>5' – /AltR1/rUrU rUrCrG rArGrC rArArC rArArU rUrUrC rUrGrA rGrUrU rUrUrA rGrArG rCrUrA rUrGrC rU/AltR2/ – 3' | IDT DNA |
| NMY-2(L981P) sgRNA #2:<br>5' – /AltR1/rUrU rCrArA rUrUrG rArArU rCrUrC rGrGrU rUrGrA rGrUrU rUrUrA rGrArG rCrUrA rUrGrC rU/AltR2/ – 3' | IDT DNA |
| NMY-2(L981P) repair template (**MspI restriction site**):<br>5'– CGAAAATTGA CGGAGATGGT TAGACATCTC GAAGAGAATC TTGAAGATGA AGAAAGAAGC AGACAGAAAT TGTTGCCTGA AAAAAATTCA ATTGAATCCC GGTTGAAAGA ACTGGAAGCA CAAGGACTCG AGCTTGAAGA TTCTGGAAAC AAG –3' | IDT DNA |
| NMY-2(L981P) FWD ODN genotyping primer:<br>5'–ATGAGACTGCGTGAATGGCA–3' | IDT DNA |
| NMY-2(L981P) REV ODN genotyping primer:<br>5'–GACTCTTTGCGCATCAGCTG–3' | IDT DNA |
| mNG::CHIN-1 sgRNA #1:<br>5' – mU*mU*mU* rGrCrA rGrGrU rArUrG rGrArA rGrArC rGrArG rUrUrU rUrArG rArGrC rUrArG rArArA rUrArG rCrArA rGrUrU rArArA rArUrA rArGrG rCrUrA rGrUrC rCrGrU rUrArU rCrArA rCrUrU rGrArA rArArA rGrUrG rGrCrA rCrCrG rArGrU rCrGrG rUrGrC mU*mU*mU* rU – 3' | IDT DNA |
| mNG::CHIN-1 sgRNA #2:<br>5' – mU*mU*mG* rCrArG rGrUrA rUrGrG rArArG rArCrG rArCrG rUrUrU rUrArG rArGrC rUrArG rArArA rUrArG rCrArA rGrUrU rArArA rArUrA rArGrG rCrUrA rGrUrC rCrGrU rUrArU rCrArA rCrUrU rGrArA rArArA rGrUrG rGrCrA rCrCrG rArGrU rCrGrG rUrGrC mU*mU*mU* rU – 3' | IDT DNA |
| mNG::CHIN-1 left homology primer (for Dickinson Lab mNG plasmid):<br>5' – GATTTTCTTC AGAATTTTAT TTATTTTCTA AGAAATTAGC TCAACTTTGC TCATTTTTCT CCAAATTTCT TCGATTTTTT TGCATTTTCA GTTAAAAAAT CAATAAAAAT CGAATTTTTG CAGGTATGGT CAGCAAAGGC GAGGAAGACA – 3' | IDT DNA |
| mNG::CHIN-1 right homology primer (for Dickinson Lab mNG plasmid):<br>5' – GGAAATTGGA AAATTTGAGA TTTTAGCTTT TCGGATATTT TTAAAGCTTC CAAAACTTGT TGAGCTTGAA AAAAATGACT TACCCGAGCT CCCAGGAGGT CCGTCGTCTT CCATCTTGTA CAGCTCGTCC ATTCCCATAA – 3' | IDT DNA |
| mNG::CHIN-1 internal left primer (for Dickinson Lab mNG plasmid):<br>5' – ATGGTCAGCAAAGGCGAGGAAGACA – 3' | IDT DNA |
| mNG::CHIN-1 internal right primer (for Dickinson Lab mNG plasmid):<br>5' – CTTGTACAGCTCGTCCATTCCCAT – 3' | IDT DNA |
| mNG::CHIN-1 FWD ODN genotyping primer:<br>5' – TCTCGATCGCTGGCACTTTT – 3' | IDT DNA |
| mNG::CHIN-1 REV ODN genotyping primer:<br>5' – ATTCGACTCCCGCACCAAAT – 3' | IDT DNA |
| Recombinant DNA | |

| | | |
|---|---|---|
| Ahringer Feeding RNAi: *perm-1* | Source BioScience | WB Clone: sjj_T01H3.4 |
| Ahringer Feeding RNAi: *ptr-2* | Source BioScience | WB Clone: sjj_C32E8.8 |
| Feeding RNAi: *ctrl* | Rodriguez et al | N/A |
| Ahringer Feeding RNAi: *par-2* | Source BioScience | WB Clone: sjj_F58B6.3 |
| Ahringer Feeding RNAi: *par-6* | Source BioScience | WB Clone: sjj_T26E3.3 |
| Ahringer Feeding RNAi: *cyk-1* | Source BioScience | WB Clone: sjj_F11H8.4 |
| Ahringer Feeding RNAi: *par-4* | Source BioScience | WB Clone: sjj2_Y59A8B.14 |
| Ahringer Feeding RNAi: *par-5* | Source BioScience | WB Clone: sjj_M117.2 |
| Ahringer Feeding RNAi: *spd-5* | Source BioScience | WB Clone: sjj_F56A3.4 |
| Software and Algorithms | | |
| Fiji | https://imagej.net/software/fiji/ | RRID:SCR_002285 |
| Metamorph | Molecular Devices | RRID:SCR_002368 |
| Spectral Autofluorescence Image Correction By Regression (SAIBR) | https://github.com/goehringlab/saibr_fiji_plugin | N/A |
| Python | https://www.python.org/ | 3.8.8; RRID:SCR_008394 |
| TensorFlow | https://www.tensorflow.org/ | |
| Others | | |
| Polybead Microspheres 20.00µm | Polysciences | 18329-5 |
| Polybead Microspheres 18.8µm | Polysciences | 18329 |

### EXPERIMENTAL MODEL AND SUBJECT DETAILS

#### *C. elegans* – strains and culture conditions

*C. elegans* strains were maintained on OP50 bacterial lawns seeded on nematode growth media (NGM) at 20˚C or 15˚C (for experiments involving temperature sensitive mutants) under standard laboratory conditions [96]. Strains harboring optogenetic constructs were grown in a dark box to minimize light exposure. Zygotes were obtained from hermaphrodites unless otherwise noted. Analysis of embryos precludes determination of animal sex.

##### *C. elegans* – transgenic animals

Point mutations were generated using CRISPR-Cas9, based on the protocol published by Arribere et al. [97]. Briefly, tracrRNA (IDT DNA, 0.5 µL at 100 µM) and crRNA(s) for the target (IDT DNA, 2.7 µL at 100µM) with duplex buffer (IDT DNA, 2.8µL) were annealed together (5 min, 95˚C) and then stored at room temperature until required. An injection mix containing Cas9 (IDT DNA, 0.5µL at 10mg/mL), annealed crRNA, tracrRNA, and the repair template (IDT Ultramer) was incubated at 37˚C for 15 min and centrifuged to remove debris (10 min, 13,000 rpm). Young gravid adults were injected along with either a *dpy-10* or *unc-58* co-CRISPR injection marker and mutants were verified by PCR and sequencing [97].

Insertion of mNeonGreen was achieved by first generating two PCR products as before [97,98]. one containing the insert DNA sequence, and another containing an insert with an additional ~100-200 bp homology to the insertion site. The products were then column purified (Qiagen, QIAquick PCR purification kit), mixed in equimolar amounts, denatured by heating to 95°C and annealed through slow cooling to room temperature to generate a pool of products with long single-stranded DNA overhangs that act as the repair template. Similar to above, an injection mix containing Cas9 (IDT DNA, 0.5 µl at 10 mg/ml), annealed crRNA, tracrRNA and the repair template was incubated at 37°C for 15 min and centrifuged to remove debris (15 min, 14,100 g). Young gravid N2 adults were injected along with a dpy-10 co-CRISPR injection marker and mutants identified by PCR and sequence verified. Resulting lines were backcrossed with N2s twice before use.

##### Bacterial strains

OP50 bacteria and HT115(DE3) were obtained from CGC. Feeding by RNAi used HT115(DE3) bacteria strains carrying the indicated RNAi feeding plasmid.

#### METHOD DETAILS

##### *C. elegans* – RNAi

RNAi by feeding was performed according to previously described methods [99]. Briefly, HT115(DE3) bacterial feeding clones were inoculated from LB agar plates to LB liquid cultures and grown overnight at 37ºC in the presence of 50 µg/mL ampicillin (until a fairly turbid culture is obtained). To induce high dsRNA expression, bacterial cultures were then treated with 1 mM IPTG before spotting 150µL of culture onto 60 mm NGM agar plates (supplemented with 10 µg/ml carbenicillin, 1 mM IPTG) and incubated for 24 hr at 20ºC. L3/L4 larvae were then added to RNAi feeding plates and incubated for 16-32 hours at either 20ºC or 25ºC.

#### Imaging – dissection, drug treatment and mounting for microscopy

Embryos were obtained by dissecting adult worms in 8-10µL of egg buffer (118 mM NaCl, 48 mM KCl, 2 mM $CaCl_2$ 2 mM $MgCl_2$, 25 mM HEPES, pH 7.3), and mounted with 18.8 µm (cortex imaging) or 20 µm (midplane imaging) polystyrene beads (Polysciences, Inc.) between a slide and coverslip as in [100], and sealed using VALAP (1:1:1, vaseline:lanolin:paraffin wax).

For acute drug treatment experiments, embryos were first permeabilized using either *ptr-2* or *perm-1* fRNAi. For experiments that do not require acute drug addition, embryos were obtained by dissecting adult worms in 8-10µL of Shelton's Growth Medium (with or without drugs), and mounted with 20.0 µm polystyrene beads between a slide and coverslip and sealed with VALAP as above. For experiments that require acute drug addition, embryos were obtained by dissecting adult worms in 8-10µL of Shelton's Growth Medium (with or without drugs), and mounted with 18.8 µm polystyrene beads between a large and small coverslip sealed on two parallel edges with VALAP as in Goehring et al [100]. Buffer exchange was achieved through capillary action by placing a drop of solution at one side of the sample, and touching a piece of filter paper at the opposite side. The timings and details of drug treatment experiments are listed on the table below.

| Figure | Experiment | Drug details | Wash in timings | Wash out timings |
|---|---|---|---|---|
| **1c**, S2 | Block cell division | Latrunculin A (0.5 µM) | Dissected embryos in buffer containing drug | N/A |
| **1e,f**, S4 | Reversibly inhibiting CDK-1 | 1NA-PP1 (20 µM) | Washed in drug after pronuclear meeting | Washed out drug 10-20 minutes after cells arrest |
| **2a**, S4 | Inhibit CDK-1 activity | 1NA-PP1 (50 µM) | Dissected embryos in buffer containing drug | N/A |
| **2c** | Inhibit CDK-1 activity and perturb cytoskeleton | 1NA-PP1 (50 µM); Latrunculin A (0.5 µM); Nocodazole (1 µg/ml) | Washed in drug after pronuclear meeting | N/A |
| **4d**, S12 | Inhibiting WEE-1 | PD0166285 (20 µM) | Washed in drug during zygotic cytokinesis | N/A |
| **5**, S14 | Inhibiting WEE-1 | PD0166285 (20 µM) | Washed in drug late during P1 cytokinesis (~2 mins before completion of cell division) | N/A |
| S5b | Inhibit PLK-1 activity and perturb cytoskeleton | 1NM-PP1 (20 µM); Latrunculin A (0.5 µM); Nocodazole (1 µg/ml) | Washed in drug after pronuclear meeting | N/A |
| S5c | Inhibit AIR-1 activity and perturb cytoskeleton | Alisertib (20 µM); Latrunculin A (0.5 µM); Nocodazole (1 µg/ml) | Washed in drug after pronuclear meeting | N/A |
| S12 | Inhibiting WEE-1 and CDK-1 simultaneously | PD0166285 (20 µM); 1NA-PP1 (20 µM) | Washed in drug during zygotic cytokinesis | N/A |

| | | | | |
|---|---|---|---|---|
| S12 | Inhibiting WEE-1 and actomyosin cortex | PD0166285 (20 μM); Latrunculin A (0.5 μM) | Washed in PD0166285 during zygotic cytokinesis, followed by Latrunculin A 3 minutes after birth of P1. | N/A |

#### Imaging - isolation and reattachment of P2 blastomeres

Ectopic attachment of EMS to P2 blastomeres at different cell cycle stages was achieved by first isolating P1 blastomeres from two-cell stage embryos as described before [7,101,102]. Briefly, adult worms were dissected in an egg salt buffer and the released zygotes were placed into freshly prepared hypochlorite solution [75% Clorox (Clorox) and 2.5 N KOH] for 40 seconds. Following two washes with Shelton's growth medium [103], embryos were transferred onto the imaging chamber. The eggshell and permeability barrier were removed by repeated mouth pipetting with hand-drawn glass microcapillary tubes (10 μL, Kimble Glass Inc.), yielding two-cell stage embryos devoid of eggshell and permeability barriers. We then separated P1 from AB by further mouth pipetting and waited for P1 division. Subsequently, an isolated EMS cell (second EMS) was attached to the P2 blastomere at distinct cell cycle stages. The early cell cycle stage of P2 was defined as 0–2 min after P1 cytokinesis, whereas late P2 was defined as 9–11 min after P1 cytokinesis.

#### Imaging – acute temperature upshift

Rapid temperature upshift for the *nmy-2(ts)* alleles was achieved by preheating a 100X objective lens to 25.5ºC while maintaining the room temperature at 18.5ºC [11,49]. Embryos were initially mounted onto an objective lens without a temperature collar at ~18.5ºC and zygotes were tracked and imaged until pronuclear migration (PNM). At this point, the objective lens was swapped with the preheated one, a process that took approximately 30 to 60 seconds, and imaging was continued. The disruption of *nmy-2(ts)* activity was confirmed by scoring cytokinesis failure following the temperature upshift.

#### Imaging – setup for optogenetic knocksideways experiments

For the optogenetic experiments, worms were only removed from the dark box immediately before experimentation to minimize light exposure. Dissections were performed under minimal light conditions in a dark room with low-light setting on the dissecting microscope. Optogenetic PAR-6 trapping was induced by exposing embryos to 488 nm blue light in 15s intervals with 1s exposure times. The trapping could be reversed by relieving blue light illumination. The release of PAR-6 from mitochondrial-like structures appeared to occur approximately 2 minutes after the blue light was turned off.

#### Imaging – live imaging

Midsection confocal images were captured on a Nikon TiE with a 100x/1.40 NA oil objective, further equipped with a custom X-Light V1 spinning disk system (CrestOptics, Rome, Italy) with 50 µm slits, Obis 488/561 fiber-coupled diode lasers (Coherent, Santa Clara, CA) and an Evolve Delta EMCCD camera (Photometrics, Tucson, AZ). Imaging systems were run using Metamorph (Molecular Devices, San Jose, CA) and configured by Cairn Research (Kent, UK). Filter sets were from Chroma (Bellows Falls, VT): ZT488/561rpc, ZET405/488/561/640X, ET535/50m, ET630/75m. For the optogenetic experiments, an additional blue light filter (Chroma) was added in the DIC light path, on top of the condenser. Imaging of CHIN-1 clusters in acute wash in experiments was achieved as above, but by acquiring a stack from the cortex to the midplane of the embryo, enabling us to select the plane best in focus for quantification. This is because WEE-1 inhibition causes the cell shape to alter significantly throughout the course of imaging, changing the position of the cortical plane.

Cortical imaging of all other experiments were carried out with a 100x/1.40 NA oil objective on a Nikon TiE microscope equipped with an iLas2 TIRF unit (Roper), a custom-made field stop, 488 or 561 fiber coupled diode lasers (Obis), and an Evolve 512 Delta EMCCD camera (Photometrics), controlled by Metamorph software (Molecular Devices) and configured by Cairn Research. Filter sets were from Chroma: ZT488/561rpc, ZET488/561x, ZET488/561m, ET525/50m, ET630/75m, ET655LP. Images were captured in bright field, GFP/mNG (ex488/ZET488/561m), RFP/mKate/mCherry (ex561/ZET488/561m). Simultaneous imaging of CHIN-1 clusters (on the cortex; mNG fluorophore) and histone (deeper into the embryo; mCherry fluorophore) was achieved by using TIRF illumination on the 488 nm laser, and widefield illumination on the 561 nm laser, the latter of which is captured as a stack. We also tagged these embryos with NMY-2::mKate, allowing us to find the cortical focal plane of the embryo during interphase, before CHIN-1 clusters emerge as a reference point for the cortical plane.

Blastomere reattachment experiments were imaged using a microscope Olympus IX83 (Olympus), equipped with a spinning-disk confocal unit CSU-W1 (Yokogawa), a scientific CMOS camera Prime 95B (Photometrics), a piezoelectric stage NANO-Z (Mad City Labs), a silicon immersion objective UPLSAPO60XS2 (NA1.3, 60X; Olympus), and a beam splitter Optosplit II (Cairn Research), which is controlled by Cellsense Dimension (Olympus). A silicone immersion oil (Z81114; refractive index: 1.406 at 23 °C; Olympus) was used as an immersion medium. Samples were illuminated by a diode-pumped laser with 488 nm wavelength, and imaging was performed with 300 ms camera exposure time and 5 sec intervals.

All embryos were imaged with a 20ºC temperature collar, except for the temperature sensitive experiments, to which the temperature collar was set at 25.5ºC, and for the blastomere dissection experiments, where temperature is set at 22.5ºC.

#### Design of an analog-sensitive CDK-1

We first predicted the position of the gatekeeper site of *C. elegans* CDK-1 by aligning an AlphaFold prediction of *C. elegans* CDK-1 structure (AF-P34556-F1-model_v4) with a crystal structure of human CDK1-cyclin B-Cks2 bound to an ATP-competitive inhibitor (5hq0). Combining this with sequence alignment of CDK-1 across various species, we identified the phenylalanine at position 98 (F98) to be the putative gatekeeper site. Mutating CDK-1(F98) to glycine as was done in other species led to homozygous embryonic lethality, suggesting that CDK-1 kinase activity was strongly perturbed. It was previously reported that an additional mutation of CDK-1(M50V) partially rescued CDK-1(F98G) kinase activity in mice. However, these worms were still homozygous inviable. Following advice by Jens Janushke [37], we introduced a *cdk-1(F98A)* mutation, which yielded homozygous viable worms with embryos that were readily inhibited by the ATP analogue 1NA-PP1. This allele was designated *cdk-1*$^{as}$.

### QUANTIFICATION AND STATISTICAL ANALYSIS

#### Image analysis – segmentation

Different segmentation strategies were employed depending on the cell type. P blastomeres were segmented manually using a custom graphical user interface built in Python. In contrast, most zygotes were segmented using a fully automated pipeline based on a convolutional neural network (CNN) built on the U-net architecture. This approach offers significant advantages over previous methods by achieving high accuracy with DIC images, freeing up a fluorescent channel as no fluorescent markers are required, and enhances segmentation consistency across different lines with varying fluorescent signals. The MobileNetV2 network was chosen as the base model due to its low computational requirements and fast segmentation speed. Transfer learning was applied, training only the upsampling step of the U-net model with our data. Binary masks that demarcate the embryo perimeter, excluding the eggshell, were provided as the training data. Data augmentation was also employed, leading to substantial improvements in validation loss. Notably, the results from automatically segmented images closely match those from manual segmentation when processed with a membrane quantification package developed by Tom Bland, which further refines the regions of interest (ROI). Additionally, the model demonstrated robustness against image artifacts, such as spacer beads and worm debris.

#### Image analysis – defining anterior and posterior poles in the zygote

The overall geometry of the zygote was first defined by fitting the shape of the ROI to an ellipsoid. The anterior and posterior poles were defined as the ROI coordinates nearest to the tip of each side of the major axis, which can be defined using a custom built Python graphical user interface.

#### Image analysis – quantification of membrane profile

Raw or SAIBR processed images were used for quantification. In order to measure cortical concentrations, a 100-pixel-wide (15.5 µm) line following the membrane around the embryo was computationally straightened, and a 20-pixel-wide (3.1 µm) rolling average filter was applied to the straightened image. Intensity profiles perpendicular to the membrane at each position were fit to the sum of a Gaussian component, representing membrane signal, and an error function component, representing cytoplasmic signal, and a constant, representing background signal. Membrane concentrations at each position were calculated as the amplitude of the Gaussian component. This protocol is similar to previously published methods and identical to Ng et al. [9,46,104].

#### Image analysis – alignment of time series data in P1, P2 and P3

Because PAR domains are more variable in position during polarization in P blastomeres, membrane profiles were aligned throughout the cell cycle for each embryo, followed by alignment between embryos to ensure accurate representation of PAR polarization dynamics. To align membrane concentration profiles throughout the cell cycle of an embryo, membrane profiles that were adjacent in time were averaged; individual profiles within that time span were aligned to the mean, and this process was iterated until a lowest mean squared error was obtained. To align membrane profiles between embryos, an averaged membrane profile for time points around NEBD was used as a reference for each embryo, and aligned in a manner identical to above, i.e. the membrane profiles of individual embryos were aligned to the mean of all embryos, and the process was iterated until the lowest mean squared error was achieved. Membrane profiles were also geometrically corrected, through automated tracing of the ROIs in both clockwise and anti-clockwise fashion, and to invert membrane profiles of individual embryos so that the average of all embryos has the lowest mean squared error when aligned.

#### Image analysis – segmentation and quantification of cortical clusters

To segment CHIN-1 cortical clusters, we first subtracted background from the image using a difference of Gaussians approach and then detected the position and size of clusters using the Laplacian of Gaussian method. Cluster intensity can then be inferred from the background subtracted image.

#### Image analysis – background subtraction of PAR-6

Due to the low signal-to-noise ratio of PAR-6, we performed background subtraction on some of the images to enhance visibility on the membrane using a difference of Gaussians approach as before. Importantly, this was done only for visualization, none of the quantifications were performed on these background subtracted images (bg-).

#### Image analysis – ASI

For calculating ASI, the following equation was used: ASI = (A – P) / (2 * (A + P)), where A and P denote the sum of fluorescence signals in the anterior or posterior of the cell respectively.

#### Image analysis – quantifying turnover rate of optogenetic experiments

To quantify the rate of PAR-6 release from mitochondria, we first enhanced the structured PAR-6 signal in the mitochondria by performing a difference of Gaussians background subtraction. This process allowed the pixel intensities of cytoplasmic PAR-6 to approximate a normal distribution centered around zero, while the structured PAR-6 trapped in the mitochondria exhibited a tailed distribution towards higher pixel intensities in the histogram. We observed that the tailed distribution of PAR-6 collapsed to approximately a normal distribution around zero two minutes after blue-light release, which suggests that PAR-6 sequestration at the mitochondria can be rapidly reversed.

#### Statistics

All statistical tests were performed in Python and indicated in the figure legends. Data points are shown along with mean values ± 95% confidence interval (bootstrapped) unless otherwise noted. Reported N are the number of embryos analyzed.

#### Modelling

##### Simplified illustrative PAR system using ODEs

We simplified the PAR system into a set of 4 ordinary differential equations, describing either aPARs or pPARs at the embryo anterior or posterior. We assumed symmetric reaction rates (Table S1) for simplified analysis. The equations were solved using the scipy.odeint function in Python. Briefly, the governing equations are as follows:

$$\frac{\mathrm{d}A_a}{\mathrm{d}t} = \tilde{D}(A_p - A_a) + k_{on}A_{cyto} - k_{off}A_a - k_{AP}P_a^{\alpha}A_a$$

$$\frac{\mathrm{d}A_p}{\mathrm{d}t} = \tilde{D}(A_a - A_p) + k_{on}A_{cyto} - k_{off}A_p - k_{AP}P_p^{\alpha}A_p$$

$$\frac{\mathrm{d}P_a}{\mathrm{d}t} = \tilde{D}(P_p - P_a) + k_{on}P_{cyto} - k_{off}P_a - k_{PA}A_a^{\beta}P_a$$

$$\frac{\mathrm{d}P_p}{\mathrm{d}t} = \tilde{D}(P_a - P_p) + k_{on}P_{cyto} - k_{off}P_p - k_{PA}A_p^{\beta}P_p$$

$$A_{cyto} = \rho_A - \psi\frac{A_a + A_p}{2}$$

$$P_{cyto} = \rho_P - \psi\frac{P_a + P_p}{2}$$

Where $A_a, A_p, P_a, P_p$ define aPARs at the anterior, aPARs at the posterior, pPARs at the anterior, and pPARs at the posterior on the membrane respectively, $\rho_A, \rho_P$ define total aPAR and pPAR pools, $\psi$ define surface-area-to-volume ratio, $\tilde{D}$ define diffusion-like terms, $k_{on}, k_{off}$ define on and off rates, and $k_{AP}, k_{PA}$ define $P \rightarrow A$ and $A \rightarrow P$ feedback respectively. All parameters for the simplified ODE model are shown in Table S1. Importantly, simplifying the system allows us to compute the topological landscape more easily, which is achieved by converting the above equation into a stochastic Euler-Maruyama equation (see Modelling Supplement). More details on the construction of the phase portrait and introduction of the cue can be found in the modelling supplement.

#### A mathematically tractable one-species polarity model based on wave-pinning model

To confirm that oscillatory feedback has the same effects on a mathematically tractable model (i.e. without dimensionality reduction), we constructed a single species polarity model based on the wave-pinning model [105]. The governing equations are:

$$\frac{\mathrm{d}X_a}{\mathrm{d}t} = \tilde{D}(X_p - X_a) + k_{on}X_{cyto} - k_{off}X_a + \gamma X_{cyto}\frac{{X_a}^n}{K^n + {X_a}^n}$$

$$\frac{\mathrm{d}X_p}{\mathrm{d}t} = \tilde{D}(X_a - X_p) + k_{on}X_{cyto} - k_{off}X_p + \gamma X_{cyto}\frac{{X_p}^n}{K^n + {X_p}^n}$$

$$X_{cyto} = \rho_X - \psi\frac{X_a + X_p}{2}$$

Where $X$ represents the polarity species. Positive feedback is in the form of a hill function in which membrane-associated $X$ locally recruits $X_{cyto}$ additionally. $\gamma$ defines the feedback strength and $K$ represents the saturation constant. Here, the quasi-potential is calculated by solving the Fokker-Planck equation, whereas the quivers were calculated using the instantaneous velocity at each system state. See modelling supplement for more information. Parameter values can be found in Table S2.

#### Representative PAR system using PDEs

A full partial differential equation describing polarization from pPAR dominant (P1-like) and reversed polarized states (P2-like) was simulated using forward Euler's method with sufficiently small time steps ($\delta t = 0.01$) using finite difference discretization, due to dynamic changes in P→A feedback. The governing equations were written as:

$$\partial_t A = D_A \partial_x^2 A + k_{on,A}A_{cyto} - k_{off,A}A - k_{AP}P^{\alpha}A$$

$$\partial_t P = D_P \partial_x^2 P + k_{on,P} P_{cyto} - k_{off,P} P - k_{PA} A^\beta P$$
$$A_{cyto} = \rho_A - \psi \bar{A}$$
$$P_{cyto} = \rho_P - \psi \bar{P}$$

Where $\bar{A}, \bar{P}$ define membrane averages of aPARs and pPARs. Note that some of the reaction rates here are no longer symmetric.

Here $k_{AP}, k_{PA}$ values were estimated using experimental data through fitting, see Table S3. We approximated dynamic changes in $k_{AP}$ as a function of changes in measured PAR-1 membrane levels during CDK-1 inhibition or throughout the cell cycle. Dynamic effects resulting from the interaction of smooth changes in feedback level with the cue are likely present and will provide additional properties to the network [106], but are not further explored as it is outside the scope of this study. Further details on the data fitting and model construction can be found in the modelling supplement.

### Supplementary Figures

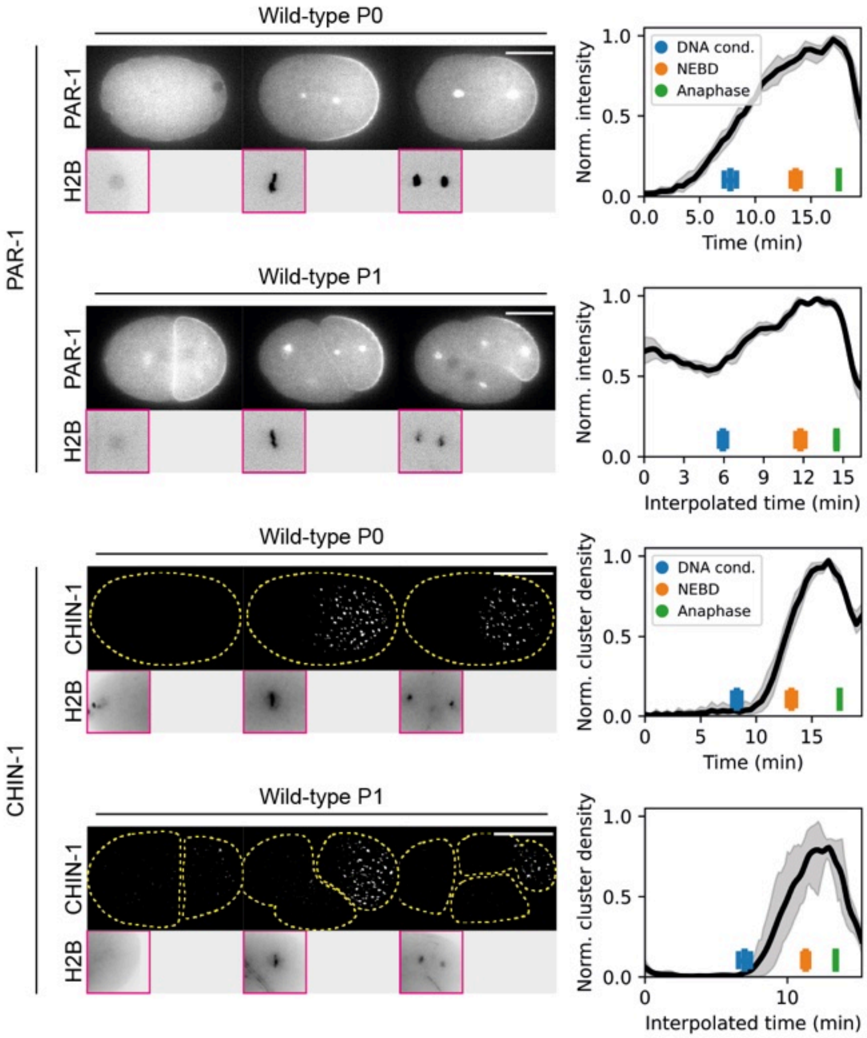

**Supplementary Figure S1. PAR-1 and CHIN-1 membrane levels oscillate through the cell cycle of zygote and P1.**

Top, time series of midplane confocal images of embryos expressing PAR-1::GFP with mCherry::PAR-2 (not shown) and H2B::mCherry in a *par-3(+/-)* background (NWG0434, roller phenotype). Sample sizes: zygote (n=4), P1 (n=5).

Bottom, time series of background subtracted cortical images of embryos expressing mNG::CHIN-1, H2B::mCherry and NMY-2::mKate2 (not shown) (NWG0528). Sample sizes: zygote (n=8), P1 (n=5).

Due to the variable cell cycle length between birth of P1 to completion of cytokinesis, we normalized the cell cycle time of each embryo to the mean of all embryos, and thus temporal PAR-1 membrane profiles were interpolated to match this new normalized time (interpolated time).

Quantifications of corresponding conditions are shown on the right.

Mean and 95% confidence interval (bootstrapped) indicated. Scale bars, 20µm.

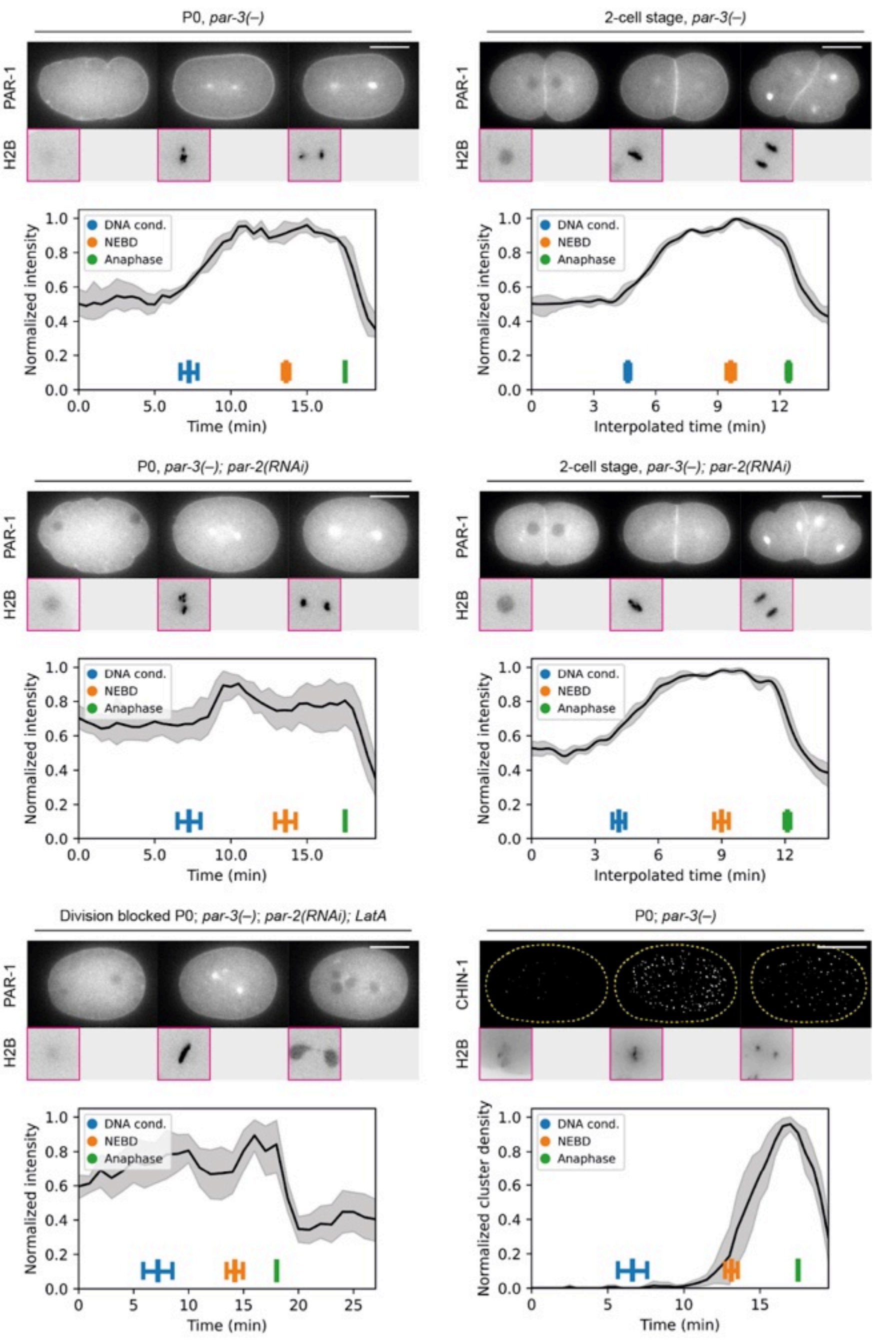
P0, par-3(–)
PAR-1
H2B
DNA cond.
NEBD
Anaphase
Normalized intensity
Time (min)
2-cell stage, par-3(–)
Interpolated time (min)
P0, par-3(–); par-2(RNAi)
2-cell stage, par-3(–); par-2(RNAi)
Division blocked P0; par-3(–); par-2(RNAi); LatA
P0; par-3(–)
CHIN-1
Normalized cluster density

**Supplementary Figure S2. Oscillations of PAR-1 and CHIN-1 membrane levels are still observable when *par-2* and *par-3* were depleted.**

All except bottom right, time series of midplane confocal images of embryos expressing PAR-1::GFP with mCherry::PAR-2 (not shown) and H2B::mCherry in a *par-3(–)* background (NWG0434, roller phenotype), subject to either *ctrl(RNAi)* or *par-2(RNAi)*. The embryo on the bottom left was additionally treated with Latrunculin A to block cell division, allowing visualization of PAR-1 without confounding effects from furrow membranes during cell division. Sample sizes: zygote + *ctrl(RNAi)* (n=4), P1 + *ctrl(RNAi)* (n=5), zygote + *par-2(RNAi)* (n=6), P1 + *par-2(RNAi*) (n=5), zygote + *par-2(RNAi)* + 0.5μM Latrunculin A (n=5).

Bottom right, time series of background subtracted cortical images of embryos expressing mNG::CHIN-1, H2B::mCherry, NMY-2::mKate2 (not shown) in a *par-3(–)* background (NWG0543) (n=4).

Quantifications of corresponding conditions are shown below each image.

Mean and 95% confidence interval (bootstrapped) indicated. Scale bars, 20μm.

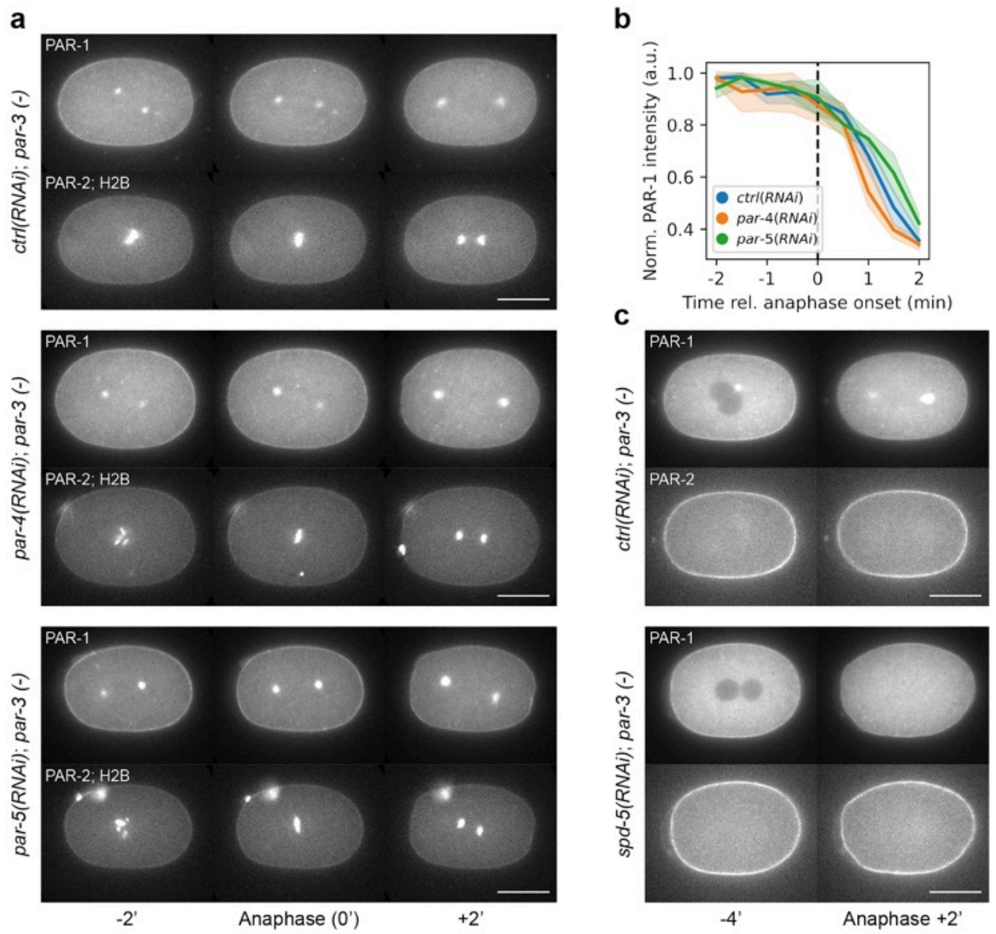
a
PAR-1
PAR-2; H2B
ctrl(RNAi); par-3 (-)
par-4(RNAi); par-3 (-)
par-5(RNAi); par-3 (-)
-2'
Anaphase (0')
+2'
b
Norm. PAR-1 intensity (a.u.)
1.0
0.8
0.6
0.4
ctrl(RNAi)
par-4(RNAi)
par-5(RNAi)
-2
-1
0
1
2
Time rel. anaphase onset (min)
c
PAR-1
PAR-2
ctrl(RNAi); par-3 (-)
spd-5(RNAi); par-3 (-)
-4'
Anaphase +2'

**Supplementary Figure S3. Oscillations of PAR-1 membrane levels are observable when *par-4*, *par-5* and *spd-5* were depleted.**

**a,** PAR-1 regulators PAR-4 and PAR-5 are likely not involved in PAR-1 membrane oscillations, as membrane reduction during anaphase was still observed. Time series of midplane confocal images of embryos expressing PAR-1::GFP with mCherry::PAR-2 in a *par-3(–)* background (NWG0132), subject to either *ctrl(RNAi)* (n=5), *par-4(RNAi)* (n=3) or *par-5(RNAi)* (n=4).

**b,** Quantifications of PAR-1 membrane levels in conditions corresponding to **a**.

**c,** PAR-1 membrane reduction during anaphase was still observed when the centrosomes were disrupted, suggesting PAR-1 sequestration by the centrosomes does not play a key role. Same as in (A), comparing *ctrl(RNAi)* (n=2) and *spd-5(RNAi)* (n=2).

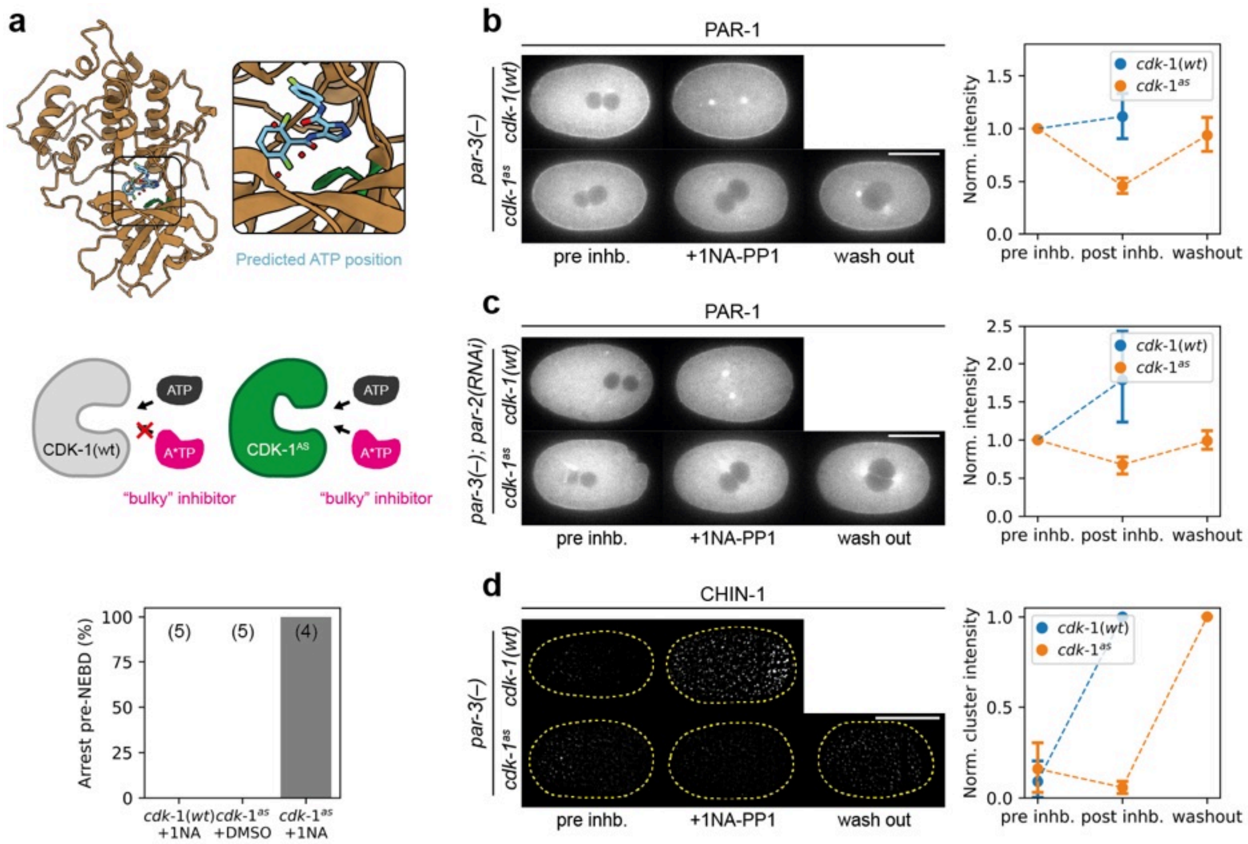
a
Predicted ATP position
ATP
A*TP
CDK-1(wt)
"bulky" inhibitor
ATP
A*TP
CDK-1^AS
"bulky" inhibitor
Arrest pre-NEBD (%)
100
75
50
25
0
(5)
(5)
(4)
cdk-1(wt) +1NA
cdk-1^as +DMSO
cdk-1^as +1NA
b
PAR-1
par-3(−)
cdk-1(wt)
cdk-1^as
pre inhb.
+1NA-PP1
wash out
Norm. intensity
1.5
1.0
0.5
0.0
cdk-1(wt)
cdk-1^as
pre inhb. post inhb. washout
c
PAR-1
par-3(−); par-2(RNAi)
cdk-1(wt)
cdk-1^as
pre inhb.
+1NA-PP1
wash out
Norm. intensity
2.5
2.0
1.5
1.0
0.5
0.0
cdk-1(wt)
cdk-1^as
pre inhb. post inhb. washout
d
CHIN-1
par-3(−)
cdk-1(wt)
cdk-1^as
pre inhb.
+1NA-PP1
wash out
Norm. cluster intensity
1.0
0.5
0.0
cdk-1(wt)
cdk-1^as
pre inhb. post inhb. washout

**Supplementary Figure S4. Inhibition of CDK-1$^{AS}$ and its effect on PAR-1 and CHIN-1 membrane levels in *par-3(-)* embryos.**

**a,** Design and testing of an analog-sensitive *cdk-1* allele (*cdk-1$^{as}$*). Top, AlphaFold predicted structure of *C. elegans* CDK-1 kinase domain superimposed with a solved human CDK-1 kinase domain bound to a competitive ATP inhibitor (light blue), allowing us to identify the ATP binding site in *C. elegans* CDK-1 kinase domains. Green amino acid residue represents the predicted gatekeeper site. Middle, schematic illustrating the rationale of *cdk-1$^{as}$* inhibition. Wildtype CDK-1 kinase domains (grey) can readily accept ATP (dark gray) but not bulky non-hydrolysable ATP analogs (magenta), such as 1NA-PP1. CDK-1$^{AS}$ kinase domains (green) can accept ATP for normal functioning, but also be inhibited by bulky non-hydrolysable ATP analogs, which acts as competitive inhibitors towards ATP. This is due to a mutation at the gatekeeper site to a smaller amino acid residue, which "opens up" the ATP binding pocket of the kinase. Bottom, quantification of pre-NEBD arrest when *cdk-1(wt)* or *cdk-1$^{as}$* embryos were treated with DMSO or 50µM 1NA-PP1. *cdk-1(wt)* + 1NA-PP1 serves as a drug control while *cdk-1$^{as}$* + DMSO serves as an allele control.

**b,** Time series of midplane confocal images of embryos expressing PAR-1::GFP and mCherry::PAR-2 (not shown) in a *par-3(–)* with *cdk-1(wt)* (NWG0132) (n=5) or *cdk-1$^{as}$* (NWG0443) (n=8) background treated with 20µM 1NA-PP1 after PNM and washed out after ~10-20 minutes. Right, quantification of normalized average PAR-1 membrane levels after treating with 1NA-PP1 and drug wash out.

**c,** Same as **b**, but with *par-2(RNAi)*. Sample sizes: *cdk-1(wt)* (n=5), *cdk-1$^{as}$* (n=8).

**d,** Same as **b**, but quantifying normalized background subtracted CHIN-1 cluster levels. Sample sizes: *cdk-1(wt)* (NWG0467; n=4) or *cdk-1$^{as}$* (NWG0474; n=5).

Mean and 95% confidence interval (bootstrapped) indicated. Scale bars, 20µm.

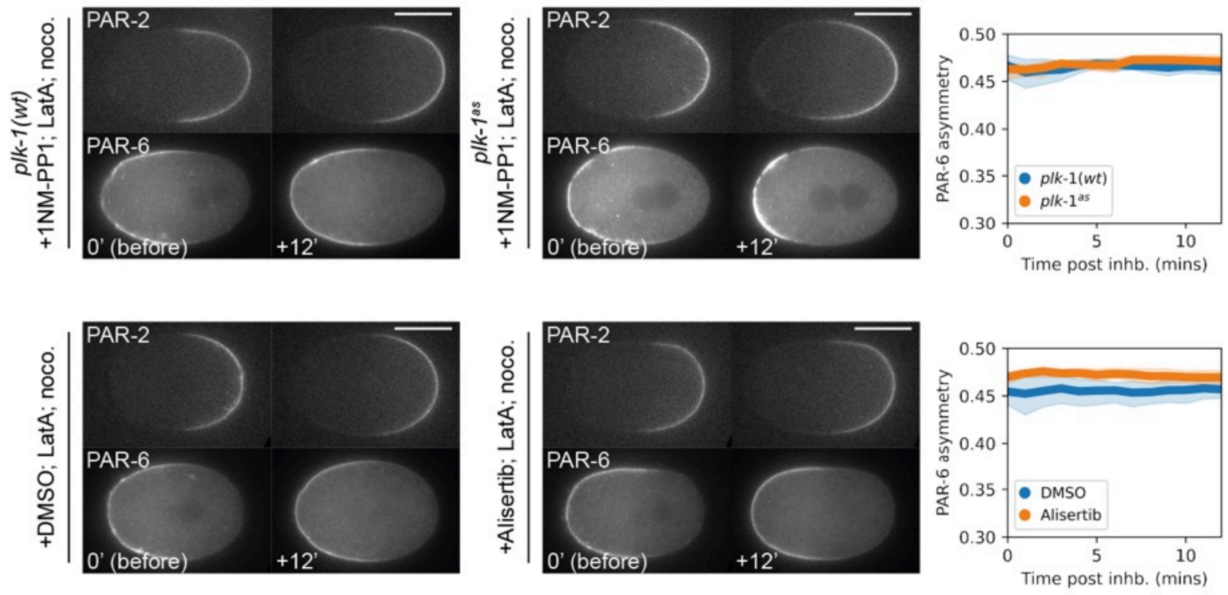
plk-1(wt)
+1NM-PP1; LatA; noco.
plk-1as
+1NM-PP1; LatA; noco.
+DMSO; LatA; noco.
+Alisertib; LatA; noco.
PAR-2
PAR-6
0' (before)
+12'
PAR-6 asymmetry
Time post inhb. (mins)
plk-1(wt)
plk-1as
DMSO
Alisertib
0.50
0.45
0.40
0.35
0.30
0
5
10

**Supplementary Figure S5. Inhibition of AIR-1 and PLK-1 alongside cytoskeletal disruption does not lead to visible PAR-6 invasion into PAR-2 occupied domains.**
CDK-1 inhibition leads to PAR-6 invasion into PAR-2 occupied membranes, and is likely not dependent on changes associated with the cytoskeleton, AIR-1 or PLK-1 activity. Midplane confocal images of embryos expressing mCherry::PAR-2 and PAR-6::mNG in different backgrounds, acutely treated with drugs after pronuclear meeting (PNM): top, *plk-1(wt)* (NWG0268; n=7) or *plk-1$^{as}$* (NWG0441; n=7) background [107] treated with 20μM 1NM-PP1, 0.5μM Latrunculin A, and 1μg/ml nocodazole; bottom, wildtype background (NWG0268) treated with 0.5μM Latrunculin A, 1μg/ml nocodazole, and DMSO (vehicle control; n=5) or alisertib (AIR-1 inhibitor; n=7) [108]. Open white arrowheads indicate ectopic posterior localisation of PAR-6. Right, quantification of average PAR-6 membrane levels at the embryo posterior after drug treatment.
Mean and 95% confidence interval (bootstrapped) indicated. Scale bars, 20μm.

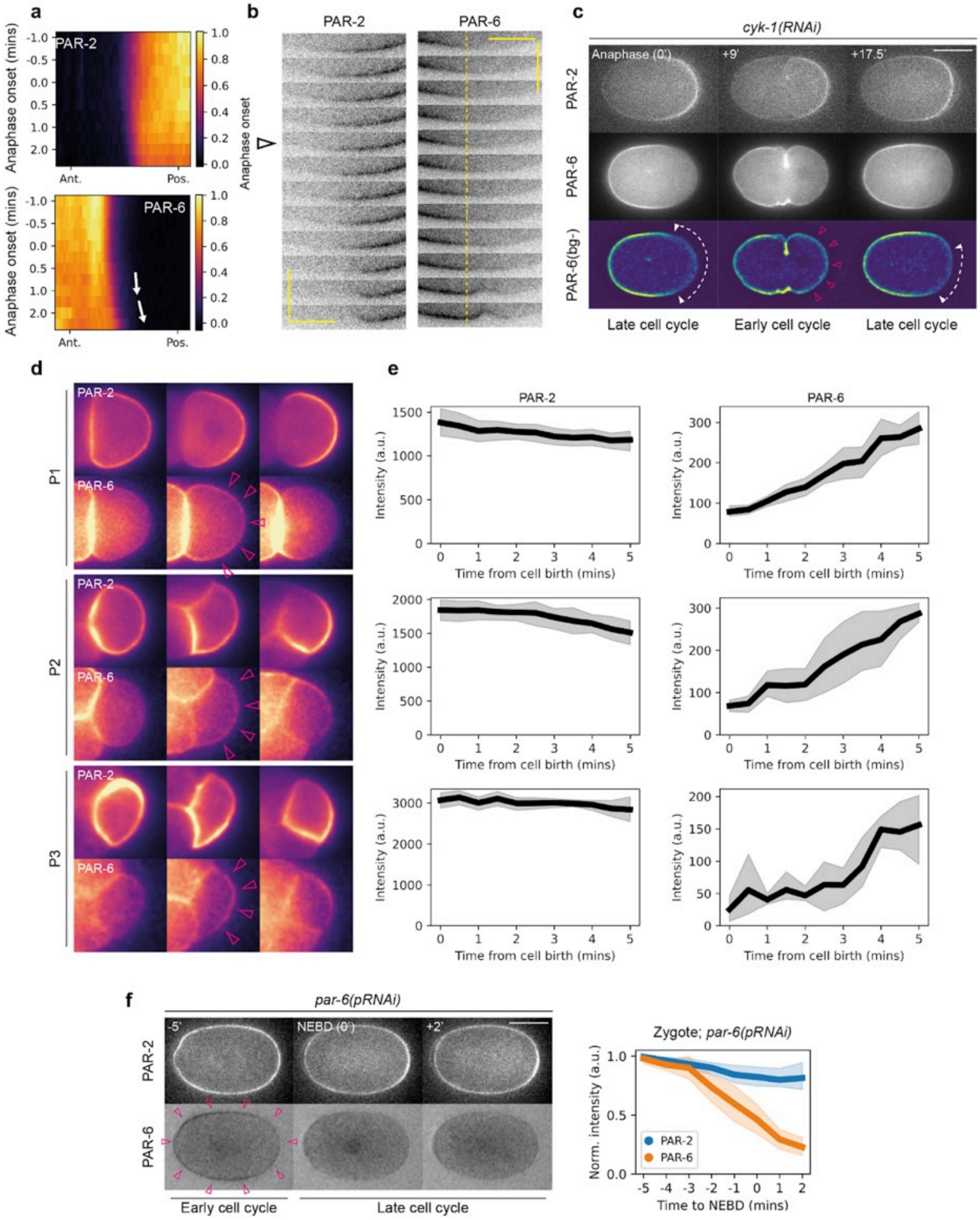
a
PAR-2
PAR-6
Anaphase onset (mins)
-1.0
-0.5
0.0
0.5
1.0
2.0
Ant.
Pos.
1.0
0.8
0.6
0.4
0.2
0.0
b
PAR-2
PAR-6
Anaphase onset
c
cyk-1(RNAi)
Anaphase (0')
+9'
+17.5'
PAR-2
PAR-6
PAR-6(bg-)
Late cell cycle
Early cell cycle
Late cell cycle
d
P1
P2
P3
PAR-2
PAR-6
e
PAR-2
PAR-6
Intensity (a.u.)
Time from cell birth (mins)
f
par-6(pRNAi)
-5'
NEBD (0')
+2'
PAR-2
PAR-6
Early cell cycle
Late cell cycle
Zygote; par-6(pRNAi)
Norm. intensity (a.u.)
Time to NEBD (mins)
PAR-2
PAR-6

**Supplementary Figure S6. Invasion of PAR-6 into PAR-2 domains is correlated with cell cycle stage, and is observable across P blastomeres.**

**a,** Quantification of average PAR-2 and PAR-6 membrane distributions during anaphase onset. White arrows indicate posterior invasion of PAR-6 during anaphase onset, as cells begin to transition to low cell cycle activity.

**b,** Corresponding kymograph corresponding to **a**, indicating a midplane cross section for a region close to the site of furrow formation. Vertical scale bar 30 seconds, horizontal scale bar, 10μm. Dotted yellow line indicates PAR-6 begins to invade into the posterior soon after anaphase onset (Open black arrowhead).

**c,** Time series of midsection confocal images of an embryo expressing mCherry::PAR-2 and PAR-6::mNG (NWG0268) in *cyk-1(RNAi)* conditions (n=3), which allows the zygote to progress through the cell cycle but not cell division. Arrowheads with dotted lines indicate clearance of PAR-6 from PAR-2 occupied membranes late in the cell cycle, magenta open arrowheads indicate loading of PAR-6 onto PAR-2 occupied membranes. Background subtraction of PAR-6 (PAR-6(bg-), see Methods) was performed to improve visibility of phenotype.

**d,** Time series of midsection confocal images of embryos expressing mNG::PAR-2 and PAR-6::mScarlet-I (NWG0623) in P1, P2 and P3 blastomeres (n=6). Magenta open arrowheads show aPAR loading onto the membrane even in the presence of pPARs early in the cell cycle.

**e,** Quantification of PAR membrane loading and distribution in P blastomeres. Note that we have used PAR-6::mNG (LP216) and GFP::PAR-2 (KK1273) for quantification instead - combining the use of a green fluorophore with SAIBR yielded better images. Left, average PAR-2 membrane concentration across the embryo early in the cell cycle of P1 (n=7), P2 (n=6) and P3 (n=6). Right, average PAR-6 membrane concentration across the embryo, outside of cell-cell contact sites, early in the cell cycle of P1 (n=6), P2 (n=4), and P3 (n=3).

**f,** Left, time series of midsection confocal images of an embryo expressing mCherry::PAR-2 and PAR-6::mNG (NWG0268) in *par-6* partial RNAi (*par-6(pRNAi))* conditions (n=4). Note that PAR-6 overlaps with PAR-2 (magenta open arrowhead) until right before NEBD, where PAR-6 membrane levels begin to decrease. Scale bar, 20μm. Right, quantification of PAR-2 and PAR-6 membrane levels for the corresponding conditions.

Mean and 95% confidence interval (bootstrapped) indicated. Scale bars, 20μm.

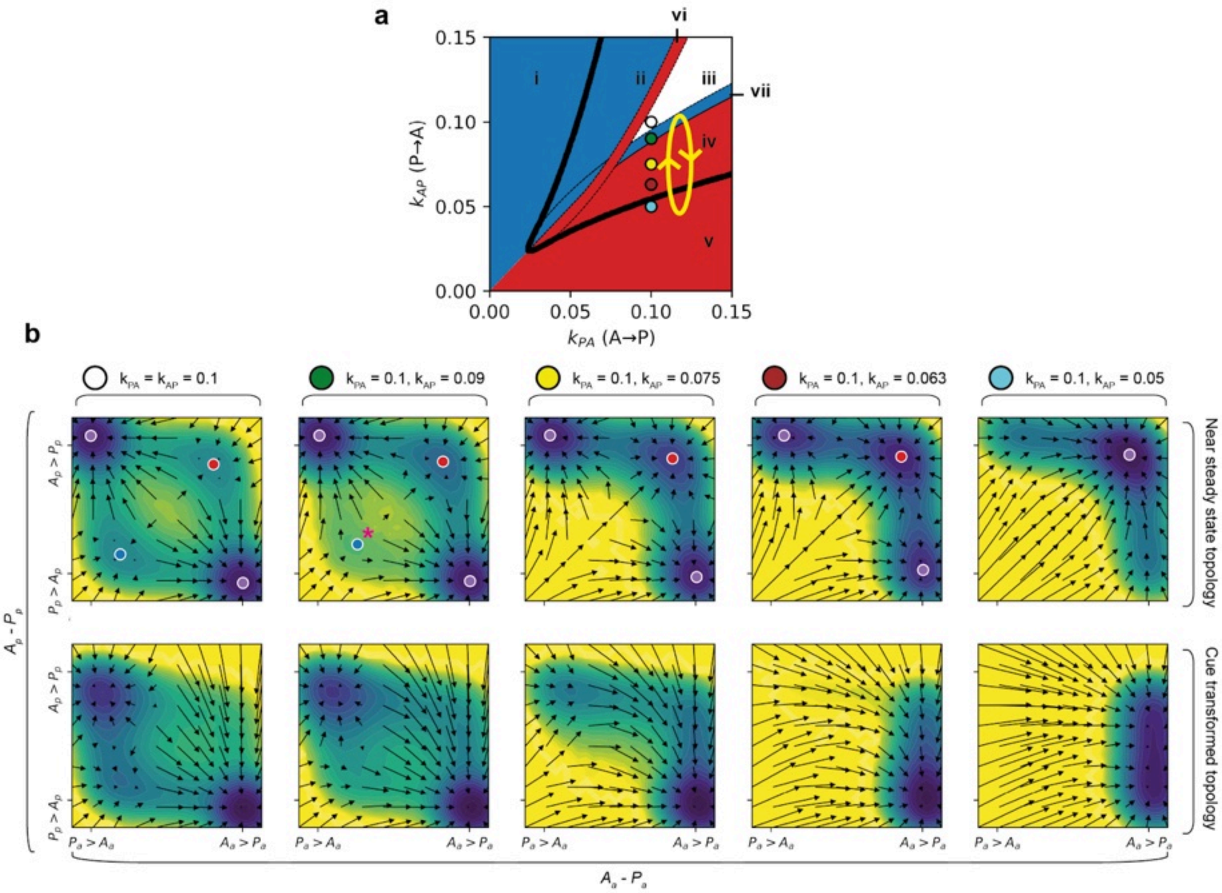
a
0.15
0.10
0.05
0.00
0.00
0.05
0.10
0.15
$k_{AP}$ (P→A)
$k_{PA}$ (A→P)
i
ii
iii
iv
v
vi
vii
b
$k_{PA} = k_{AP} = 0.1$
$k_{PA} = 0.1$, $k_{AP} = 0.09$
$k_{PA} = 0.1$, $k_{AP} = 0.075$
$k_{PA} = 0.1$, $k_{AP} = 0.063$
$k_{PA} = 0.1$, $k_{AP} = 0.05$
Near steady state topology
Cue transformed topology
$A_p - P_p$
$A_p > P_p$
$P_p > A_p$
$P_a > A_a$
$A_a > P_a$
$A_a - P_a$

**Supplementary Figure S7. Landscape of a simplified 2-step discretized PAR model with oscillating $k_{AP}$ (P→A) feedback.**

**a,** Parameter-space topology of a 2-step discretized PAR model (see Methods). Blue region (i, ii) supports pPAR dominant homogeneous states. Red region (iv, v) supports aPAR dominant homogenous states. White region (iii) supports both aPAR and pPAR dominant homogenous states, depending on the initial state. Dotted blue region (vi) indicates region that can undergo spontaneous symmetry breaking if beginning from pPAR initial states. Dotted red region (vii) indicates the same, but beginning from aPAR initial states. Solid black lines indicate regions permissible to stable polarization (ii, iii, iv, vi, vii). This topology is similar to previously described works [8,52]. Colored circular points with black outlines indicate points that were sampled and further examined using a concentration difference landscape in **b**, representing what the system experiences when it undergoes $k_{AP}$ (P→A) oscillations.

**b,** Examination of system behavior at changing $k_{AP}$ levels using a phase plane of concentration differences, between aPAR and pPAR at the anterior ($A_a$ - $P_a$) for the x-axis, and between aPAR and pPAR at the posterior ($A_p$ - $P_p$) for the y-axis. Top, near steady state topology of the system (see Methods). Dotted circles represent steady state points for either polarized states (purple), homogenous aPAR high states (red) or homogenous pPAR high states (blue). Magenta asterisk represents an unstable steady state. Bottom, topology of the system after transformation with an aPAR acting cue.

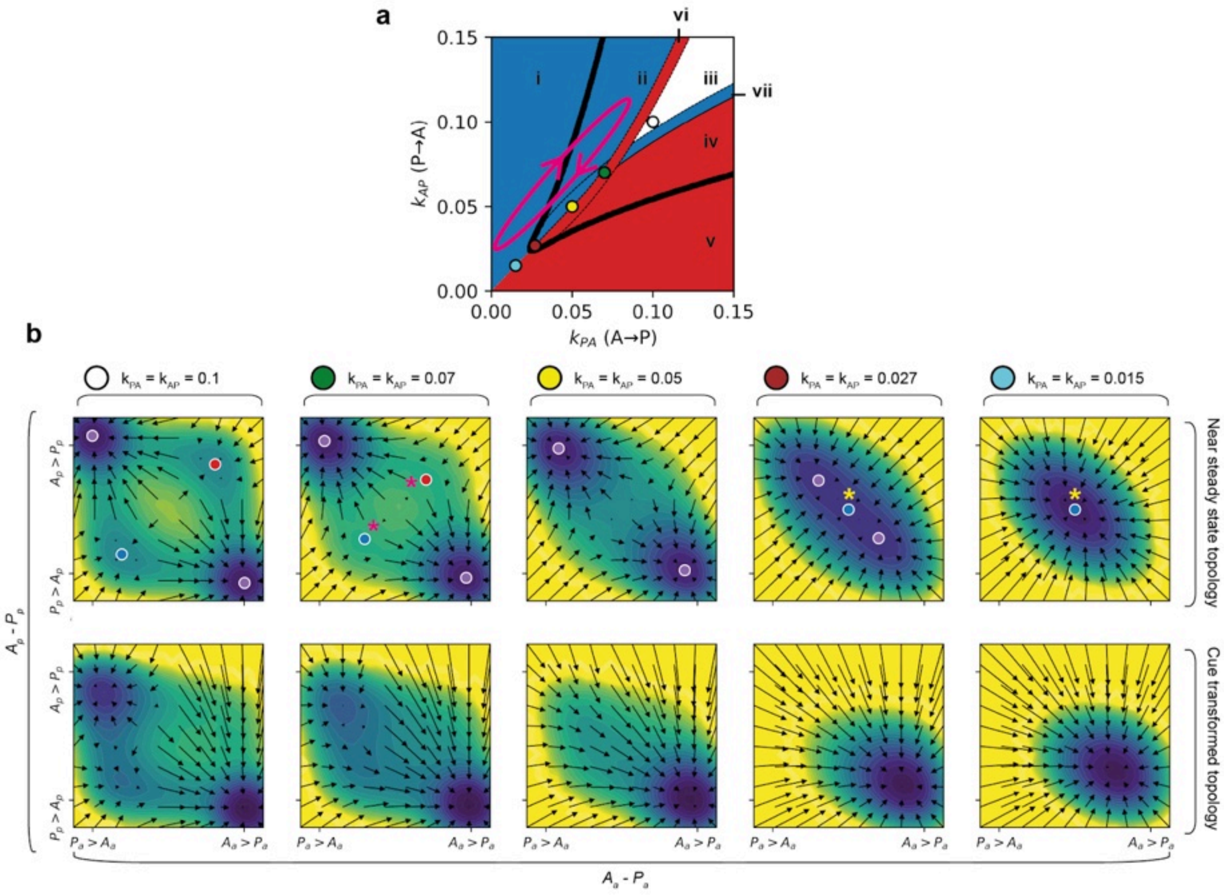
a
vi
i
ii
iii
vii
iv
v
0.15
0.10
0.05
0.00
0.00 0.05 0.10 0.15
$k_{AP}$ (P→A)
$k_{PA}$ (A→P)
b
$k_{PA} = k_{AP} = 0.1$
$k_{PA} = k_{AP} = 0.07$
$k_{PA} = k_{AP} = 0.05$
$k_{PA} = k_{AP} = 0.027$
$k_{PA} = k_{AP} = 0.015$
Near steady state topology
Cue transformed topology
$A_p - P_p$
$A_p > P_p$
$P_p > A_p$
$P_a > A_a$
$A_a > P_a$
$A_a - P_a$

**Supplementary Figure S8. Landscape of a simplified 2-step discretized PAR model, oscillating both kAP (P→A) and kPA (A→P) feedback simultaneously.** Magenta asterisk represents unstable steady states. Yellow asterisk represents the stable point of the system which is homogenous high for both pPARs and pPARs, with both species overlapping on the membrane.

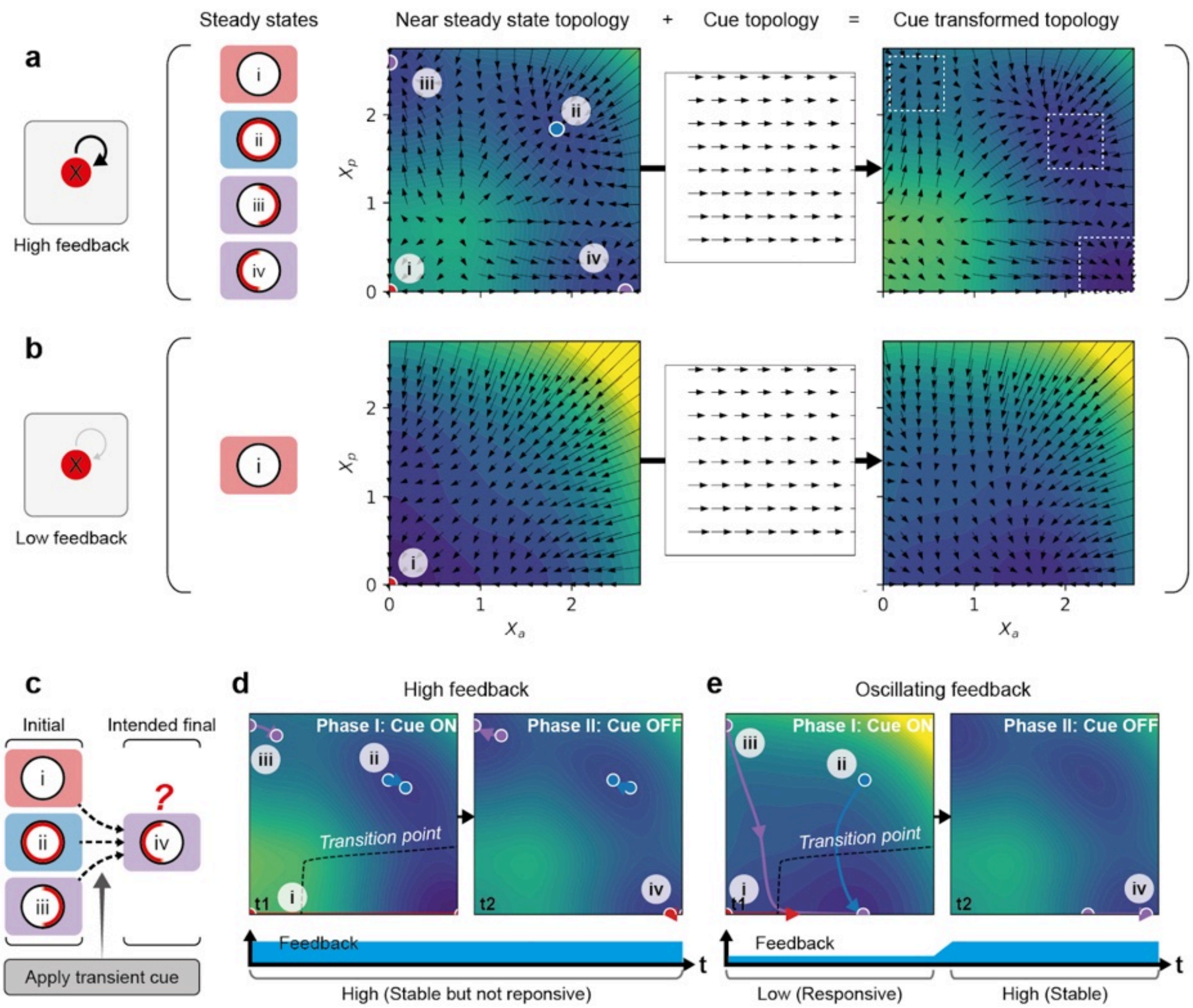
a
b
c
d
e
Steady states
Near steady state topology
+
Cue topology
=
Cue transformed topology
High feedback
Low feedback
i
ii
iii
iv
$X_p$
$X_a$
0
1
2
Initial
Intended final
Apply transient cue
High feedback
Oscillating feedback
Phase I: Cue ON
Phase II: Cue OFF
Transition point
t1
t2
Feedback
t
High (Stable but not reponsive)
Low (Responsive)
High (Stable)

**Supplementary Figure S9. Oscillating feedback facilitates robust, cue-responsive polarization from diverse initial states in a mathematically tractable one species polarity model.**

**a**, Left, schematic showing a simplified one-species polarity model based on the wave-pinning model developed by Mori et al. [105]. Here, rather than using double negative feedback between aPARs and pPARs to polarize, the system relies on positive feedback within the polarity species X for the local recruitment of X from the cytoplasm by existing membrane-associated X. Within specific parameter ranges, as noted in Trong et al. [52], 4 possible steady states exist as with the PAR model, (i) uniform X low, (ii) uniform X high, (iii) polarised with X high at posterior but low at anterior, and (iv) polarised with X high at anterior but low at posterior. These states can all be captured in a phase space varying the concentration of X in the anterior, $X_a$, or X in the posterior, $X_p$. Cue is modelled as a local increase in on rate at the anterior of the cell, similar to previous work [54,105]. Quivers represent movement across the system state space, and the colors represent the quasi-potential of the system, calculated using the Fokker-Planck equation (see Modelling Supplement).

**b,** Same as **a,** but for low levels of positive feedback.

**c,** Schematic illustrating the full simulation of polarization related to **d**, intending to polarize systems from state i, ii and iii to state iv using the same cue described above.

**d**, Oscillatory feedback allows stable yet adaptable polarization from all system states in the landscape. Left, in the constant high feedback condition, only points around state i are able to respond reliably to cues and move beyond the transition point for polarization towards state iv. Right, in the oscillatory feedback condition, temporary low feedback facilitates cue-responsiveness towards the transition point for all initial states, before increasing feedback locks in the stable polarized state. Dotted black lines represent the transition point of the system, converging towards the basin of attraction representing state iv.

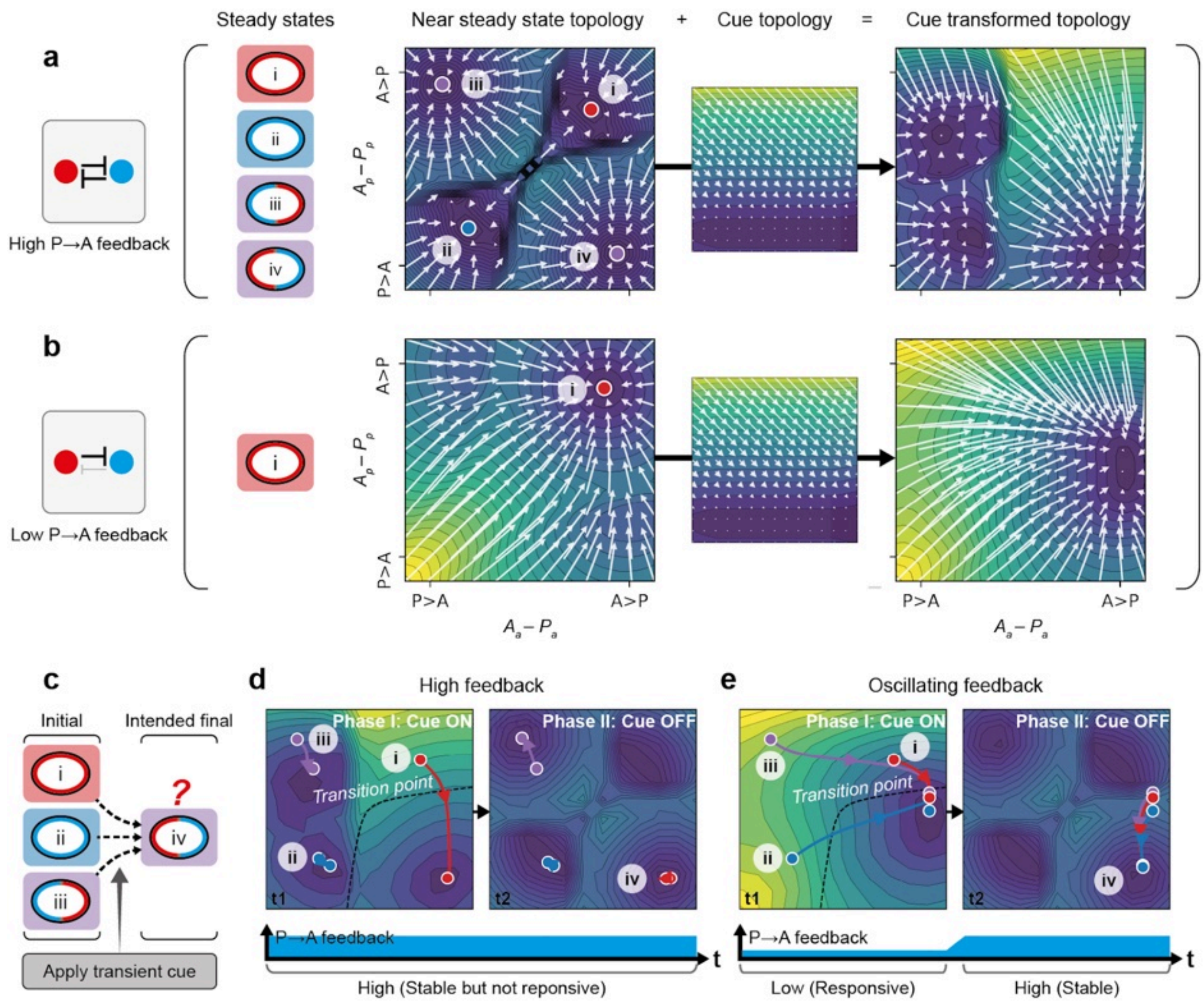
a
High P→A feedback
Steady states
Near steady state topology
+
Cue topology
=
Cue transformed topology
i
ii
iii
iv
A>P
P>A
b
Low P→A feedback
P>A
A>P
c
Initial
Intended final
?
Apply transient cue
d
High feedback
Phase I: Cue ON
Phase II: Cue OFF
Transition point
t1
t2
P→A feedback
t
High (Stable but not reponsive)
e
Oscillating feedback
Phase I: Cue ON
Phase II: Cue OFF
Transition point
P→A feedback
Low (Responsive)
High (Stable)

**Supplementary Figure S10. Oscillating feedback facilitates robust, cue-responsive polarization from diverse initial states in a full PDE PAR model.**

**a-e** Identical to Figure 3, but using a system described with partial differential equations instead. Note that the colors of the contours here do not represent the quasi-potential of the system, but indicate the velocity of the arrows instead.

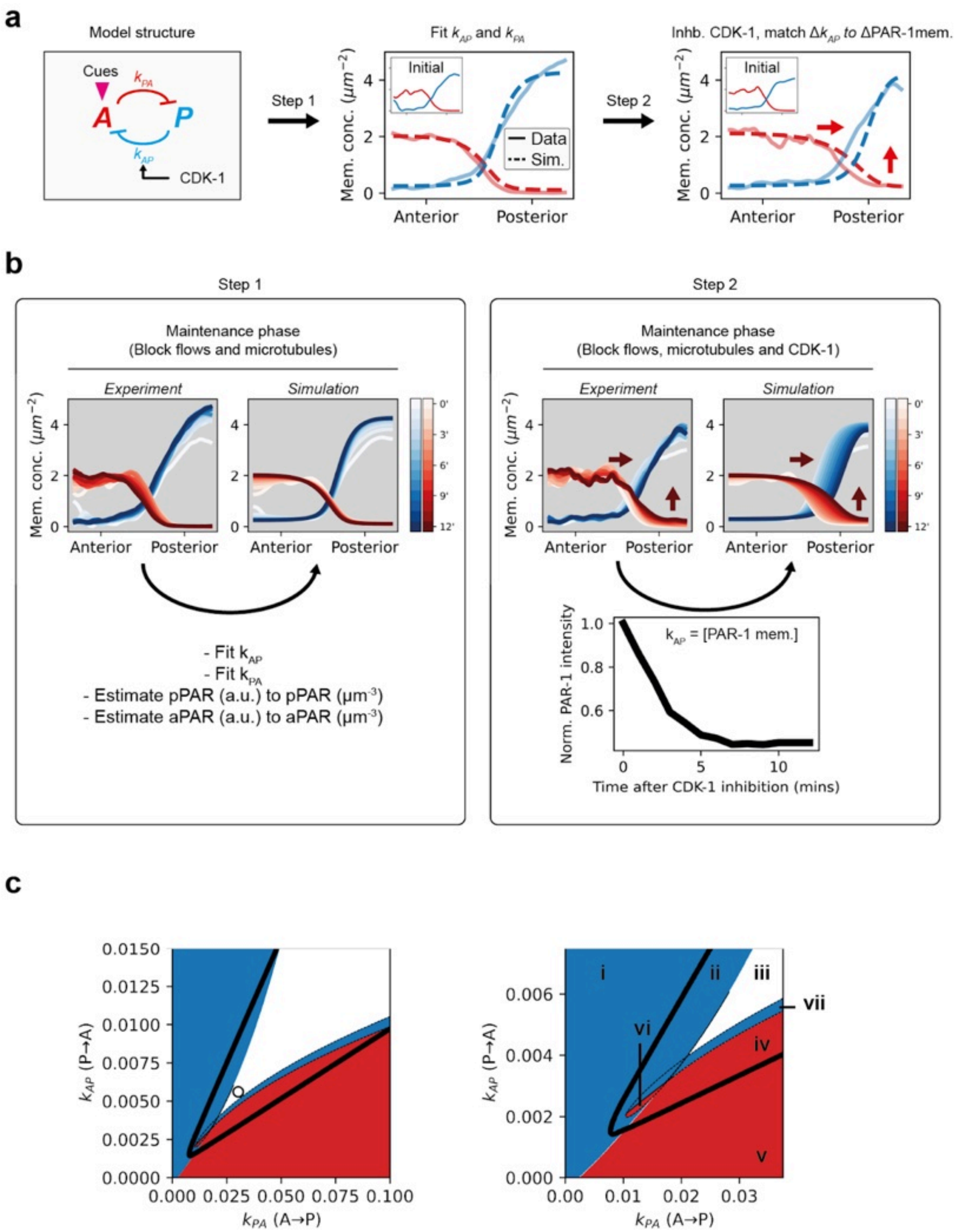
a
Model structure
Cues
A
P
CDK-1
Step 1
Fit $k_{AP}$ and $k_{PA}$
Initial
Data
Sim.
Mem. conc. ($\mu m^{-2}$)
Anterior
Posterior
Step 2
Inhb. CDK-1, match $\Delta k_{AP}$ to ΔPAR-1mem.
b
Step 1
Maintenance phase
(Block flows and microtubules)
Experiment
Simulation
- Fit $k_{AP}$
- Fit $k_{PA}$
- Estimate pPAR (a.u.) to pPAR ($\mu m^{-3}$)
- Estimate aPAR (a.u.) to aPAR ($\mu m^{-3}$)
Step 2
Maintenance phase
(Block flows, microtubules and CDK-1)
$k_{AP}$ = [PAR-1 mem.]
Norm. PAR-1 intensity
Time after CDK-1 inhibition (mins)
c
$k_{AP}$ (P→A)
$k_{PA}$ (A→P)
i
ii
iii
iv
v
vi
vii

**Supplementary Figure S11. Estimation of *C. elegans* $k_{AP}$ (P→A) and $k_{PA}$ (A→P) feedback via fitting with experiments.**

**a,** Schematic illustrating the strategy used for estimating dynamic antagonistic feedback rates in changing CDK-1 activity. Step 1, antagonistic feedback was first obtained by fitting a simplified model structure with measured diffusion/ on/ off rates (see Modelling Supplement) to data roughly around pronuclear centration, when CDK-1 activity is thought to be high. Step 2, estimating how pPAR to aPAR antagonistic feedback ($k_{AP}$) changes during CDK-1 inhibition. We approximated changing $k_{AP}$ activity as changes in PAR-1 membrane levels following CDK-1 inhibition, and saw that the PAR-2 and PAR-6 membrane profiles fit well with the simulations.

**b,** Comparison of simulations and data when fitting Step 1 and Step 2 shown in **a**. Changing colors of membrane line profiles reflect time relative to drug addiction.

**c,** Parameter-space topology of fitted parameters of the PAR model. Blue region (i, ii) supports pPAR dominant homogeneous states. Red region (iv, v) supports aPAR dominant homogenous states. White region (iii) supports both aPAR and pPAR dominant homogenous states, depending on the initial state. Dotted blue region (vi) indicates region that can undergo spontaneous symmetry breaking if beginning from pPAR initial states. Dotted red region (vii) indicates the same, but beginning from aPAR initial states. Solid black lines indicate regions permissible to stable polarization. White dotted circle represents where the fitted parameters lie.

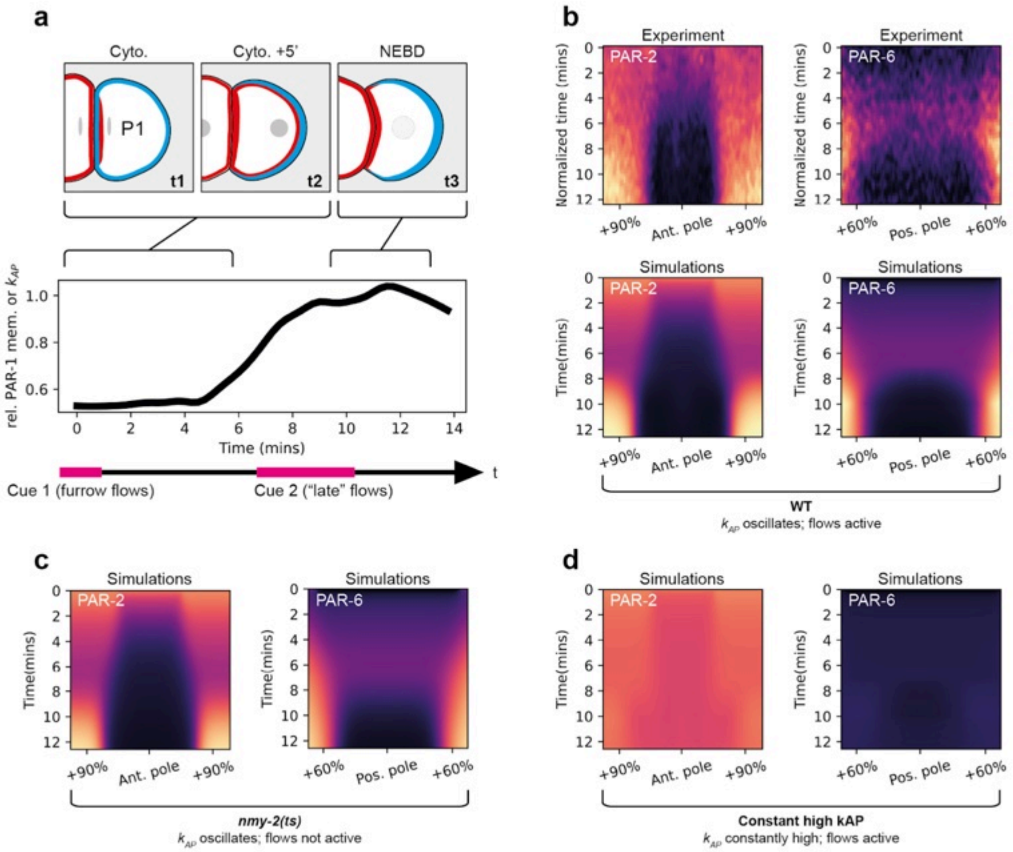
a
Cyto.
Cyto. +5'
NEBD
P1
t1
t2
t3
rel. PAR-1 mem. or kAP
Time (mins)
Cue 1 (furrow flows)
Cue 2 ("late" flows)
t
b
Experiment
PAR-2
PAR-6
Normalized time (mins)
Simulations
Time(mins)
+90%
Ant. pole
+60%
Pos. pole
WT
kAP oscillates; flows active
c
nmy-2(ts)
kAP oscillates; flows not active
d
Constant high kAP
kAP constantly high; flows active

**Supplementary Figure S12. Simulations of a modified PAR model representing P1 polarization.**

**a,** Top, schematic illustrating P1 polarization pattern through the cell cycle. Bottom, estimated $k_{AP}$ used for simulations, which is defined by relative PAR-1 membrane levels through the cell cycle.

**b,** Top, average spatiotemporal profiles of PAR-2 and PAR-6 during P1 polarization of wildtype embryos, data from Ng et al., 2023 [11]. Bottom, simulations of a P1-specific model closely resembles the experimental data.

**c,** Simulations of a P1-specific model shows that "late" flows observed in P1 are not required for polarization, matching previous experiments [11,12].

**d,** Simulations of a P1-specific model shows that oscillatory feedback in P1 is important for polarization, as a system with constant high feedback cannot polarize.

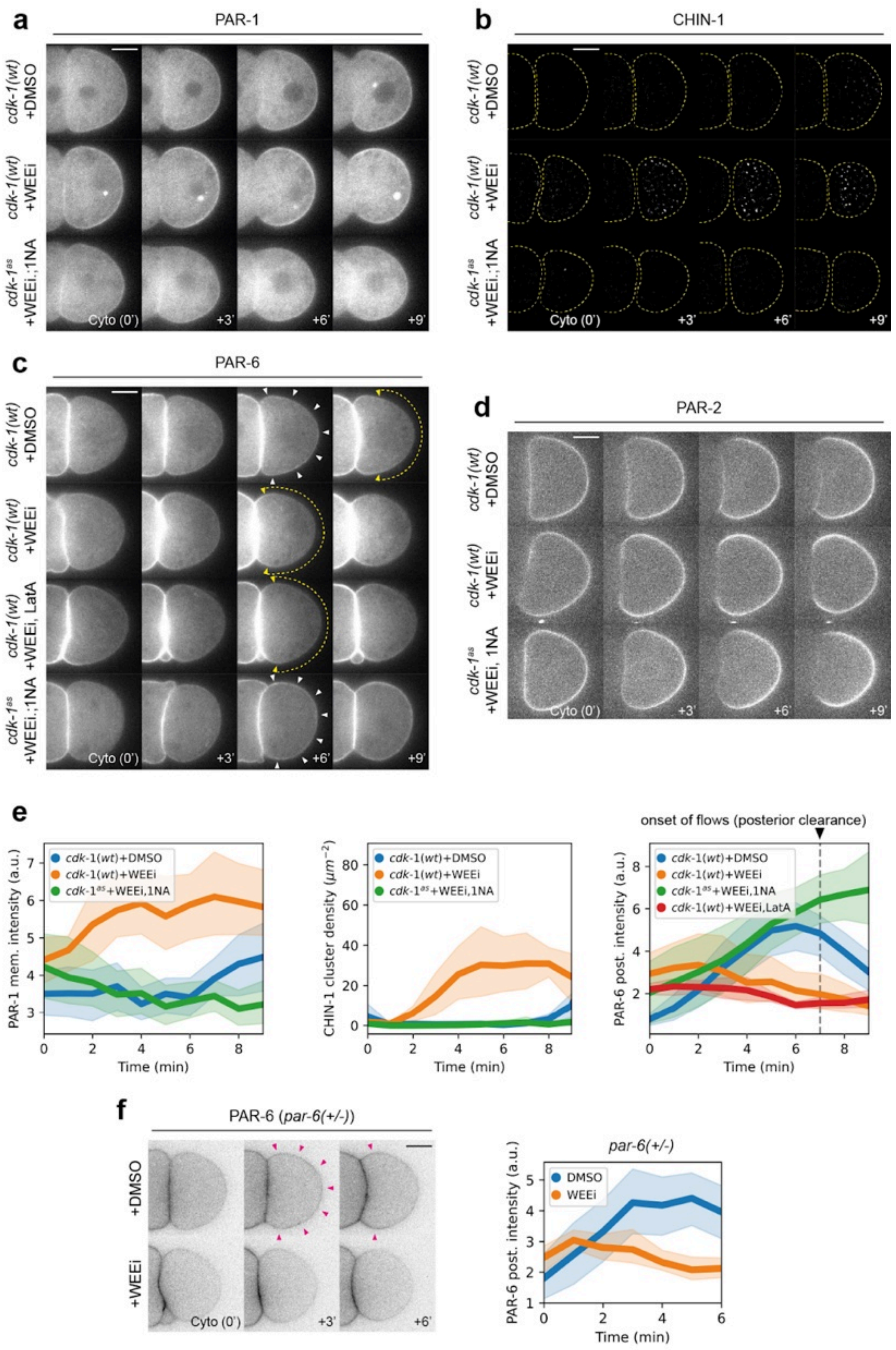
a
PAR-1
b
CHIN-1
cdk-1(wt) +DMSO
cdk-1(wt) +WEEi
cdk-1as +WEEi.;1NA
Cyto (0')
+3'
+6'
+9'
c
PAR-6
cdk-1(wt) +WEEi, LatA
d
PAR-2
cdk-1as +WEEi, 1NA
e
cdk-1(wt)+DMSO
cdk-1(wt)+WEEi
cdk-1as+WEEi,1NA
cdk-1(wt)+WEEi,LatA
PAR-1 mem. intensity (a.u.)
CHIN-1 cluster density (μm−2)
PAR-6 post. intensity (a.u.)
Time (min)
onset of flows (posterior clearance)
f
PAR-6 (par-6(+/-))
+DMSO
+WEEi
par-6(+/-)
DMSO
WEEi

**Supplementary Figure S13. Effects of WEE-1 inhibition on PAR proteins during P1 polarization.**

**a,** WEE-1 inhibition accelerates PAR-1 membrane loading. Time series of midplane confocal images of embryos expressing PAR-1::GFP and mCherry::PAR-2 (not shown) in either a *cdk-1(wt)* (NWG0332) or *cdk-1$^{as}$* (NWG0566) background, acutely treated with DMSO, 20µM WEEi (PD0166825) or 20µM 1NA-PP1. Sample sizes: *cdk-1(wt)* + DMSO (n=5), *cdk-1(wt)* + WEEi (n=6), *cdk-1$^{as}$* + WEEi + 1NA-PP1 (n=8).

**b,** WEE-1 inhibition accelerates CHIN-1 membrane loading. Time series of background subtracted CHIN-1 cortical images in embryos expressing mNG::CHIN-1 in either a *cdk-1(wt)* (NWG0451) or *cdk-1$^{as}$* (NWG518) background, acutely treated with DMSO, 20µM WEEi or 20µM 1NA-PP1. Sample sizes: *cdk-1(wt)* + DMSO (n=5), *cdk-1(wt)* + WEEi (n=6), *cdk-1$^{as}$* + WEEi + 1NA-PP1 (n=4).

**c,** WEE-1 inhibition reduces PAR-6 membrane loading onto the posterior membrane. Same setup as (A) but for PAR-6. Embryos expressing PAR-6::mNG were used, in either a *cdk-1(wt)* (LP216) or *cdk-1$^{as}$* (NWG0559) background, acutely treated with DMSO, 20µM WEEi, 20µM 1NA-PP1 or 0.5 µM Latrunculin A. Sample sizes: *cdk-1(wt)* + DMSO (n=7), *cdk-1(wt)* + WEEi (n=8), *cdk-1(wt)* + WEEi + Latrunculin A (n=4), or *cdk-1$^{as}$* + WEEi + 1NA-PP1 (n=5).

**d,** WEE-1 inhibition has modest effects on PAR-2 polarization. Same experiments as (A) but showing mCherry::PAR-2 instead.

**e,** Quantification of average membrane levels for conditions corresponding to **a-c**. Note that for PAR-6, only the posterior membrane outside of the contact site was considered for quantification.

**f,** Left, a time series of midsection confocal images of embryos expressing PAR-6::mNG in *par-6* heterozygous conditions (*par-6::mNG/–*) (NWG0141 x LP216), acutely treated with either DMSO (n=10) or 20µM WEEi (n=8). Magenta arrowheads indicate significantly higher levels of PAR-6 posterior membrane loading in DMSO treated embryos compared to WEEi treated embryos. Right, the corresponding quantification of average PAR-6 membrane levels at the posterior region of P1, outside of the contact site, after cell birth.

Mean and 95% confidence interval (bootstrapped) indicated. Scale bars, 10µm.

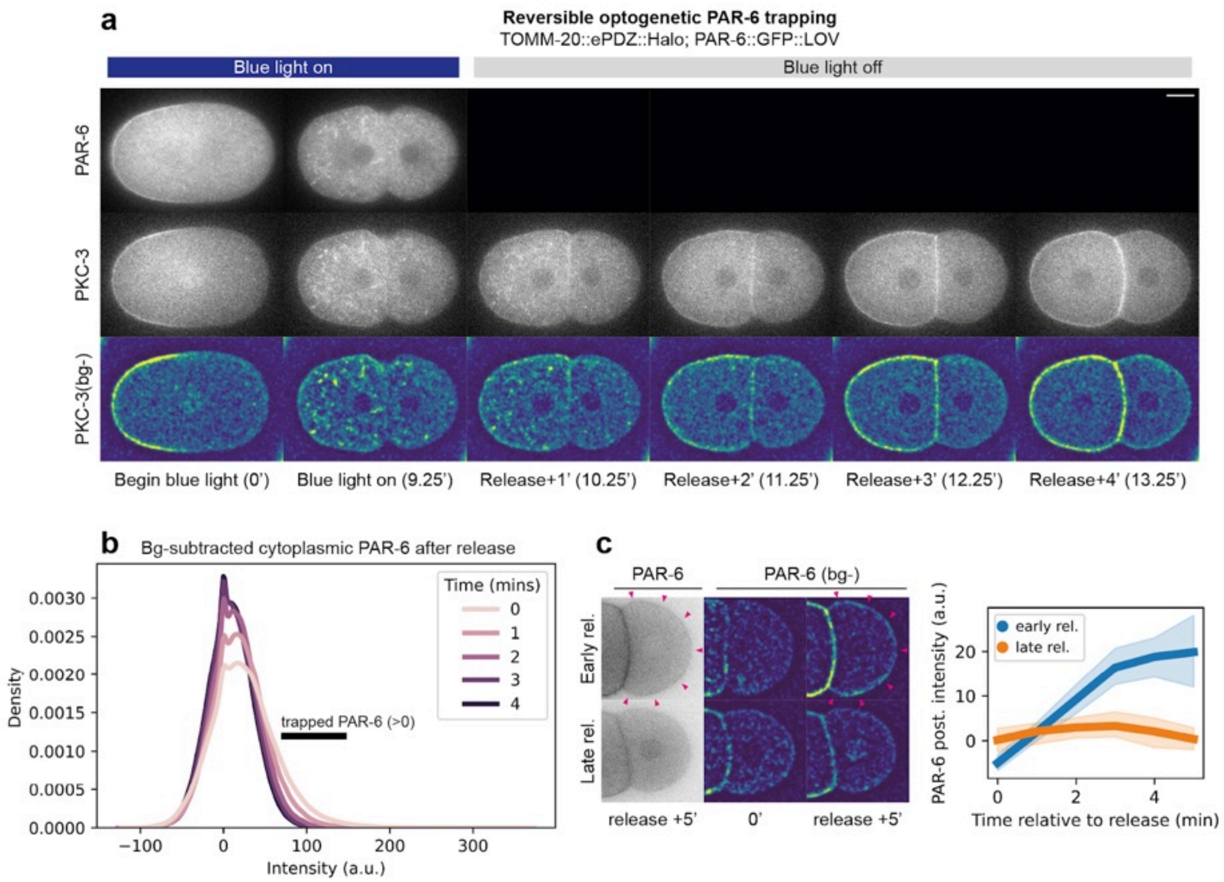
a
Reversible optogenetic PAR-6 trapping
TOMM-20::ePDZ::Halo; PAR-6::GFP::LOV
Blue light on
Blue light off
PAR-6
PKC-3
PKC-3(bg-)
Begin blue light (0')
Blue light on (9.25')
Release+1' (10.25')
Release+2' (11.25')
Release+3' (12.25')
Release+4' (13.25')
b
Bg-subtracted cytoplasmic PAR-6 after release
Time (mins)
0
1
2
3
4
trapped PAR-6 (>0)
Density
0.0030
0.0025
0.0020
0.0015
0.0010
0.0005
0.0000
−100
0
100
200
300
Intensity (a.u.)
c
PAR-6
PAR-6 (bg-)
Early rel.
Late rel.
release +5'
0'
release +5'
early rel.
late rel.
PAR-6 post. intensity (a.u.)
20
10
0
0
2
4
Time relative to release (min)

**Supplementary Figure S14. Dynamics of reversible optogenetic knocksideways of PAR-6.**

**a,** Time series of midplane confocal images of a representative embryo expressing PAR-6::GFP::LOV, TOMM-20::ePDZ::Halo and mScarlet-I::PKC-3 (NWG0630) subjected to blue light illumination and removal. Note that upon blue light illumination, PAR-6 membrane levels reduce and become sequestered into cytoplasmic structures (presumably mitochondria). This process is reversible by relieving blue light illumination.

**b,** Quantification of PAR-6 release dynamics following removal of blue light illumination. Histogram of fluorescence distribution after background subtraction of AB cytoplasm is shown. PAR-6 sequestered into the mitochondria will appear as fluorescence distributions larger than 0 after background removal. In approximately 2 minutes, PAR-6 fluorescence distribution collapses to a distribution centered around 0, suggesting that PAR-6 release from mitochondria occurs in around the same time frame following blue light removal.

**c,** Left, a time series of midsection confocal images of embryos expressing PAR-6::GFP::LOV, TOMM-20::ePDZ::Halo and mScarlet-I::PKC-3 (NWG0630), with mitochondria-trapped PAR-6 release early (~3mins) (n=6) or late (~7.5mins) (n=6) after cell birth. Magenta arrowheads indicate significantly higher levels of PAR-6 posterior membrane loading in early release (~3 mins after cell birth) embryos compared to late release (~7 mins after cell birth) embryos. Right, the corresponding quantification of average PAR-6 membrane levels at the posterior region of P1, outside of the contact site, after cell birth.

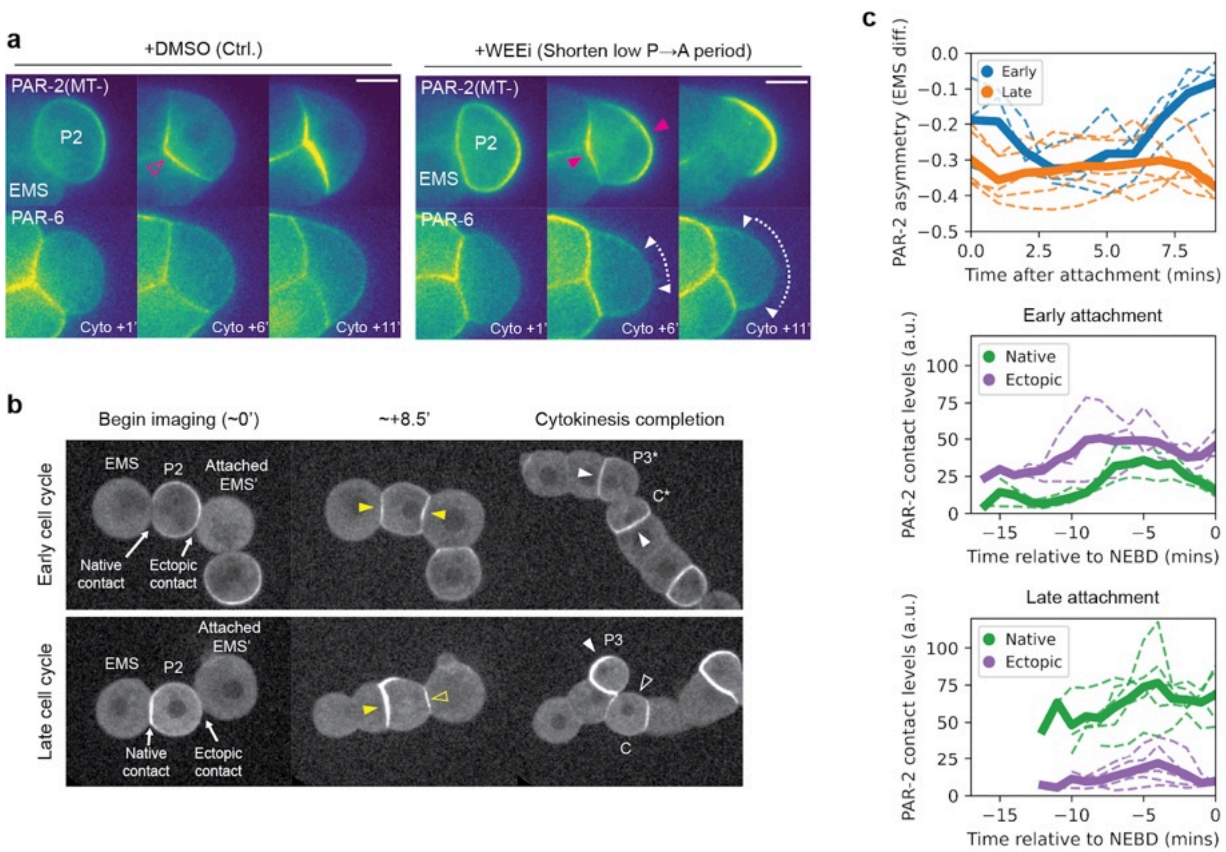
a
+DMSO (Ctrl.)
+WEEi (Shorten low P→A period)
PAR-2(MT-)
P2
EMS
PAR-6
Cyto +1'
Cyto +6'
Cyto +11'
b
Begin imaging (~0')
~+8.5'
Cytokinesis completion
Early cell cycle
Late cell cycle
EMS
P2
Attached EMS'
Native contact
Ectopic contact
P3*
C*
P3
C
c
PAR-2 asymmetry (EMS diff.)
Early
Late
Time after attachment (mins)
Early attachment
Late attachment
Native
Ectopic
PAR-2 contact levels (a.u.)
Time relative to NEBD (mins)

**Supplementary Figure S15. Additional information on cue-sensitivity of P2 to EMS signalling when decoupled with wild type cell cycle progression.**

**a,** Schematic showing PAR-2(MT-) and PAR-6 behavior when P2 blastomeres were treated with the WEE-1 inhibitor. Same samples as shown in Fig. **5c**, but showing PAR-6::mScarlet-I as well. Magenta arrowheads indicate differences in the number of polarity domains in DMSO or WEEi treated embryos, when PAR-2 is recruited to the contact, presumably by the onset of signaling from EMS. White arrowheads with dotted lines indicate posterior PAR-6 clearance by the ectopic domain in WEEi treated conditions.

**b,** Uncropped images of embryo attachment experiments, traced until after cell division. Yellow closed arrowheads indicate the presence of PAR-2(MT-) domains, while yellow open arrowheads indicate a second smaller PAR-2(MT-) domain. Note that PAR-2 asymmetry is greater between the P2 daughter cells (P3 and C) of late attachment embryos than early.

**c,** Quantification of conditions corresponding to **b**. Mean (bold lines) and results of individual experiments (dotted lines) shown.

Scale bars, 20µm.

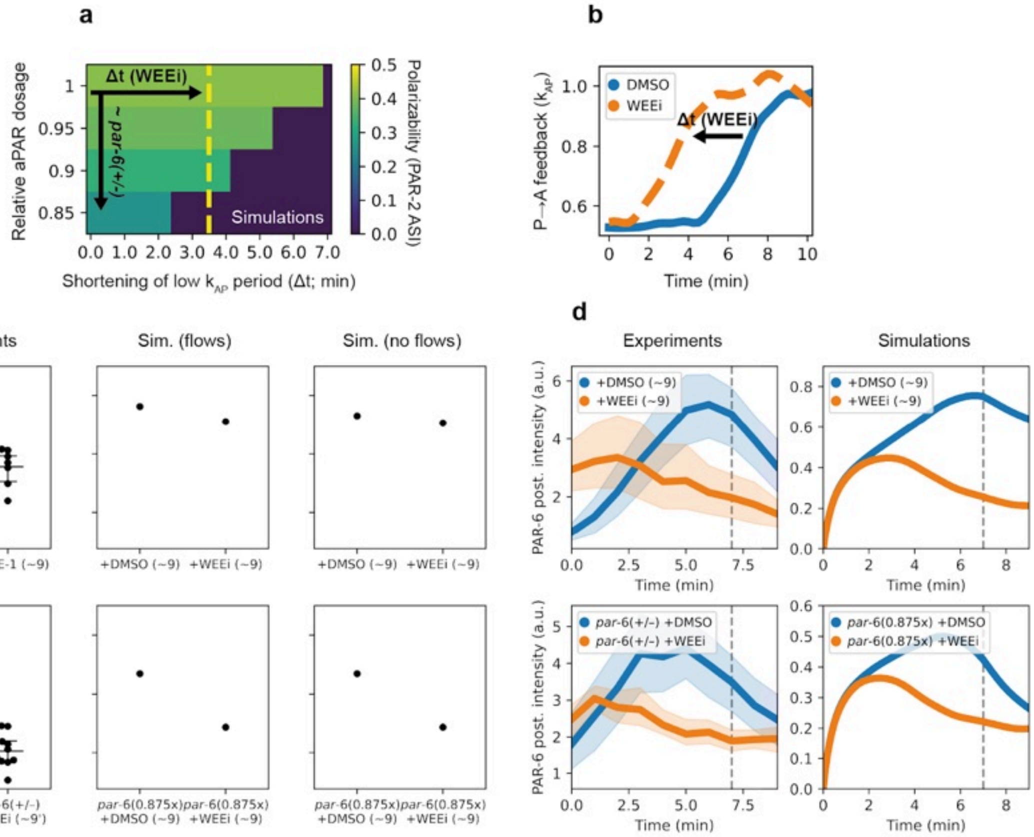
a
Δt (WEEi)
~ par-6(+/-)
Simulations
Relative aPAR dosage
Shortening of low $k_{AP}$ period (Δt; min)
Polarizability (PAR-2 ASI)
b
DMSO
WEEi
Δt (WEEi)
P→A feedback ($k_{AP}$)
Time (min)
c
Experiments
Sim. (flows)
Sim. (no flows)
PAR-2 ASI
+DMSO (~9) +WEE-1 (~9)
+DMSO (~9) +WEEi (~9)
par-6(+/-) +DMSO (~9')
par-6(+/-) +WEEi (~9')
par-6(0.875x) +DMSO (~9)
par-6(0.875x) +WEEi (~9)
d
Experiments
Simulations
+DMSO (~9)
+WEEi (~9)
par-6(+/-) +DMSO
par-6(+/-) +WEEi
par-6(0.875x) +DMSO
par-6(0.875x) +WEEi
PAR-6 post. intensity (a.u.)
Time (min)

**Supplementary Figure S16. Simulation outcomes resemble P1 polarization results from WEE-1 inhibition experiments.**

**a,** Simulations of P1 polarizability when the period of low $k_{AP}$ is shortened and in changing aPAR levels. Yellow dotted line indicates the estimated shortening of the low $k_{AP}$ period when P1 blastomeres are treated with a WEE-1 inhibitor (PD0166825 or simply WEEi; see right and Supplementary Figure **S13**).

**b,** Left, WEE-1 inhibition should lead to precocious activation of CDK-1, leading to a corresponding increase in pPAR to aPAR antagonistic feedback.

**c,** Predicted dynamics of pPAR to aPAR antagonistic feedback when WEE-1 is inhibited compared to wildtype (see Supplementary Figure **S13**).

**d,** A comparison between experimental results and theory for PAR-2 asymmetry (ASI) ~9 mins after cytokinesis completion/ birth of cell, when WEE-1 is inhibited. The simulations were performed in the presence or absence of “late” flows described in Supplementary Figure **S12a**. We chose to compare PAR-6 heterozygotes with 0.875x total dosage in the models, as we noted that polarity was already disrupted at this level of dosage when combined with a shortened low feedback period.

**e,** A comparison between experimental results and theory for PAR-6 membrane levels following WEE-1 inhibition.

Dotted lines indicate onset of “late” flows, which advects aPARs towards the embryo anterior.

Mean and 95% confidence interval (bootstrapped) indicated.

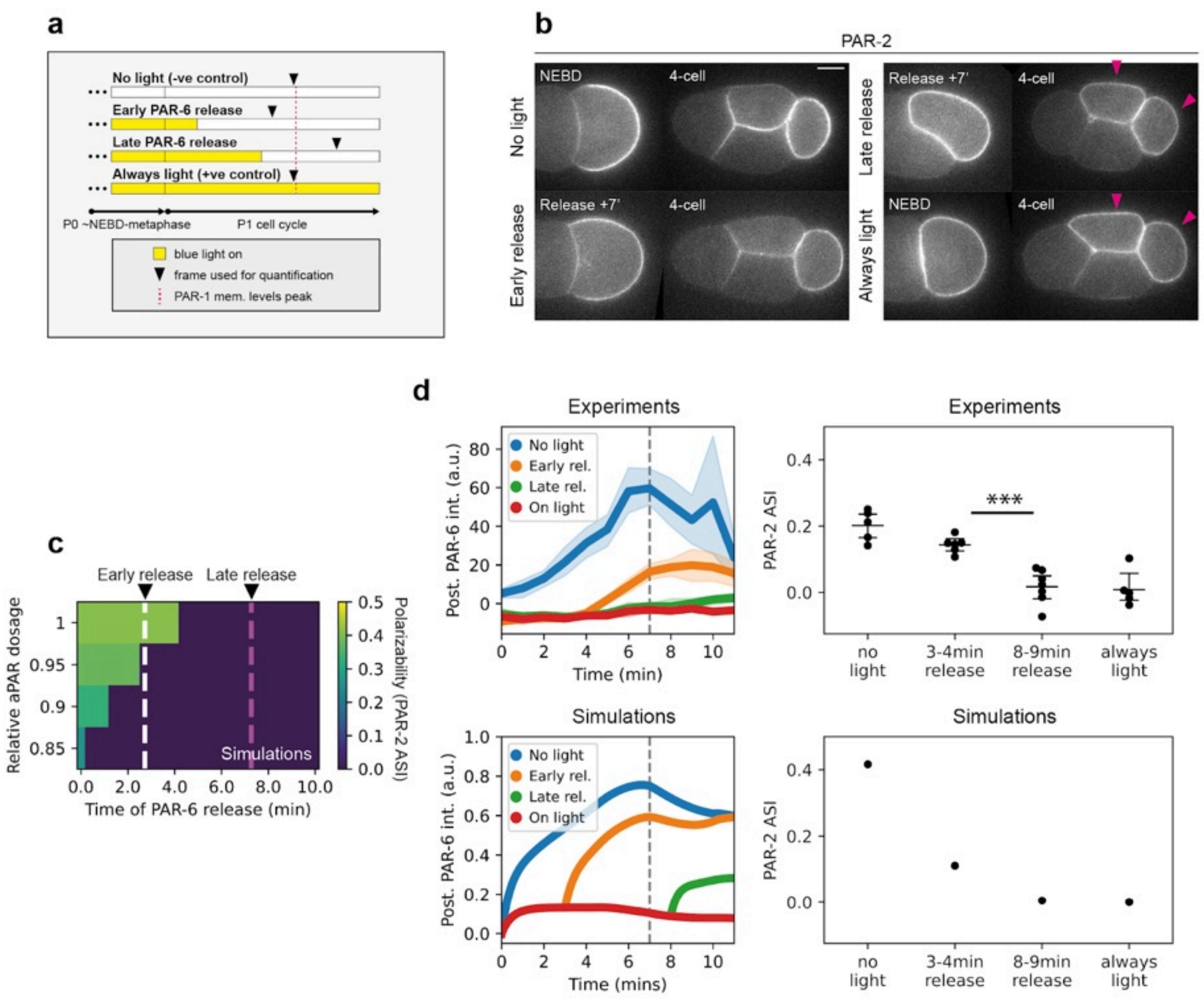
a
No light (-ve control)
Early PAR-6 release
Late PAR-6 release
Always light (+ve control)
P0 ~NEBD-metaphase
P1 cell cycle
blue light on
frame used for quantification
PAR-1 mem. levels peak
b
PAR-2
No light
NEBD
4-cell
Early release
Release +7'
4-cell
Late release
Release +7'
4-cell
Always light
NEBD
4-cell
c
Early release
Late release
Relative aPAR dosage
Polarizability (PAR-2 ASI)
Simulations
Time of PAR-6 release (min)
d
Experiments
Post. PAR-6 int. (a.u.)
No light
Early rel.
Late rel.
On light
Time (min)
Experiments
PAR-2 ASI
***
no light
3-4min release
8-9min release
always light
Simulations
Post. PAR-6 int. (a.u.)
No light
Early rel.
Late rel.
On light
Time (mins)
Simulations
PAR-2 ASI
no light
3-4min release
8-9min release
always light

**Supplementary Figure S17. Simulation outcomes resemble P1 polarization results from optogenetic knocksideways experiments.**

**a,** Schematic illustrating the timeline for optogenetic knocksideways experiments for PAR-6. Some embryos were either never or always illuminated with blue light to serve as negative and positive controls respectively.

**b,** Left, time series of midsection confocal images of 2- and 4-cell stage embryos expressing PAR-6::GFP::LOV, TOMM-20::ePDZ::Halo and GFP::PAR-2 (NWG0597) [109,110] subjected to experimental conditions corresponding to (C). Sample sizes: no light (n=5), early release (n=6), late release (n=7), always light (n=5). Magenta arrowheads indicate symmetric PAR-2 inheritance in 4-cell stage embryos. Right, quantifications of PAR-2 asymmetry index (ASI) for the corresponding conditions.

**c,** Top, simulations predict that P1 polarization should be sensitive to activation of polarity at different times, which can be achieved by “releasing” aPARs at different times. Bottom, simulations of P1 polarizability at different times of aPAR release and in changing aPAR levels. White and magenta dotted lines indicate release times which allow polarization or not respectively in wildtype aPAR levels, which is the timings we chose for experimentation.

**d,** Left, comparison of posterior PAR-6 membrane profiles in P1 (outside of contact region) between simulations and experimental results. Dotted lines indicate onset of “late” flows, which advects aPARs towards the embryo anterior. Right, comparison of PAR-2 asymmetry (ASI) between experimental results and theory.

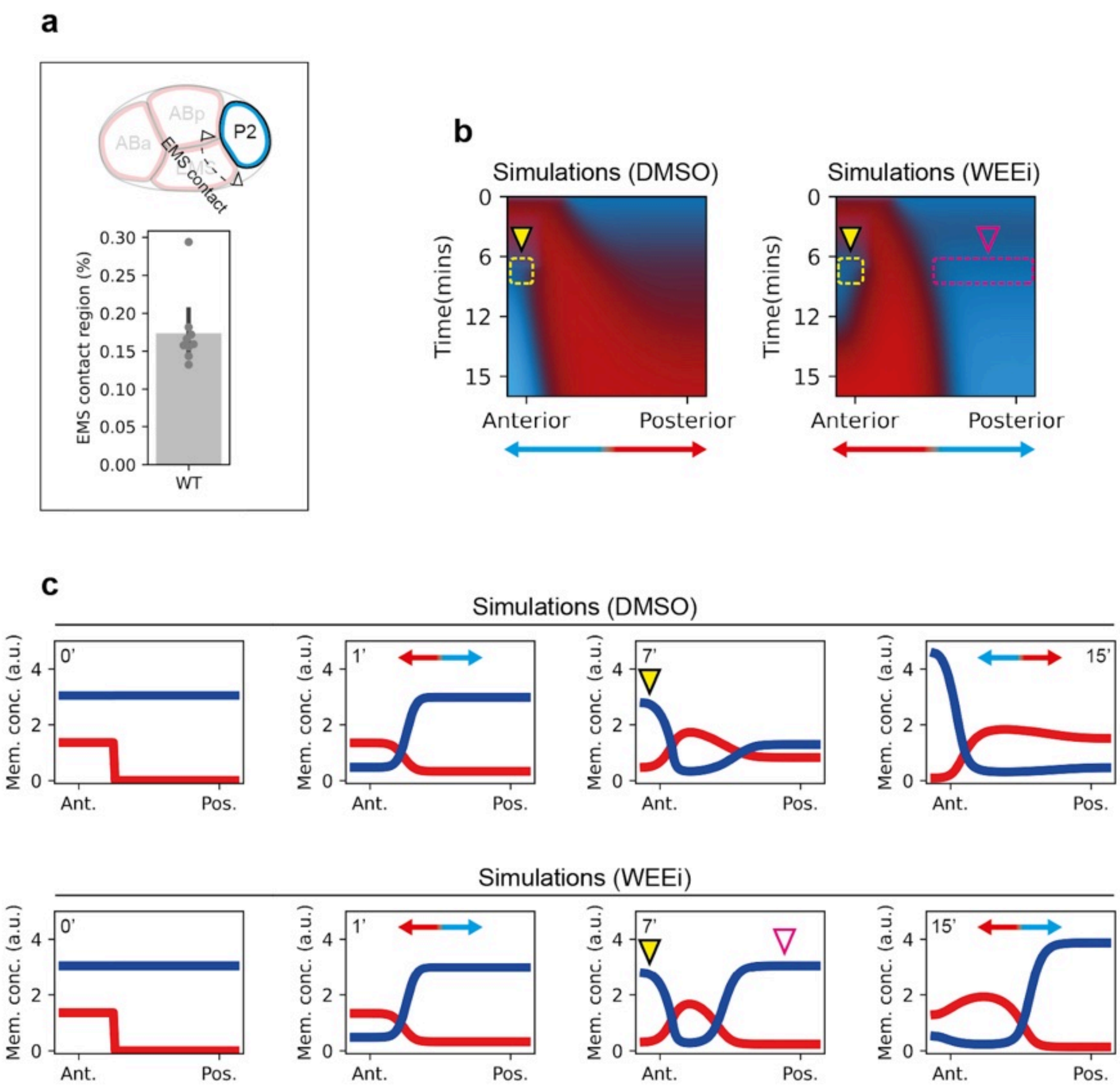
a
ABa
ABp
P2
EMS contact
EMS contact region (%)
0.30
0.25
0.20
0.15
0.10
0.05
0.00
WT
b
Simulations (DMSO)
Simulations (WEEi)
Time(mins)
0
6
12
15
Anterior
Posterior
c
Simulations (DMSO)
Simulations (WEEi)
Mem. conc. (a.u.)
0'
1'
7'
15'
4
2
0
Ant.
Pos.

**Supplementary Figure S18. Simulation outcomes resemble P2 polarity reversal results from WEE-1 inhibition experiments.**

**a,** Quantification of the fraction of EMS-P2 contact region, which is used to then simulate the fractional domain size of the EMS signalling cue.

**b,** Spatiotemporal membrane profiles of PAR-2(MT-) simulations in P2, subjected to DMSO or WEEi, which changes when pPAR to aPAR antagonistic feedback levels increase. PAR-2(MT-) mutation is captured by increasing pPAR to aPAR antagonistic feedback rates. Yellow arrowhead indicates the “correct” developmental cue originating from MES-1/SRC-1 signaling from EMS, magenta open arrowhead indicates ectopic polarity domain formed by furrow accumulation of PAR-3 at EMS contact. Blue arrow indicates orientation of the final pPAR domain.

**c,** Same as in (B) but showing membrane profiles of pPARs (blue) and aPARs (red) at indicated time points. Double sided arrow graded from red to blue indicates the orientation of the polarity axis.

### Supplementary Tables

**Table S1**. Parameters used for simplified PAR model

| Parameter | Species | |
|---|---|---|
| | A (aPAR) | P (pPAR) |
| L | 50 µm | |
| ψ | 0.5 $\mu m^{-1}$ | |
| $\rho_A = \rho_P$ | 1 $\mu m^{-3}$ | |
| D | 0.05 $\mu m^2\ s^{-1}$ | |
| $k_{off}$ | 0.005 $s^{-1}$ | |
| $k_{on}$ | 0.006 µm $s^{-1}$ | |
| $\alpha = \beta$ | 2 | |
| $k_{PA} = k_{AP}$ | 0.1 $\mu m^4\ s^{-1}$ | |
| $k_{cue}$ | 0.002 | |

**Table S2**. Parameters used for one-species polarity model based on the wave-pinning model

| Parameter | Species (X only) |
|---|---|
| L | 10 μm |
| $\psi$ | 1 $\mu m^{-1}$ |
| $\rho_X$ | 3.5 $\mu m^{-3}$ |
| D | 0.01 $\mu m^2 s^{-1}$ |
| $k_{off}$ | 1 $s^{-1}$ |
| $k_{on}$ | 0.001 $\mu m\ s^{-1}$ |
| n | 4 |
| $\gamma$ | 1.2 $\mu m\ s^{-1}$ |
| K | 1 $\mu m^{-2}$ |

Parameters are modified to capture 4 different polarity states instead of 3 in the original model by Mori et al. [105].

**Table S3**. Parameters used for *C. elegans*-specific PAR model

| Parameter | Species | |
|---|---|---|
| | A (aPAR) | P (pPAR) |
| L | 67.3 μm | |
| ψ | 0.174 $\mu m^{-1}$ | |
| $\rho_A$ / $\rho_P$ | $\rho_A$ = 1.56 $\mu m^{-3}$ | $\rho_P$ = 1 $\mu m^{-3}$ |
| D | 0.28 $\mu m^2\ s^{-1}$ | 0.15 $\mu m^2\ s^{-1}$ |
| $k_{off}$ | $5.4 \cdot 10^{-3}\ s^{-1}$ | $7.3 \cdot 10^{-3}\ s^{-1}$ |
| $k_{on}$ | $8.58 \cdot 10^{-3}\ \mu m\ s^{-1}$ | $4.74 \cdot 10^{-2}\ \mu m\ s^{-1}$ |
| α / β | 2 | |
| $k_{PA}$/ $k_{AP}$ | $k_{PA} = 3.03 \cdot 10^{-2}\ \mu m^4\ s^{-1}$ | $k_{AP} = 5.61 \cdot 10^{-3}\ \mu m^4\ s^{-1}$ |

All parameters used here are taken from [8] except from $k_{PA}$ and $k_{AP}$, which were fitted with data from Figure 2C.

### Supplementary Videos

**Supplementary Video 1**
PAR-1 membrane dynamics respond to changing CDK-1 activity. Midsection confocal video of an embryo expressing endogenous PAR-1::GFP in a *cdk-1$^{as}$* background, acutely treated with 1NA-PP1, followed by wash out of the drug.

**Supplementary Video 2**
CHIN-1 membrane dynamics respond to changing CDK-1 activity. HiLo video of an embryo expressing endogenous mNG::CHIN-1 in a *cdk-1$^{as}$* background, acutely treated with 1NA-PP1, followed by wash out of the drug.

**Supplementary Video 3**
PAR-2 and PAR-6 membrane dynamics after inhibition of CDK-1 and disruption of cytoskeleton. Midsection confocal video of an embryo expressing endogenous mCherry::PAR-2 and PAR-6::mNG in a *cdk-1$^{as}$* background, acutely treated with 1NA-PP1, Latrunculin A and Nocodazole.

**Supplementary Video 4**
Cue-induced polarization dynamics in a simplified PAR model with constant high feedback. Simulation output reflecting cue-induced polarization beginning from (i) homogenous aPAR high state (red circle), (ii) homogenous pPAR high state (blue circle), and (iii) PA-polarized state with aPARs high at posterior and pPARs high at anterior (posterior circle), towards (iv) AP-polarized state with aPARs high at anterior and pPARs high at posterior. Note that only homogenous aPAR high states can be effectively directed towards the AP-polarized state.

**Supplementary Video 5**
Cue-induced polarization dynamics in a simplified PAR model with oscillatory pPAR→aPAR feedback. Simulation output reflecting cue-induced polarization beginning from (i) homogenous aPAR high state (red circle), (ii) homogenous pPAR high state (blue circle), and (iii) PA-polarized state with aPARs high at posterior and pPARs high at anterior (posterior circle), towards (iv) AP-polarized state with aPARs high at anterior and pPARs high at posterior. Compared to the constant high feedback system, all states can be effectively directed towards the AP-polarized state.

**Supplementary Video 6**
PKC-3 dynamics during optogenetic knocksideways of PAR-6. Midsection confocal movie of an embryo expressing PAR-6::GFP::LOV, TOMM-20::ePDZ::Halo and mScarlet-I::PKC-3 subject to acute blue light illumination and removal. Note that PAR-6 and PKC-3 appear to aggregate in the cytoplasm upon illumination, suggesting sequestration to the mitochondria.

**Supplementary Video 7**

Spatiotemporal dynamics of PAR-2 in P2 under WEE-1 inhibition. Midsection confocal movie of an embryo expressing GFP::PAR-2(MT-; microtubule binding defective mutant) treated with either DMSO or PD0166825 (WEE-1 inhibitor; WEEi).

### MODELLING SUPPLEMENT

Ng et al., 2025

#### 1 Constructing phase portrait of PAR behaviors

Feedback oscillations have been proposed as a mechanism to balance sensitivity and stability in cell polarity networks. To investigate whether CDK-1–coupled oscillations within the PAR network can achieve this balance, we developed a modeling framework that illustrates the system's behavior as feedback levels vary. Specifically, we constructed a phase portrait of polarity behaviors based on a prototypical reaction-diffusion model of PAR polarity (Fig. 3) [1, 2, 3, 4, 5], which allowed us to analyze the system's dynamics as a landscape under different feedback strengths. Our approach was inspired by the work of Nandan and colleagues [6], who used a similar framework to describe the excitability and stability of the Cdc42 wave-pinning polarity model in yeast.

In the case of the Cdc42 wave-pinning model by Nandan et al., the x-axis and y-axis of the graph are described by the amount of membrane-associated Cdc42 at the left of the cell ($u_L$) and at the right of the cell ($u_R$) respectively. This representation allowed all possible states of the system to be captured: high levels of homogeneous Cdc42 on the membrane would be on the top right of the graph, Cdc42 uniformly off the membrane would be at the bottom left, Cdc42 polarized towards the left of the cell would be at the bottom right, and Cdc42 polarized towards the right of the cell would be at the top left.

PAR polarity involves two polarity species, i.e. aPARs and pPARs, instead of one. Thus, to allow visualization of different polarity behaviors using this approach, we used a functional representation of the axes instead. Specifically, the x-axis and y-axis are defined by the concentration difference between aPARs and pPARs at the anterior membrane ($A_a - P_a$) and at the posterior membrane ($A_p - P_p$). This should allow us to capture all possible polarity configurations defined in previous work by Goehring et al. [1], which includes two polarized states and two unpolarized states. Here, aPAR homogenous high states would have high $A_a > P_a$ and $A_p > P_p$ and would thus be on the top right corner of the graph. pPAR homogenous high states would be at the bottom left for the same reason, as $P_a > A_a$ and $P_p > A_p$ . Polarized states with aPARs high at the anterior and pPARs high at the posterior would be located at the bottom right ($A_a > P_a$ and $P_p > A_p$), while the opposite configuration of polarized state would be at the top left ($P_a > A_a$ and $A_p > P_p$).

We wanted to take advantage of this framework to analyze system sensitivity and stability due to its ability to capture a diverse range of polarity phases. In a sensitive system, cues should be able to direct polarity coherently across the landscape; in a stable system, polarized configurations should exist and be stably maintained.

#### 1.1 A simplified ODE PAR model

A theoretical model of PAR polarization was previously described using the following set of partial differential equations (PDEs) with zero-flux Neumann boundary conditions:

$$\begin{aligned}\partial_t A &= D_A \partial_x^2 A + k_{on,A} A_{cyto} - k_{off,A} A - k_{AP} P^\alpha A \\ \partial_t P &= D_P \partial_x^2 P + k_{on,P} P_{cyto} - k_{off,P} P - k_{PA} A^\beta P\end{aligned} \tag{1}$$

where $k_{AP}$ and $k_{PA}$ refers to pPAR to aPAR antagonistic feedback and aPAR to pPAR antagonistic feedback respectively, and $\alpha$ and $\beta$ refer to non-linear antagonism terms. Assuming fast cytoplasmic diffusion and mass conservation, the equations governing the cytoplasmic pool can be described as:

$$\begin{aligned}A_{cyto} &= \rho_A - \psi \bar{A} \\ P_{cyto} &= \rho_P - \psi \bar{P}\end{aligned} \tag{2}$$

where $\rho_A$ and $\rho_P$ describe total aPAR and pPAR concentrations respectively, $\psi$ describes the surface-area-to-volume ratio, and $\bar{A}$ and $\bar{P}$ represent average aPAR and pPAR concentrations on the membrane respectively.

For convenience, we simplified this reaction-diffusion model by discretizing the system into two compartments, representing the whole of the anterior and posterior domains respectively. For instance, aPARs at the anterior half of the cell are represented as a single species $A_a$, and aPARs at the posterior half of the cell as $A_p$. For simplicity, we symmetrized the system, such that diffusion, on and off rates, and the total concentration of $A$ and $P$ proteins are identical. Finally, we also modified the diffusion terms accordingly, to represent exchange between the anterior and posterior compartments of the cell. Together, this gives us the following four ordinary differential equations (ODEs):

$$\begin{aligned}\frac{\mathrm{d}A_a}{\mathrm{d}t} &= \tilde{D}(A_p - A_a) + k_{on} A_{cyto} - k_{off} A_a - k_{AP} P_a^\alpha A_a \\ \frac{\mathrm{d}A_p}{\mathrm{d}t} &= \tilde{D}(A_a - A_p) + k_{on} A_{cyto} - k_{off} A_p - k_{AP} P_p^\alpha A_p \\ \frac{\mathrm{d}P_a}{\mathrm{d}t} &= \tilde{D}(P_p - P_a) + k_{on} P_{cyto} - k_{off} P_a - k_{PA} A_a^\beta P_a \\ \frac{\mathrm{d}P_p}{\mathrm{d}t} &= \tilde{D}(P_a - P_p) + k_{on} P_{cyto} - k_{off} P_p - k_{PA} A_p^\beta P_p\end{aligned} \tag{3}$$

where $\tilde{D}$ are diffusion-like terms. Parameters are provided in Table S1.

Descriptions of the cytoplasmic pool of aPARs and pPARs are formally written as:

$$\begin{aligned} A_{cyto} &= \rho_A - \psi(\frac{A_a + A_p}{2}) \\ P_{cyto} &= \rho_P - \psi(\frac{P_a + P_p}{2}) \end{aligned} \tag{4}$$

#### 1.2 Phase diagram in parameter space varying antagonism terms in the ODE model

To verify that the simplified two-step discretization model exhibited qualitative features similar to the full PDE model described previously [1, 2], we calculated the parameter space of the ODE system varying the antagonism terms $k_{AP}$ and $k_{PA}$ as in Goehring et al. and Trong et al. (Fig. S7) [1, 4].

Homogeneous steady states were calculated analytically. First we remove the spatial component from equation (3) and (4) for simplification, to get:

$$\begin{aligned} \frac{\mathrm{d}A}{\mathrm{d}t} &= k_{on}A_{cyto} - k_{off}A - k_{AP}P^{\alpha}A \\ \frac{\mathrm{d}P}{\mathrm{d}t} &= k_{on}P_{cyto} - k_{off}P - k_{PA}A^{\beta}P \\ A_{cyto} &= \rho_A - \psi A \\ P_{cyto} &= \rho_P - \psi P \end{aligned} \tag{5}$$

In equation (5), we first solve for $\frac{\mathrm{d}P}{\mathrm{d}t} = 0$ and rearrange to get:

$$P = \frac{k_{on}\rho_P}{k_{on}\psi + k_{off} + k_{PA}A^{\beta}} \tag{6}$$

and substituted this expression of $P$ into the first equation from (5), assuming $\frac{\mathrm{d}A}{\mathrm{d}t} = 0$, i.e.

$$0 = k_{on}A_{cyto} - k_{off}A - k_{AP}P^{\alpha}A \tag{7}$$

We then solved the values for $A$ and $P$.

We identified 3 possible steady states as in Goehring et al. [1]: aPAR dominant regions ($A > P$), pPAR dominant regions ($P > A$), and regions that can support both aPAR and pPAR dominant regions as well as a third unstable steady state where $A = P$.

Regions where homogeneous aPAR or pPAR dominant regions can spontaneously polarize were further identified by performing linear stability analysis considering the following ansatz:

$$\begin{aligned} U(t) &= U_0 + \delta U(t) \\ U &\in \{A_a, A_p, P_a, P_p\} \end{aligned} \tag{8}$$

where $\delta U = (\delta A_a, \delta A_p, \delta P_a, \delta P_p)^T \mathrm{e}^{\lambda t}$, representing a small perturbation with a growth rate $\lambda$, and $U_0$ reflects steady state distributions of PAR proteins. Substituting this into equation (3) and linearizing, we get the following Jacobian:

$$J = \begin{pmatrix} \frac{k_{on}A\psi}{2} + k_{off}A + k_{AP}P_0^{\alpha} & \frac{k_{on}A\psi}{2} & k_{AP}\alpha {P_0}^{\alpha-1}A_0 & 0 \\ \frac{k_{on}A\psi}{2} & \frac{k_{on}A\psi}{2} + k_{off}A + k_{AP}{P_0}^{\alpha} & 0 & k_{AP}\alpha {P_0}^{\alpha-1}A_0 \\ k_{PA}\beta {A_0}^{\beta-1}P_0 & 0 & \frac{k_{on}P\psi}{2} + k_{off}P + k_{PA}{A_0}^{\beta} & \frac{k_{on}P\psi}{2} \\ 0 & k_{PA}\beta {A_0}^{\beta-1}P_0 & \frac{k_{on}P\psi}{2} & \frac{k_{on}P\psi}{2} + k_{off}P + k_{PA}{A_0}^{\beta} \end{pmatrix} \tag{9}$$

where instability occurs when the eigenvalues of $J$ have at least one positive real part.

Finally, stably polarizable regions were numerically calculated by initializing the system with polarized distribution of aPARs and pPARs. Specifically, aPARs are initialized with two times the homogeneous steady state value at the anterior and zero at the posterior, and vice versa for pPARs. The simulation was considered capable of supporting polarity if $(A_a - P_a)$ and $(P_p - A_p)$ were both greater than 10%.

Importantly, we found the shape of the parameter space to be qualitatively similar to previous descriptions [1, 4], suggesting that the ODE simplification is reasonable.

#### 1.3 Calculating system dynamics in the phase portrait

Having confirmed that the simplified ODE system behaves qualitatively similar to the full PDE model, we sought to calculate how the system "moves" towards the steady state points, given a position in the phase portrait. These movements were represented by the quiver arrows in the phase portrait (Fig. 3). While each point in the phase portrait is degenerate, e.g. $(A_a - P_a) = 0.5$ can be satisfied with both $A_a = 2, P_a = 1.5$ or $A_a = 0.5, P_a = 0$, we can calculate how movement takes place near steady state, providing us with information on how the system moves away from steady state initial conditions during symmetry breaking. How the system is initialized across the phase portrait is formally written as:

where $x \in \{a, p\}$, and $A_0 = P_0$, which represent the values of homogeneous high aPAR and pPAR steady states respectively. This formulation effectively minimizes deviations from the midpoint of the homogeneous steady states. The conditions in the first and third rows ensure that the $A_x$ and $P_x$ values do not fall below zero.

To ensure that this initialization is comparable for different feedback strengths, which

| Initialization ($A_x - P_x$) | $A_x$ value | $P_x$ value |
|---|---|---|
| if $(A_x - P_x) > 2P_0$, | $(A_x - P_x)$ | 0 |
| if $2P_0 \geq (A_x - P_x) \geq -2A_0$, | $A_0 + \frac{1}{2}(A_x - P_x)$ | $P_0 - \frac{1}{2}(A_x - P_x)$ |
| if $-2A_0 > (A_x - P_x)$, | 0 | -$(A_x - P_x)$ |

would have different steady state values, we fixed $A_0$ and $P_0$ values for all simulations, based on simulations with $k_{AP} = k_{PA} = 0.1$. This tells us how the system would move towards the new steady state point(s) if the feedback strength suddenly changes, providing intuition on the effects of oscillatory feedback during PAR polarization.

Initialized in this way, we next approximated how the system behaves at each point of the phase portrait by numerically simulating equations (3) and (4) for 750 seconds. The movement of the system for each point can then be estimated by simply comparing the start and end points of the simulations, with the magnitude represented as the Euclidean distance. These give rise to the directionality and length of the arrows in the phase portrait.

#### 1.4 Estimating quasi-potential landscape

We estimated the quasi-potential landscape of the system by approximating equation (3) as stochastic differential equations (SDEs), using the Euler-Maruyama method. Formally:

$$\begin{aligned} U_x(t + \mathrm{d}t) &\approx U_x(t) + (\frac{\mathrm{d}U_x}{\mathrm{d}t})\mathrm{d}t + \sigma\sqrt{\mathrm{d}t}\xi \\ U_x &\in \{A_a, A_p, P_a, P_p\} \end{aligned} \tag{10}$$

where $\sigma$ define the noise term and $\xi$ is a vector of independent standard normal random variables (i.e. $\xi \sim \mathcal{N}(0, 1)$).

Specifically, we performed stochastic simulations $1 * 10^7$ times for 3,000s, with a $\sigma$ of 0.01, and recorded the final position of the simulation. The system is initialized as in section 1.3 at a random point across the phase portrait. We then reconstructed the quasi-potential landscape by first binning the final positions using a 2d histogram, and calculated quasi-potential ($Q$) as $Q = -\ln(\text{histogram density} + 1e - 9)$.

#### 1.5 Incorporating an aPAR-acting cue into the system

To incorporate a minimal aPAR-acting cue which mimics advective flows to the simplified reaction-diffusion model, we modified equation (3) as follows:

$$
\begin{aligned}
\frac{\mathrm{d}A_a}{\mathrm{d}t} &= \tilde{D}(A_a - A_p) + k_{on}A_{cyto} - k_{off}A_a - k_{AP}P_a^{\alpha}A_a + k_{cue}A_p \\
\frac{\mathrm{d}A_p}{\mathrm{d}t} &= \tilde{D}(A_p - A_a) + k_{on}A_{cyto} - k_{off}A_p - k_{AP}P_p^{\alpha}A_p - k_{cue}A_p \\
\frac{\mathrm{d}P_a}{\mathrm{d}t} &= \tilde{D}(P_a - P_p) + k_{on}P_{cyto} - k_{off}P_a - k_{PA}A_a^{\beta}P_a \\
\frac{\mathrm{d}P_p}{\mathrm{d}t} &= \tilde{D}(P_p - P_a) + k_{on}P_{cyto} - k_{off}P_p - k_{PA}A_p^{\beta}P_p
\end{aligned}
\tag{11}
$$

where $k_{cue}$ represents the strength of the cue. Note that the cue here does not act on pPARs, as symmetry breaking in *C. elegans* embryos is typically achieved by modulating the spatial activity or localization of aPARs [7, 1].

#### 1.6 Simulating PAR polarization using the simplified model

The full PAR polarization process shown in Fig. 3 is achieved by first initiating the system as in section 1.3 with $(A_a - P_a)$ and $(A_p - P_p)$ values obtained from calculating homogeneous and polarized states. Equation (11) was then simulated for 1250s with $k_{cue} = 0.002s^{-1}$ representing the polarity establishment phase, followed by simulation of equation (3) for 1250s, representing polarity maintenance phase.

We also simulated PAR polarization with dynamically changing cues and feedback. To ensure that the cues and feedback change smoothly, we modified $k_{cue}$, $k_{AP}$ and $k_{PA}$ to depend on time, i.e. $k_{cue}(t)$, $k_{AP}(t)$ and $k_{PA}(t)$, using a tanh function. These equations can be written as:

$$
\begin{aligned}
k_{cue}(t) &= k_{cue}\frac{1}{2}(\tanh(\frac{t - t_{on}}{w_1}) - \tanh(\frac{t - t_{off}}{w_2})) \\
k_{fb}(t) &= k_{fb}(1 - (1 - \frac{1}{F})\frac{1}{2}(\tanh(\frac{t - t_{dec}}{w_3}) - \tanh(\frac{t - t_{inc}}{w_4}))) \\
& fb \in \{PA, AP\}
\end{aligned}
\tag{12}
$$

Here, $t_{on}$ and $t_{off}$ define when the cue is turned on and off, $t_{dec}$ and $t_{inc}$ define when the feedback strength decreases and increases, $w_1, w_2, w_3, w_4$ defines the smoothness of the transition, and $F$ defines the fold change in amplitude during each oscillation cycle.

#### 1.7 Calculating transition points into the AP polarized basin

We wanted to calculate the transition point in the system toward the AP polarized basin ($A$ high at the anterior $P$ high at the posterior) during PAR polarization in the simplified model. This is complicated by the fact that our phase portrait projects a four-dimensional system, i.e. $A_a, A_p, P_a, P_p$ into two, i.e. $(A_a - P_a)$, $(A_p - P_p)$. To account for this, we generated different combinations of values for $A_a, A_p, P_a, P_p$ through a linear spacing of 50 points, ranging from 0 to $\psi\rho_U, U \in \{A, P\}$ and numerically solved for the steady state values. We then map the results and the $(A_a - P_a)$ and $(A_p - P_p)$ values back to the phase portrait. The transition point is defined by regions of space (approximated by a 2d histogram) where it is only possible to converge towards the AP polarized steady state.

#### 1.8 Comparing results with a tractable one species polarity model

We compared our topological analysis of the PAR model with a single-species polarity model to determine whether similar results can be reproduced in a mathematically tractable framework that does not require dimensionality reduction (Fig. S9). Following Nandan et al. [6], we adopted the minimal wave-pinning model of Mori et al. [8], in which positive feedback (representing Cdc42 auto-activation) drives polarization. The model makes three key assumptions: (i) the protein interconverts between an active, membrane-bound form and an inactive, cytoplasmic form; (ii) diffusion is fast in the cytoplasm and slow on the membrane; and (iii) the total amount of protein is conserved.

The system is described by:

$$\begin{aligned} \frac{\mathrm{d}X_a}{\mathrm{d}t} &= \tilde{D}\,(X_p - X_a) + k_{on}X_{cyto} - k_{off}X_a + \gamma\, X_{cyto}\frac{X_a^n}{K^n + X_a^n} \\ \frac{\mathrm{d}X_p}{\mathrm{d}t} &= \tilde{D}\,(X_a - X_p) + k_{on}X_{cyto} - k_{off}X_p + \gamma\, X_{cyto}\frac{X_p^n}{K^n + X_p^n}. \end{aligned} \tag{13}$$

The well-mixed cytoplasmic pool satisfies:

$$X_{cyto} = \rho_X - \psi\,\frac{X_a + X_p}{2} \tag{14}$$

Parameters can be found in Table S2.

A localized polarity cue is modelled as an additional on-rate acting only in the anterior compartment as before [8, 6],

$$\begin{aligned} \frac{\mathrm{d}X_a}{\mathrm{d}t} &= \tilde{D}\,(X_p - X_a) + k_{on}X_{cyto} - k_{off}X_a + \gamma\, X_{cyto}\frac{X_a^n}{K^n + X_a^n} + k_{cue}X_{cyto} \\ \frac{\mathrm{d}X_p}{\mathrm{d}t} &= \tilde{D}\,(X_a - X_p) + k_{on}X_{cyto} - k_{off}X_p + \gamma\, X_{cyto}\frac{X_p^n}{K^n + X_p^n}. \end{aligned} \tag{15}$$

Because the model is two-dimensional, the phase-space quiver can be obtained directly from Eqs. (13)–(15), without the need for time-averaging to average out curling in higher dimensions.

The quasi-potential landscape was estimated from the Fokker–Planck equation using a modified solver built by Holubec et al. [9],

$$\frac{\partial P(X_a, X_p, t)}{\partial t} = -\frac{\partial}{\partial X_a}[f_a\, P] - \frac{\partial}{\partial X_p}[f_p\, P] + D\left(\frac{\partial^2 P}{\partial X_a^2} + \frac{\partial^2 P}{\partial X_p^2}\right), \tag{16}$$

with $f_a = \frac{\mathrm{d}X_a}{\mathrm{d}t}$ and $f_p = \frac{\mathrm{d}X_p}{\mathrm{d}t}$ from Eqs. (13) or (15), and $D$ the (isotropic) noise strength, representing diffusing probabilities to allow stochastic state transitions. The quasi-potential is then $U = -\ln\big(P + 10^{-12}\big)$.

To locate the transition point, we simulated Eqs. (13)–(14) for $10^5$ seconds over a grid of initial conditions, is at high feedback ($\gamma$) levels. A trajectory was classified as polarized when $X_a > 1.1\, X_p$; the boundary of this basin of attraction defines the transition point.

#### 1.9 Simulating the full PDE model

We also corroborated our results using a full PDE model by simulating equations (1) and (2) with a custom built adaptive Runge-Kutta scheme in Python for calculating dynamics of the system [10], and using Euler's method with sufficiently small time steps if $k_{AP}$ , $k_{PA}$ and $k_{cue}$ are dynamic. Here, we represented $A_a, P_a, A_p, P_p$ as follows:

$$U_a = \int_0^{L/2} U(x)\mathrm{d}x,\ U_p = \int_{L/2}^{L} U(x)\mathrm{d}x,\ U \in \{A, P\} \tag{17}$$

To incorporate the cue, equation 1 is rewritten as follows:

$$\begin{aligned} \partial_t A &= D\partial_x^2 A + k_{on}A_{cyto} - k_{off}A - k_{AP}P^{\alpha}A + C_A(x) \\ \partial_t P &= D\partial_x^2 P + k_{on}P_{cyto} - k_{off}P - k_{PA}A^{\beta}P \\ C_A(x) &= \begin{cases} k_{cue}\int_{L/2}^{L} A(x)\mathrm{d}x, & \text{if } x < L/2 \\ -k_{cue}A(x), & \text{otherwise} \end{cases} \end{aligned} \tag{18}$$

where $C_A(x)$ is used to approximate the way cues act in the ODE form (equation (12)), and $k_{cue}$ simply defines the cue strength.

### 2 Constraining antagonism parameters to fit *C. elegans* embryos

Our simplified model suggests that oscillatory feedback enables temporal organization of either polarity-stable or cue-sensitive states, enabling polarization in various cellular contexts or with different cues. We next wanted to investigate whether this regulatory strategy could also be applied to early *C. elegans* embryos, using parameters specific to the system [1, 2]. Although most of the parameters have been previously determined, the antagonistic feedback parameters were arbitrarily chosen from regions within a parameter space that support both polarization and aPAR homogeneous states. Here, we took a fitting approach to better estimate the antagonism parameters [2].

#### 2.1 Estimating rate parameters for the antagonism terms

To estimate the antagonism rates, we fitted $k_{AP}$ and $k_{PA}$ from equation (1) with data from Nocodazole and Latrunculin A treated embryos (Fig. 2) to estimate rate parameters in a purely reaction-diffusion case, as both microtubules and actomyosin cortex have been shown to stabilize polarity [7, 11]. Specifically, we used a differential evolution approach to fit the data over the course of 12 minutes, minimizing the root mean square error of the sum of differences between the experiments and simulations over time (Fig. S11). The loss function L is formally written as:

$$\mathrm{L}(k_{convA}, k_{convP}, k_{PA}, k_{AP}) = \frac{1}{T}\int_0^T \left|\mathrm{experiment}(t) - \mathrm{simulation}\Big(t; k_{convA}, k_{convP}, k_{PA}, k_{AP}\Big)\right| \mathrm{d}t \tag{19}$$

where $k_{convA}$ and $k_{convP}$ represents conversion from fluorescence intensity obtained by experimental measurements to match the simulations, which have normalized $\rho_A$ and $\rho_P$ values [1].

To estimate how oscillating PAR-1 and CHIN-1 membrane levels affect $k_{AP}$, we simply assumed that $k_{AP}$ = membrane PAR-1 levels (Fig. S11).

#### 2.2 Examining polarity behaviors of the fitted antagonism parameters using a phase diagram

The parameter space defined by $k_{AP}$ and $k_{PA}$ for the *C. elegans* model is calculated similarly to the simplified ODE model (Fig. S11). However, the Jacobian used for calculating the instable states and how the polarity states differ.

Briefly, the Jacobian used for linear stability analysis is identical to Trong et al. [4], written as:

$$J = -\begin{pmatrix} D_A k^2 + k_{off,A} + \psi k_{on,A}\delta_{k,0} + k_{AP}P_o^{\alpha} & \alpha k_{AP}A_o P_o^{\alpha-1} \\ \beta k_{PA}P_o A_o^{\beta-1} & D_P k^2 + k_{off,P} + \psi k_{on,P}\delta_{k,0} + k_{PA}A_o^{\beta} \end{pmatrix} \tag{20}$$

where $\delta_{k,0}$ represents the Kronecker-delta term.

Next, polarity regions are defined when the following conditions are satisfied: $\int_0^{0.1L} A(x)\mathrm{d}x > 1.1 \int_0^{0.1L} P(x)\mathrm{d}x$ and $\int_{0.9L}^{L} P(x)\mathrm{d}x > 1.1 \int_{0.9L}^{L} A(x)\mathrm{d}x$.

### 3 Constructing representative P1 and P2 models

Our modeling results from the simplified ODE system suggest that in the absence of oscillatory feedback, PAR polarization becomes compromised when initiated from either homogeneous pPAR-high states or reversed polarity states. Notably, PAR polarization observed in the P blastomeres P1 and P2 provides clear examples of each case, respectively (Fig. 4 and 5). Therefore, we sought to represent polarization in these cell types using the fitted antagonism parameters and to test the role of oscillatory feedback.

#### 3.1 Developing an initial pPAR uniform model, roughly based on P1

PAR polarization in P1 involves the use of multiple polarity pathways (Fig. S12) [12, 13]. Among these are early cleavage furrow directed flows, which advects aPARs towards the nascent cell contact during cytokinesis, leading to a corresponding accumulation of aPARs. This serves as the first symmetry breaking cue, polarizing pPARs towards the embryo posterior. As the contact site occupies roughly 40% of the embryo perimeter, we represented this in the model by incorporating an aPAR pool at the embryo anterior. While it is unclear how enriched aPARs are at the anterior quantitatively, we found that a large range of values satisfy the early pPAR polarization away from the contact. We arbitrarily chose 30% of aPAR homogenous steady state values to represent this enrichment.

Following an early enrichment of aPARs at the contact site by furrow flows, P1 cells experience a second wave of advection, beginning roughly 6-7 minutes after cell birth. To represent this second wave of flows, we modified equation (1) as follows:

$$\begin{aligned} \partial_t A &= D\partial_x^2 A + k_{on}A_{cyto} - k_{off}A - k_{AP}P^{\alpha}A + \partial_x(\upsilon A) \\ \partial_t P &= D\partial_x^2 P + k_{on}P_{cyto} - k_{off}P - k_{PA}A^{\beta}P \end{aligned} \tag{21}$$

where $v$ represents cortical flow velocity, identical to previous work [14]:

$$v = \frac{60 - x}{74} e^{-\frac{(60-x)^2}{391}} - \frac{x}{1000} e^{-\frac{x^2}{100}} \tag{22}$$

Parameters can be found in Table S3.
Note that for simplicity, we considered these flows to act only on aPARs.

To incorporate oscillations into the model, we let $k_{AP}$ depend on time, based on the relative PAR-1 membrane levels, i.e. $k_{AP}(t)$ = PAR-1 mem($t$). We used PAR-1 membrane profiles from *par-3(-)* embryos, to isolate aPAR antagonism effects on PAR-1 membrane levels. Importantly, we rescaled the PAR-1 membrane levels over time to the cell cycle length of wildtype embryos as "P1" cells in *par-3(-)* embryos divide faster (Fig. S12).

We decided not to change the total dosage of the system ($\rho_A$ and $\rho_P$), cell size ($L$) and surface area to volume ratio ($\psi$) relative to the zygote, to minimize changes to the original model.

#### 3.2 Incorporation of WEE-1 perturbations

To represent WEE-1 inhibition in the model we shifted the PAR-1 membrane profile forwards in time to match experimental data, such that levels began to rise roughly 1.5 minutes after cell birth (Fig. S16).

#### 3.3 Incorporation of optogenetic knocksideways of PAR-6

To represent optogenetic knocksideways of PAR-6 in the model, we let the total aPAR pool depend on time, i.e $\rho_A(t)$ (Fig. S17). We noticed that release of PAR-6 from the mitochondria following blue light release was rapid and had detectable effects within one minute. Thus, we simplified the release kinetics in the model by assuming that PAR-6 at the mitochondria is removed instantly after alleviating blue-light induced sequestration. We arbitrarily assumed that blue light-induced sequestration of PAR-6 reduces $\rho_A$ by two thirds: the sequestration is unlikely to be complete, as we note that aPARs are still present on the membrane in the presence of blue light, but at significantly lower levels.

#### 3.4 Developing a polarity reversal model, roughly based on P2

PAR polarization in P2 involves at least two cues: the first cue polarizes the cell in the incorrect direction towards the embryo posterior, followed by a second cue that polarizes the cell towards the embryo anterior due to signals from the neighboring cell EMS (Fig. S18) [15, 12]. This polarity reversal phenotype was particularly obvious in the PAR-2 microtubule binding mutants. It is likely that cortical flows act to also reinforce the asymmetry of P2 cells as well, but these were neglected as these flows act much later and are not well characterized.

Due to the fast-acting nature of the first cue, we suspected that it is the same furrow cue found in P1 cells. We thus represented this cue in the same way as P1 cells.

The second cue enriches pPARs at the EMS-P2 cell contact. Although it is not clear how this enrichment is achieved, it is less consistent with the direct recruitment of pPARs to the contact, as this enrichment is lost when PKC-3 activity is disrupted [3]. Instead, the phenotype is most consistent with a local reduction in PKC-3 activity at the contact. Thus, we assumed that EMS signalling, which acts through MES-1/SRC-1, causes a local reduction in $k_{PA}$ at the cell contact, encompassing roughly 17.5% of the embryo. We incorporated this into the model by modifying equation (1) as follows:

$$\begin{aligned} \partial_t A &= D\partial_x^2 A + k_{on}A_{cyto} - k_{off}A - k_{AP}C_{EMScue}(x,t)P^{\alpha}A \\ \partial_t P &= D\partial_x^2 P + k_{on}P_{cyto} - k_{off}P - k_{PA}A^{\beta}P \end{aligned} \tag{23}$$

where $C_{EMScue}$ defines the signaling cue that acts to locally reduce aPAR activity at the contact, formally written as:

$$C_{EMScue}(x,t) = \begin{cases} \dfrac{1}{k_{EMScue}}, & \text{if } x \leq 0.175L \text{ and } t \in [t_{EMScueon}, t_{EMScueoff}] \\ 1, & \text{otherwise} \end{cases} \tag{24}$$

where $k_{EMScue}$ defines the reduction in antagonism strength at the contact, when the EMS signaling is active. We assume that the cue switches on 90 seconds after cell birth ($t_{EMScueon}$) and switches off 420 seconds after cell birth ($t_{EMScueoff}$) to match the observed PAR-2 distribution over the cell cycle. Notably, recent work has shown that SRC phosphorylation of PKC-3 reduces its membrane binding ability [16], which could provide a direct mechanism for reduction in this local antagonism.

As details of PAR polarization in P2 are poorly characterized, we used the same $k_{AP}$ oscillation profile and parameters as P1 as a proof of principle, including the effects of WEE-1 inhibition. We noted that the polarity reversal phenotypes in the simulations were subtle under these conditions, matching experiments (data not shown). However, polarity reversal was clear in *par-2(MT-)* mutants, which renders PAR-2 more sensitive to PKC-3 phosphorylation. We found that when we increase the antagonism of aPAR to pPAR ($k_{PA}$) to reflect this, arbitrarily by 20%, we are able to clearly capture the polarity reversal phenotype.
